# Resolving thyroid lineage cell trajectories merging into a dual endocrine gland in mammals

**DOI:** 10.64898/2026.03.11.710917

**Authors:** Macrina Lobo, Ellen Johansson, Sima Kumari, Elin Schoultz, Isak Ahlinder, Shawn Liang, Therese Carlsson, Bengt R Johansson, Pina Marotta, Mario De Felice, Jakob Dahlberg, Carolina Guibentif, Henrik Fagman, René Maehr, Mikael Nilsson

## Abstract

The thyroid has a remarkable evolution, first appearing in invertebrate chordates as an integral exocrine constituent of the pharyngeal endostyle that is transformed into an endocrine gland during metamorphosis in basal vertebrates. In mammals, the thyroid acquires a second endocrine cell type, calcitonin-producing C-cells, which for long were inferred a neural crest origin, shuttled to the embryonic thyroid by the ultimobranchial bodies. However, recent lineage tracing experiments firmly establish these neuroendocrine cells also derive from foregut endoderm. Key questions remaining unanswered are how thyroid primordia independently develop and, unlike in all non-mammalian vertebrates, merge into a dual endocrine organ. Here, by leveraging a single-cell transcriptome atlas derived from mouse pharyngeal endoderm and its subsequent cell fates, we characterize the global gene expression profile of thyroid- and ultimobranchial-derived progenitor cells and identify comprehensive gene regulatory networks of lineage-specific transcription factors and novel target genes predicted to differentially regulate cell proliferation, plasticity and differentiation during development. Spatiotemporal analyses reveal C-cell precursors are triggered to undergo epithelial-mesenchymal transition (EMT) and cell-autonomously down-regulate collagen IV and degrade laminin that delineates the ultimobranchial body epithelium. However, the EMT program is not fully deployed until both cell lineages are mixed and propagate conjointly thus forming the typical thyroid histoarchitecture of follicles and parafollicular C-cells, every follicle/C-cell unit being enveloped by a renewed basement membrane. Mixed-type thyroid carcinoma recapitulates a synchronous lineage growth pattern but only the neuroendocrine tumor cells are able to escape the compound follicle boundaries and become invasive adopting a C-cell precursor-like migratory phenotype.

## Introduction

The mammalian thyroid has a dual embryonic origin in the anterior foregut that in this context gives rise to two endocrine cell types, follicular epithelial cells (or thyrocytes) and neuroendocrine C-cells, producing thyroid hormone and calcitonin, respectively. Thyroid organogenesis implies that the midline thyroid primordium migrates downwards after having budded off sublingually from the pharyngeal floor, and then coalesce with the paired ultimobranchial bodies (Ubb), which derive from a more distal or posterior portion of the pharyngeal apparatus [1]. Progenitor cells of different origins thereafter mix and eventually differentiate, being closely associated in the thyroid tissue proper. This scenario contrasts to thyroid development in all other vertebrate species where the thyroid primordium and the Ubbs do not merge but establish separate endocrine glands that remain functionally independent throughout life [2]. The reason for thyroid evolving as a bifunctional gland consisting of different endocrine cell types in mammals – first recognized microscopically more than 100 years ago ([3] and refs therein) – is essentially unknown. Nonetheless, the dual tissue composition of the thyroid and its embryonic development is important to consider regarding tumor cell origin of different types of thyroid carcinomas, and for the putative influence of developmental traits might have on thyroid tumor evolution into distinct tumor phenotypes [3].

It was long assumed that thyroid C cells derive from a subpopulation of neural crest cells routed to the developing pharyngeal apparatus, and that this might explain their neuron-like nature [4]. However, from recent lineage tracing experiments in mice it is evident that calcitonin positive parafollicular cells originate directly from a division of pharyngeal endoderm that forms the fourth pharyngeal pouches [5]. Although Ubb and C cell development has been characterized in some detail [6], the precise morphogenetic mechanisms that govern disintegration of the Ubb in mammals followed by dissemination of differentiated C cells into the embryonic thyroid are poorly understood. It is evident that Ubbs do not develop in knockout mutants of transcription factors (TFs) with dedicated roles in early stages of pharyngeal development, such as Pax9 [7], Hoxa3 [8] and Tbx1 [9, 10], due to the posterior pouches being rudimentary or missing. Absence of thyroid C cells may also depend on failure of the prospective Ubb to separate from the pharyngeal endoderm or impaired thyroid-Ubb fusion as in *Nkx2-1* hemi- and homozygous knockout mouse embryos [11]. In humans, haploinsufficiency of NKX2-1 (also known as thyroid transcription factor-1 or TTF-1) variably leads to congenital hypothyroidism, respiratory distress and benign chorea, collectively named brain-lung-thyroid syndrome [12, 13].

Whether Ubb dysgenesis and shortage of calcitonin might contribute to disease development is not elucidated. Since both thyroid progenitor and Ubb/C-cell precursor cells express Nkx2-1, it can by hypothesized that yet unknown bidirectional signals, generated or modified by putative Nkx2-1 target genes, might be of morphogenetic importance to merging embryonic tissues into a composite gland with dual endocrine functions.

To get better insight into the molecular basis of mammalian thyroid development, we here characterize both thyroid and ultimobranchial cell lineages at single cell level by in-depth analysis of a recently published mouse pharyngeal endoderm-derived transcriptome atlas [14] and correlate differential gene expression patterns to morphogenesis spatiotemporally. This identifies new groups of TFs and target genes and predicts lineage-specific gene regulatory networks that promote not only thyroid and neuroendocrine differentiation but also regulate the merging of primordia and interactions of progenitor cells by which the unique histoarchitecture of the mammalian thyroid gland is established. In this process, C-cell precursors adopt a mixed epithelial-mesenchymal phenotype that likely facilitate thyroid colonization of C-cells, and which might be reactivated in invasive medullary thyroid carcinoma (MTC) cells.

## Results

### Single-cell transcriptome analysis recapitulates thyroid and Ubb/C-cell lineage development from different foregut endoderm origins

The mammalian thyroid develops from a median and two lateral anlagen in pharyngeal endoderm giving rise to the midline thyroid primordium and the paired Ubb that subsequently merge into a bilobed gland (Fig. 1a, b). At embryonic day (E or Eday) E15.5, as the lobes are being formed by branching morphogenesis in mice [15], it is possible to spatially distinguish cells of different embryonic origin by their specific gene expression patterns (Fig. 1c-f). Hence, whereas Nkx2-1 is expressed in both lineages (Fig. 1c), Pax8 and Foxa2 are exclusively expressed in thyroid or Ubb cells, respectively (Fig. 1d, e; [5, 16]). Moreover, C cells entering the embryonic thyroid can be distinguished due to the expression of calcitonin from residual Ubb cells that are not yet differentiated (Fig. 1e). To elucidate if global analysis of TF and target gene expression across Eday enables tracing of lineage cell fates committed in earlier developmental stages, we employed a single cell transcriptome database primarily established from pharyngeal endoderm encompassing E9.5-E12.5 [14] to map their developmental trajectories foregoing thyroid-Ubb fusion. Previous analyses [14] readily captured cluster- and organ-specific molecular signatures of pharyngal endoderm derivatives (Supplementary Fig. 1a). We pursued subset analysis of UMAP clusters 12, 13, 17 and 22 amounting to 5,904 cells or 10.9% of the total atlas cell number which, as identified by expression of signature genes (Supplementary Fig. 1b), comprise most, if not all, thyroid and Ubb lineage cells traced by this procedure (Fig. 1g, h).

**Fig. 1.**
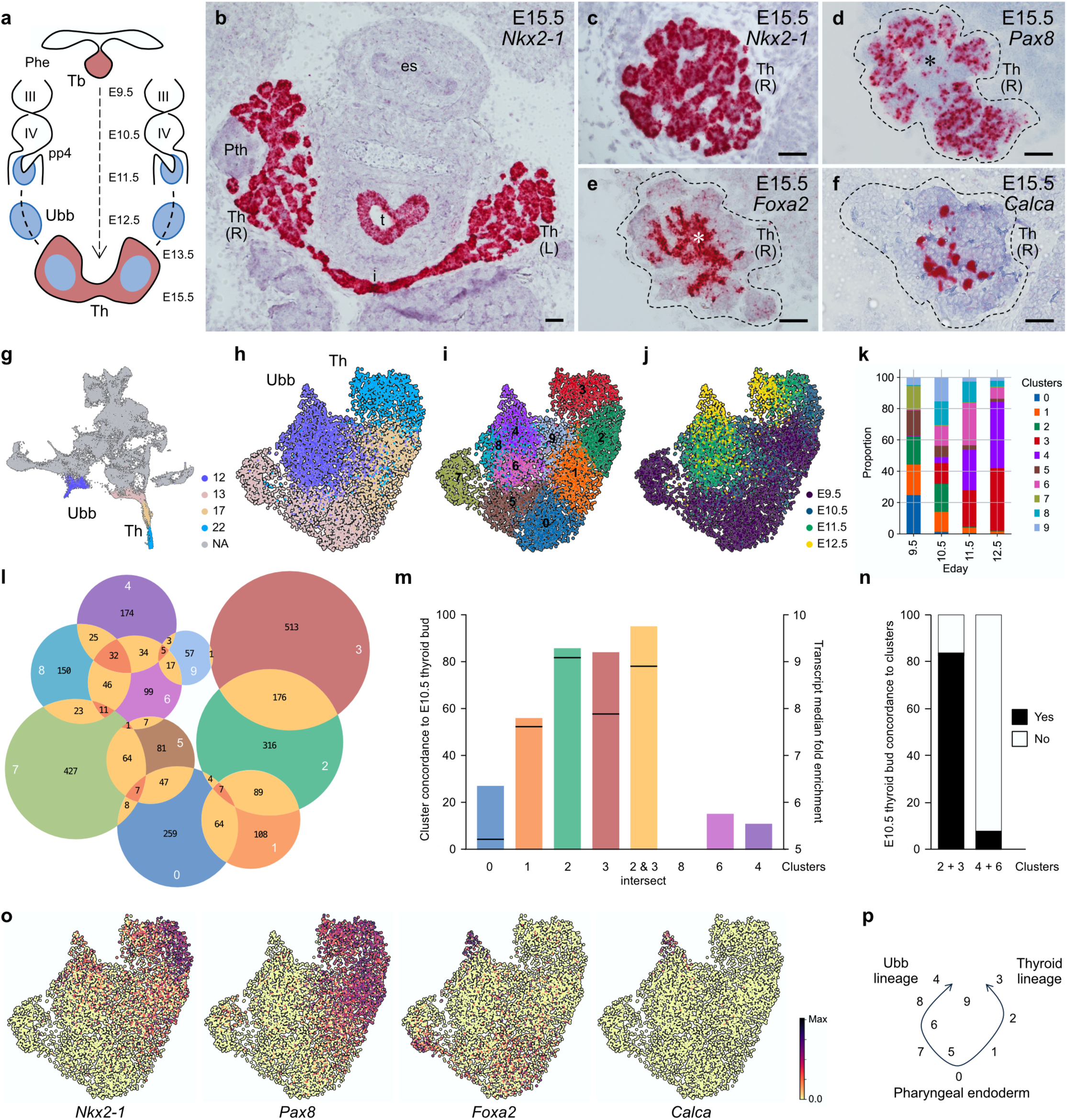
Single-cell transcriptome mapping of mouse thyroid and ultimobranchial body development. **a** Routes of thyroid organogenesis from median and lateral anlagen depicted across embryonic (E) day (Eday) in mice. III and IV indicates branchial arch numbers. **b-f**. RNAscope imaging of lineage-specific expression of *Nkx2-1* (**b, c**), *Pax8* (**d**), *Foxa2* (**e**) and *Calca* (encoding calcitonin) (**f**) at E15.5 i.e. after thyroid primordium and Ubb merged. Scale bars: 100 µm. **g-j** UMAP visualization obtained from a pharyngeal endoderm single-cell transcriptomic time-course dataset (n = 54,044 cells) captured daily between E9.5 and E12.5 (from [14]) with louvain clusters of thyroid (Th), Ubb and corresponding progenitor populations (clusters 12, 13, 17 and 22) highlighted (**g**) re-analyzed separately and colored by original clusters in g (**h**) colored by Leiden clusters 0-9 (**i**) and colored by Eday (**j**). Each dot represents a cell. **k** Stacked bar plot showing the proportion of cells in every cluster identified at each Eday with clusters belonging to the Ubb and thyroid labeled. **l** Venn diagrams of number of enriched genes per cluster and intersects. Diagram colors correspond to cluster numbers 0-9 as indicated. **m, n** Comparison of genes enriched in single-cell clusters of the predicted thyroid and Ubb lineage identities with transcripts previously found to be enriched in the E10.5 thyroid bud [17]. Fraction of top 50-ranked cluster genes concordant with genes identified *in vivo* (**m**) and fraction of top 50-ranked bud genes recovered in clusters (**n**). Bar-transversing lines in (m) indicate mean fold enrichment of bud transcripts for each cluster comparison. **o** UMAP embedding of selected marker genes at single-cell resolution with the cells colored by the log2-normalized expression. UMAPs of selected genes overlaid original clusters and across Eday, respectively, are shown in Supplementary Figs. 1 and 2. **p** Lineage progression suggested from cluster intersect distribution (n) and confirmed by cluster current analysis (see Fig. 2). Phe pharyngeal endoderm, Tb thyroid bud, Th, thyroid (primordium or lobe), L left (lobe), R right (lobe), Pth parathyroid, es esophagus, t trachea.

Unbiased re-clustering of these endodermal subpopulations delineates ten clusters numbered 0-9 that are distinguished by cluster-specific gene expression signatures (Fig. 1i, Supplementary Fig. 2). Notably, these clusters harmonize with progression across Eday (Fig. 1j), which conceivably identify clusters 3 and 4 as the most differentiated lineage cell stages (Fig. 1k).

Next, cluster-wise comparison were conducted to identify unique versus shared gene expression profiles, which confirmed cluster-specific expression signatures (Fig. 1l). Although a significant number of transcripts show a considerable overlap there is a clear separation of clusters 1-3 on one side and clusters 4-9 on the other side whereas cluster 0 intersects with both lineages. To further corroborate lineage identity of clusters, we compared single-cell global gene enrichment with published embryonic thyroid transcriptome datasets obtained from microarray analysis of laser-captured mouse thyroid and lung buds at E10.5 [17]. This shows that nearly 85% of the 50 top-ranked genes and 95% of the overlapping genes in clusters 2 and 3 are concordant with transcripts previously found to be enriched (>3-fold, cutoff level) in the thyroid bud (Fig. 1m) [17]. Lower concordance is evident for enriched genes in cluster 1 (56%) and cluster 0 (27%), suggesting these clusters consist of immature cells of the thyroid lineage. By contrast, enriched genes of the presumed Ubb clusters 4 and 6 show low concordance (11% and 15%, respectively) to thyroid bud transcripts (Fig. 1m). Moreover, a reverse comparison reveals that 85% of previously identified 50 top-ranked E10.5 bud transcripts are collectively recovered in clusters 2+3 whereas only 8% are present in clusters 4+6 (Fig. 1n).

UMAP embedding of individual TF transcripts confirmed lineage identity of annotated clusters (Fig. 1o, Supplementary Fig. 3). Most cells in clusters 1-3 co-express *Nkx2-1* and *Pax8* consistent with being identical to thyroid progenitor cells that eventually will develop into follicles (Fig. 1c, d). Accordingly, Pax8 shows a broader cluster distribution than Nkx2-1 reminiscent of their sequential induction *in vivo*, Pax8 being expressed earlier and more widespread in foregut endoderm whereas Nkx2-1 expression is confined to a limited number of cells forming the thyroid placode [18]. By contrast, clusters 4, 6 and 8 comprise cells that variably co-express *Foxa2*, *Meox1* and *Ripply3* along with *Nkx2-1* and, in cluster 4 only, *Calca* encoding calcitonin (Fig. 1o; Supplementary Fig. 3). These are known or suggested TFs of the fourth pharyngeal pouch and the Ubb lineage [5, 11, 14] which, taking all clusters together, display distinct expression patterns over time: Ripply3 transiently, Meox1 evenly across Eday, and Foxa2 being expressed both early and late seemingly re-induced along with calcitonin. Accumulation of Ripply3^+^ cells suggests cluster 7 likely represents caudal pharyngeal endoderm [19], which therefore might comprise putative Ubb progenitors. Interestingly, upregulation of Ripply3 restricted to cluster 7 (z-score: 33.9), cluster 8 (15.1) and cluster 9 (7.1) mimics that of Hoxb1 (z-score in cluster 7: 36.6; in cluster 8: 18.4; cluster 9: 7.6), the expression of which is restricted to the most inferior pharyngeal pouch during pharyngeal segmentation [20]. Ripply3^+^/Hoxb1^+^ cells in clusters 8 and 9 might thus potentially retain immature Ubb features predominating in cluster 7. Meox1, a mesenchyme homeobox gene, is transcriptionally expressed in the Ubb *in vivo* [14] but its role in Ubb development is yet unknown.

Altogether, these observations clearly indicate that re-clustering faithfully reproduced the originally identified thyroid and Ubb lineage transcriptomes confined to clusters 1-3 (thyroid) and clusters 4, 6 and 8 (Ubb), and that clusters 0, 5 and 7 likely represent pharyngeal endoderm in the computed model (Fig. 1p).

### Pseudotime analysis and knockout simulations distinguish cell fate probabilities of acquiring thyroid and Ubb/C-cell terminal states

Using Harmony [21] to join adjacent time points, we performed pseudotime analysis with Palantir [22] and visualized trajectories on a force directed embedding (Fig. 2a). Next, we applied CellRank [23] for cell fate mapping on the pseudotime derived transition matrix and inferred streamlines confirming predicted cluster currents (Fig. 2b), allowing us to reconstruct fate probabilities (Fig. 2c) and to identify lineage drivers (Supplementary Dataset 1 and 2). In this model, cluster 0 consists of cells that share features of an immature endoderm stage from which both lineages originate, whereas clusters 5 and 7 represent an intermediate stage corresponding to the fourth pharyngeal pouch from which Ubb develops (Fig. 2c). Complementarily, we employed an optimal transport (OT) based method (moscot) [24, 25] to map cells across time points and automatically pinpoint terminal states for both lineages. By modelling differentiation as a stochastic process from E9.5 to E12.5 it is thus feasible to automatically infer thyroid and Ubb terminal clusters (Fig. 2d).

**Fig. 2.**
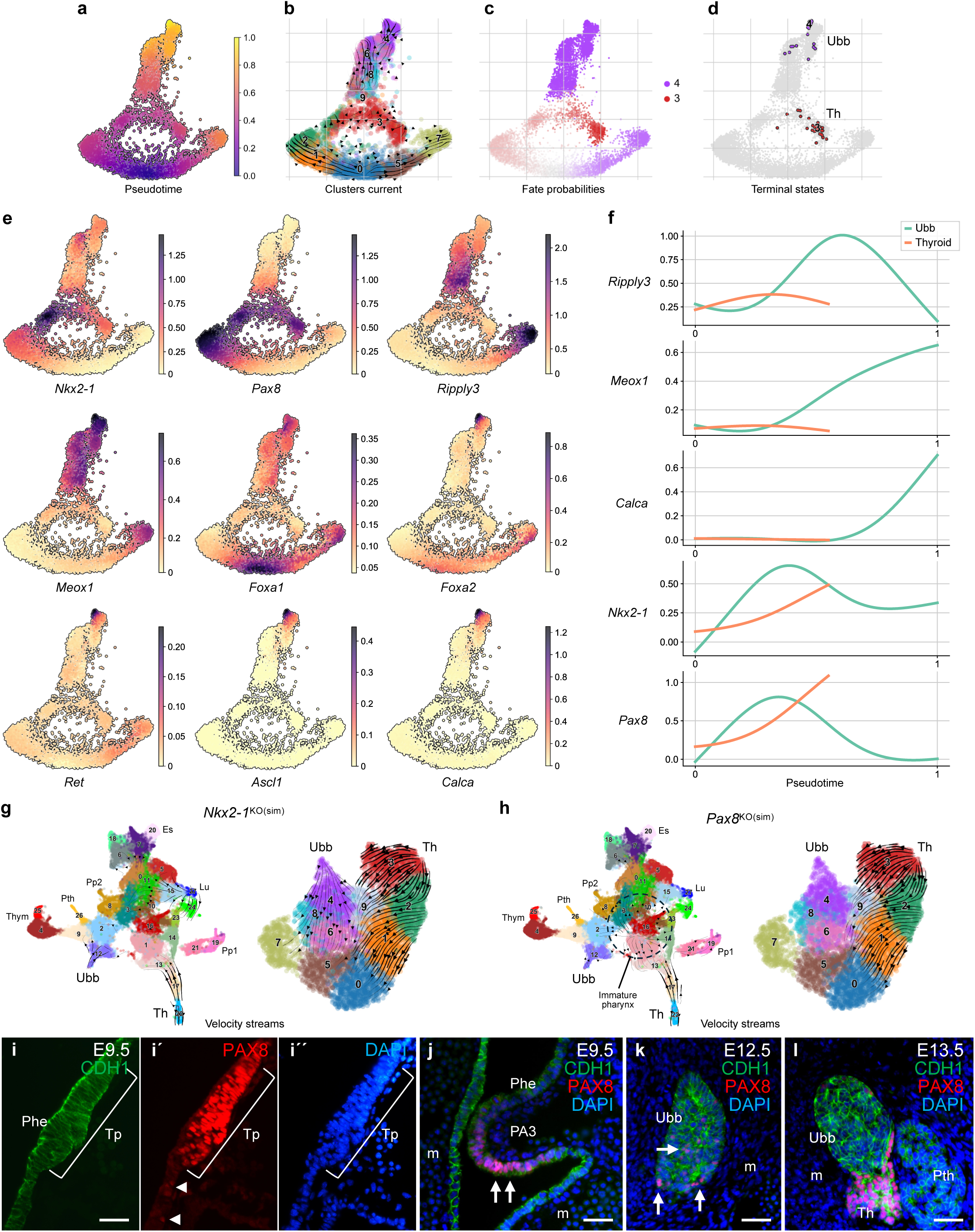
Trajectory interference and validation of thyroid and Ubb/C-cell lineage fates. **a-d** Force directed embedding of the Ubb and thyroid lineage cells from Fig. 1 obtained via integrating successive timepoints with Harmony [21]. **a** Cells are colored by inferred pseudotime using Palantir [22]. **b** Cell streamlines inferred using CellRank [23] from the transition matrix obtained via pseudotime analysis in (a) with cells are colored by the clusters from Fig. 1. **c** Cells are colored by the probability of maturing to the Ubb (purple) or thyroid (red) fate determined according to the transition matrix from (a). **d** Force directed embedding with all cells shown in gray except the cells most confidently assigned to each of the two predicted terminal states (Ubb and thyroid). The transition matrix was inferred using a complementary optimal transport (OT) based method (moscot) to map cells across time points [24, 25]. **e** Force embeddings colored by the MAGIC [139] imputed gene expression of key marker genes. **f** Gene trends showing the imputed expression of key markers along pseudotime (from (a)) for the UBB (green) and thyroid (red) lineages. **g, h** Knockout simulations (ko(sim)) of *Nkx2-1* (**g**) and *Pax8* (**h**) based on multiomic GRN inference with CellOracle and predicted cell-state changes. Visualizations overlaid on the scRNA pharynx atlas according to (left panels) and clusters 0-9 of this study (right panels). Changes in gene expression between the simulated knockouts and atlas cells are displayed as velocity streams on the UMAP embedding using scVelo [30]. **i-l** Immunofluorescent detection of PAX8+ cells in pharyngeal endoderm and derivatives across Eday; epithelial and mesenchymal tissues distinguished by CDH1/E-cadherin and DAPI nuclear stain. **i** Thyroid placode at E9.5; single channels shown for clarity (in i-i’’). **j** Inferior pouch endoderm at E9.5. **k** Ubb at E12.5. **l** Ubb and thyroid merging at E13.5. Arrowheads (in i’) and arrows (in j, k) indicate Pax8^+^ cells. Phe pharyngeal endoderm, Tp thyroid placode, PA3 third pharyngeal arch, Ubb ultimobranchial body, Th thyroid, Pth parathyroid, m mesenchyme. Scale bars: 50 μm.

Next, we visualized imputed expression profiles of key markers featuring pharyngeal endoderm by Nkx2-1^-^/Pax8^low^/Ripply3^+^/Meox1^+^/Foxa1^+^/Foxa2^+^ cells in clusters 0, 5 and 7, thyroid lineage by Nkx2-1^+^/Pax8^+^/Ripply3^low^/Meox1^low^/Foxa1^low^/Foxa2^-^ cells in clusters 1-3, and Ubb lineage by Nkx2-1^+^/Pax8^-^/Ripply3^+^/Meox1^+^/Foxa1^+^/Foxa2^+^ cells in clusters 4, 6 and 8 (Fig. 2f). Calcitonin^+^ cells co-expressing Ascl1 (also known as Mash1, a TF required for C-cell survival colonizing the embryonic thyroid [26, 27]) and the protooncogene Ret (frequently mutated in malignant C-cell derived tumors [28]) confirm cluster 4 as the most differentiated stage (Fig. 2e). Different expression kinetics for each of these genes is evident from the imputed expression along pseudotime (Fig. 2f).

Dual lineage expression of Nkx2-1 is expected but transient expression of Pax8 in the Ubb lineage is surprising (Fig. 2e, f). However, the gene regulatory network (GRN) inferred from scRNAseq and scATACseq data in Magaletta et al [14] using CellOracle [29] supports that Pax8 regulates thyroid lineage genes only whereas similar *in silico* analyses identify Nkx2-1 targets in both thyroid- and Ubb-GRNs. Using the GRN, we simulated perturbation of both TF genes and visualized the perturbation kinetics using scVelo-based stream plots [30] projected onto the scRNA atlas of pharyngeal organ development [14] and the present UMAP embedding of clusters 0-9 (Fig. 2g, h). As expected from expression patterns and functions *in vivo* [11], *Nkx2-1* knockout simulations reveal cluster-specific velocity streams toward clusters representing earlier developmental stages for cells with thyroid, Ubb and lung signatures (Fig. 2g). Of these lineages, only thyroid cells are influenced by perturbation of *Pax8* (Fig. 2h). However, widespread streams from centrally distributed immature cells towards several clusters with other tissue identities (Fig. 2h, left panel) suggest that Pax8 additionally might be expressed in endodermal cells not committed to a thyroid fate. Indeed, besides the Pax8 positive thyroid placode (Fig. 2i-i’’) we identified Pax8^+^ cells in a second location initially confined to the inferior-most pharyngeal pouch and later transiently in the Ubb (Fig. 2j-l). Since *Pax8* null mice has no obvious Ubb phenotype [16], a putative role of Pax8 in pharyngeal development apart from the midline thyroid might be redundant or in cooperation with Pax2 [31–33].

### Identification of lineage drivers and gene regulatory networks associated with embryonic thyroid differentiation

To further determine lineage specificity annotated clusters were investigated for enriched pathways. Upregulation of genes involved in thyroid hormone (TH) production and function is evident in cluster 3 and to a lesser extent in cluster 2 (Fig. 3a, b). Enriched thyroid genes comprise factors of critical importance for thyroid differentiation (*Pax8*, *Foxe1*), thyroid regulation (*Tshr*), and TH biosynthesis (*Tg*, *Duox2, Duoxa2, Iyd*) (Fig. 3c). Although the number of differentiating cells are not in majority and no transcripts of genes required for iodide uptake (*Slc5a5* encoding sodium-iodide symporter) and iodination (*Tpo* encoding thyroid peroxidase) are recovered by scRNA-seq, these results confirm recent observations that embryonic thyroid differentiation is initiated earlier than previously understood [34].

**Fig. 3.**
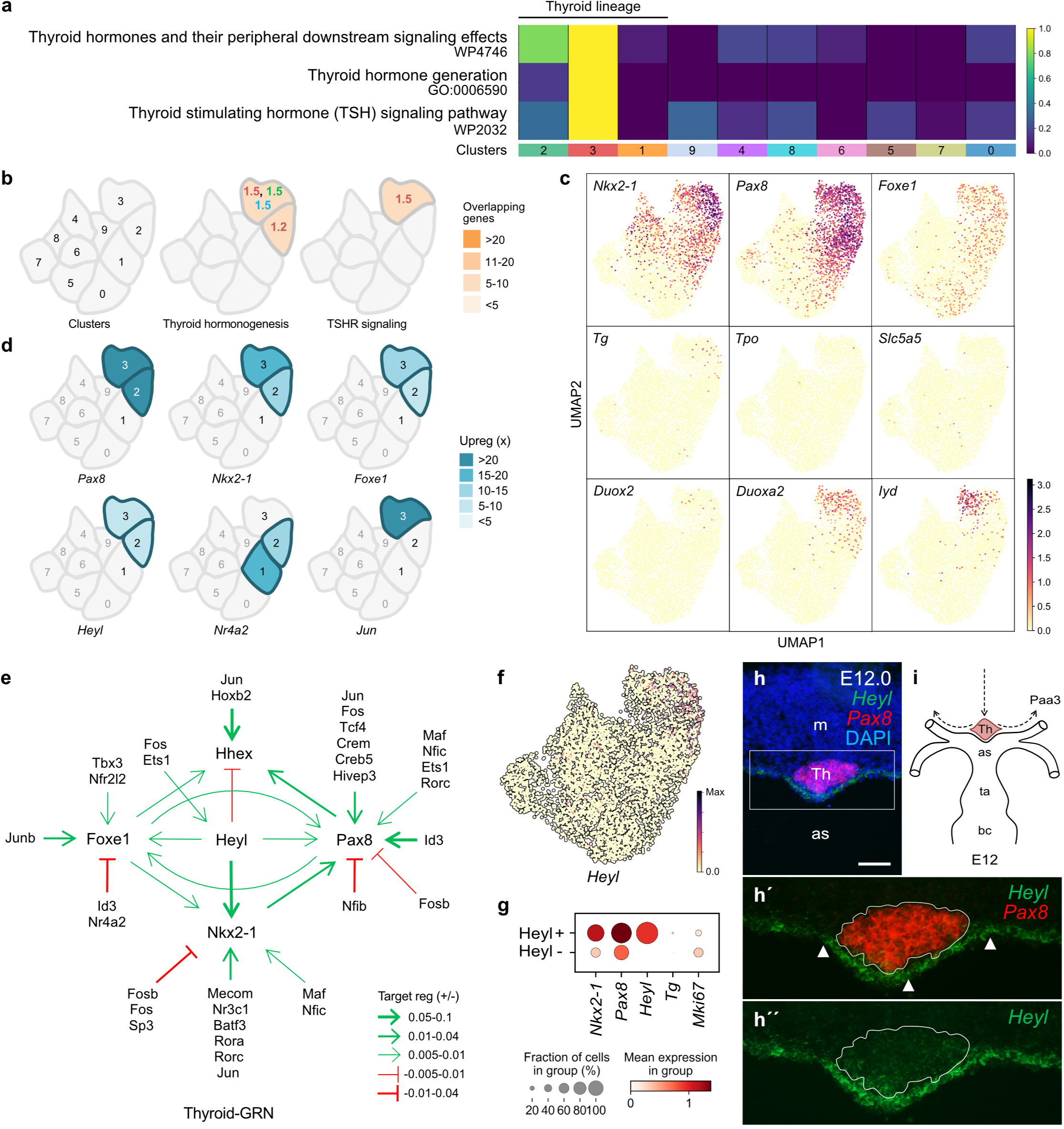
Gene enrichment in thyroid lineage cells and the predicted gene regulatory network of thyroid differentiation. **a** Heatmap of a curated list of pathways associated with the differentiated thyroid cell state. Heat color intensity indicates the row normalized −log10 adjusted p-values (Benjamini–Hochberg) obtained from an enrichment analysis (with EnrichR) of the genes differentially upregulated (adjusted p-value < 0.01, log2-fold change > 1.0) in each cluster over a background comprising the remaining clusters. Clusters are indicated by numbers and colors according to Fig. 1i. **b** Cluster-specific enrichment of thyroid pathway-associated genes with −log10 adjusted p-values indicated curated from GO Biological Process (green), KEGG (blue) and WikiPathway (red). Color intensity corresponds to number of genes overlapping between the list of genes enriched in the pathway(s) and the upregulated genes (adjusted p-value <0.01, log2-fold change >1.0) in the corresponding cluster (numbers 0-9 overlaid ghost UMAP to the left). **c** UMAP visualizations of thyroid-specific gene expression implicated in thyroid hormone synthesis and regulation. **d** Selection of genes encoding transcription factors (TFs) primarily identified in the predicted thyroid gene regulatory network (Thyroid-GRN; see below) and being exclusively upregulated in thyroid lineage clusters 1-3 (Table 1a). Curated from lists comprising all enriched genes per cluster. **e** Schematic GRN in thyroid lineage cells predicted by CellOracle network inference from multiomic profiles. Sharp (green) or blunt (red) arrows indicate source-target direction and arrow thickness represents positive or negative average cluster-specific GRN TF-target interaction scores of up- and downregulated genes. TFs not directly targeting one or more of the key thyroid TFs Nkx2-1, Pax8, Foxe1 and Hhex are not included. **f** UMAP visualization of *Heyl*-expressing cells accumulated in clusters with thyroid identity. **g** Dot plots of *Heyl*^+^ and *Heyl* negative cells co-expressing *Nkx2-1*, *Pax8* and *Tg*. Dot size indicates the fraction of cells in thyroid clusters expressing the gene, and the color indicates the mean log2-normalized expression of the gene. **h** *Heyl* expression in *Pax8*^+^ thyroid cells at E12.0 identified by RNAscope Multiplex fluorescence; **h’-h’’** show high power of boxed area in (h) without dapi (h’) and green channel only (h’’), with thyroid tissue outlined, for clarity. Arrowheads indicate arterial wall. **i** Close association of the thyroid primordium with central embryonic vessels emerging from the cardiac outflow tract; arrows indicate prior and subsequent thyroid migration pathways [52]. Th thyroid primordium (midline location), Paa pharyngeal arch artery, as aortic sac, ta truncus arteriosus, bc bulbus cordis. Scale bar: 100 µm.

**Table 1.** Cluster-specific enrichment of transcription factors identified by CellOracle in the predicted gene regulatory subnetworks of thyroid and ultimobranchial lineage cells.

| A. Thyroid-GRN |  |  |  | B. Ubb-GRN |  |  |  |
| --- | --- | --- | --- | --- | --- | --- | --- |
| TF | Cluster 1 | Cluster 2 | Cluster 3 | TF | Cluster 6 | Cluster 8 | Cluster 4 |
| *Arntl* | - | - | 8.2 (2.32) | *Bach2* | - | - | 9.9 (1.47) |
| *Bach1* | - | - | - | *Etv1* | 7.2 (1.71) | 6.8 (1.82) | - |
| *Batf3* | - | 5.1 (1.22) | 5.5 (1.29) | *Foxp2* | - | - | - |
| *Bcl11b* | - | - | 15.7 (1.87) | *Hoxa2* | 8.5 (1.56) | - | - |
| *Creb5* | - | 5.6 (1.34) | 5.8 (1.33) | *Hoxa5* | 5.6 (1.25) | 10.4 (2.41) | 10.6 (1.89) |
| *Crem* | - | 10.2 (1.34) | - | *Hoxb1* | - | 18.4 (2.27) | - |
| *Cux1* | - | - | - | *Hoxb2* | - | 10.1 (1.00) | - |
| *Egr1* | - | - | 13.7 (2.38) | *Hoxb4* | 9.2 (1.41) | 10.9 (2.04) | 16.0 (2.23) |
| *Elf3* | - | 8.3 (1.38) | - | *Hoxb5* | 8.1 (1.26) | 13.9 (2.53) | 7.9 (1.16) |
| *Elk3* | - | - | 6.8 (1.40) | *Hoxb6* | - | 13.7 (2.93) | 13.3 (2.16) |
| *Ets1* | - | - | 6.6 (1.83) | *Hoxc5* | 8.6 (1.30) | 14.6 (2.43) | 24.0 (2.98) |
| *Fos* | - | - | 21.9 (2.96) | *Id2* | - | - | - |
| *Fosb* | - | - | 11.3 (2.91) | *Id3* | - | - | - |
| *Foxe1* | - | 6.9 (1.26) | 10.4 (2.02) | *Irx2* | - | 4.3 (1.22) | - |
| *Foxp2* | - | - | - | *Meis1* | 12.9 (1.27) | - | 12.6 (1.15) |
| *Heyl* | - | 5.9 (2.09) | 7.5 (2.70) | *Meis2* | 27.8 (1.36) | - | - |
| *Hivep3* | - | 8.6 (1.73) | 17.2 (3.41) | *Meis3* | - | - | - |
| *Hoxb2* | - | - | - | *Meox1* | 6.8 (1.00) | 10.0 (1.64) | 14.2 (2.02) |
| *Id1* | - | - | - | *Nfia* | - | 3.3 (1.26) | 10.9 (2.44) |
| *Id3* | - | - | - | *Nfib* | - | - | 20.7 (2.16) |
| *Isl1* | - | - | - | *Nkx2-1* | - | - | - |
| *Jun* | - | - | 25.5 (1.93) | *Nkx2-5* | - | - | - |
| *Junb* | - | - | 14.6 (2.30) | *Pax9* | 20.7 (1.79) | 15.5 (1.56) | 23.1 (1.73) |
| *Klf6* | 10.7 (1.01) | 14.0 (1.45) | - | *Pbx1* | - | - | 26.5 (1.27) |
| *Lef1* | - | - | - | *Prdm1* | - | 7.5 (1.44) | - |
| *Maf* | - | - | 12.3 (2.27) | *Sox4* | - | - | - |
| *Mecom* | - | - | 10.7 (1.60) | *Ybx1* | - | - | - |
| *Nfat5* | - | - | - | *Zfhx3* | 13.0 (1.09) | - | 22.7 (1.60) |
| *Nfe2l2* | - | 8.8 (1.01) | - |  |  |  |  |
| *Nfib* | - | - | - |  |  |  |  |
| *Nfic* | - | - | 7.8 (1.91) |  |  |  |  |
| *Nkx2-1* | - | 14.2 (1.75) | 19.6 (2.35) |  |  |  |  |
| *Nr3c1* | - | - | 12.8 (1.79) |  |  |  |  |
| *Nr4a2* | 15.1 (2.07) | 11.7 (1.63) | - |  |  |  |  |
| *Pax8* | - | 35.2 (2.70) | 33.3 (2.55) |  |  |  |  |
| *Rora* | - | - | - |  |  |  |  |
| *Rorc* | - | - | 6.5 (1.97) |  |  |  |  |
| *Sox9* | - | - | 14.2 (1.88) |  |  |  |  |
| *Sp3* | - | - | - |  |  |  |  |
| *Tbx3* | - | 19.7 (1.66) | - |  |  |  |  |
| *Tcf4* | - | - | 19.7 (1.40) |  |  |  |  |

TF transcription factor (alphabetically listed), GRN gene regulatory network
Cluster scores: times upregulated (logfoldchange)

Putative TF-gene associations were characterized to get insight into regulatory mechanisms that might govern *de novo* functional differentiation of the embryonic thyroid, as yet an unsolved issue. After filtering (see Methods), the number of TFs identified by GRN modeling is limited to 41 for the thyroid lineage. Cluster analysis predicted that 23 thyroid TFs including the classical quartet Hhex, Pax8, Foxe1 and Nkx2-1 [35] are enriched progressively along with thyroid differentiation i.e. in the cluster order 1→ 2→ 3 (Table 1, left panel; Fig. 3d). Many of these upregulated TFs are predicted key lineage drivers (Supplementary Dataset 1), the ranking list being additionally validated by concordance with strongly enriched genes e.g. *Prlr* and *Bcl2* in early thyroid development *in vivo* [17]. Among the top-ranked TFs are (rank order): Hhex (8), Fos (11), Pax8 (19), Jun (24), Hivep3 (27), Egr1 (35), Junb (36), Fosb (40), Bcl11b (48), Nr3c1 (68) and Tcf4 (69).

Nkx2-1 takes a lower rank (139) among thyroid lineage drivers, seemingly not conforming with the fact that both Nkx2-1 and Pax8 are required and sufficient to generate functional thyroid cells from embryonic stem cells [36]. A fairly low number of predicted Nkx2-1 target genes (n=51; Supplementary Table 1) as compared to >900 targets for Pax8 (not listed) is remarkable, arguing that Nkx2-1 might regulate transcription in embryonic thyroid cells by transactivating or altering the expression of other TFs. Strong lineage drivers among Nkx2-1 target genes are Pax8 and Tcf4, the expression of which is also increased in cluster 3 (Table 1, left panel). In the predicted thyroid-GRN, Nkx2-1 regulates Pax8 but none of the other classical thyroid TFs (Fig. 3e). This contrasts to thyroid progenitors at E10.5 in which Pax8 expression is independent of Nkx2-1 [37], suggesting that transcriptional regulation of Pax8 expression evolves differently beyond the bud stage.

We investigated this possibility in heterozygous *Nkx2.1-CreERT2* mice in which one *Nkx2-1* allele is disrupted and hence equivalent to *Nkx2-1^+/-^* mice [38]. This showed that monoallelic deletion of *Nkx2-1* downregulates *Pax8* along with decreased mRNA expression levels of Nkx2-1 in adult thyroid follicular cells (Supplementary Fig. 4a-a’’’, 4b-b’’’). Similarly, the expression of *Tcf4* diminishes in *Nkx2.1-CreERT2^+/-^* mutants (Supplementary Fig. 4c-f). Flattened follicular epithelia and altered lumen size indicate that cells deficient of one *Nkx2-*1 allele are functionally affected (Supplementary Fig. 4a-f), i.e. corresponding to thyroid abnormalities previously reported for heterozygous *Nkx2-1* null mice [39]. Altogether, this agrees with previous findings that Nkx2-1 gene dosage matters for both organogenesis and maintenance of the thyroid phenotype [11, 39], and suggests that Pax8 and potentially also Tcf4 might contribute being upregulated by Nkx2-1.

It is important to note that the thyroid-GRN as outlined in Fig. 3e is a simplified model limited to TFs putatively regulating directly or indirectly one or several of the classical four TFs. For example, Egr1 with most GRN interactions of all thyroid TFs (>1,000 predicted target genes) is strongly upregulated in cluster 3 (Table 1a) but excluded from the subnetwork presumed to target Hhex, Pax8, Foxe1 and Nkx2-1. Nonetheless, its reliability is further evidenced by mapping the transcriptional subnetwork of *Cdh4* (encoding R-cadherin) and *Cdh16* (encoding Ksp-cadherin) implicated in embryonic thyroid differentiation [40, 41]. Both cadherins are exclusively enriched in cluster 3 (Supplementary Fig. 5a) and ranked 210 for Cdh16 and 238 for Cdh4 among thyroid lineage drivers (Supplementary Dataset 1). Their predicted regulatory network involves Pax8 and Foxe1 and several other TFs that are also enriched in cluster 3 (Supplementary Fig. 5b, c). It is previously known that Cdh16 is a Pax8 regulated gene that govern the acquisition of apical polarity [42, 43], which during development correlates with onset of folliculogenesis [34]. Increased transcriptional activity of Pax8 downstream of Nkx2-1 thus offers a plausible mechanism by which follicle formation and the differentiated thyroid phenotype involving Cdh16 are promoted by Nkx2-1 [44]. Similarly, being targeted exclusively by Pax8 in the developing thyroid [37], Foxe1 might mediate Nkx2-1 driven regulation of Cdh4 downstream of Pax8, as suggested from the GRN model (Supplementary Fig. 5b).

### Heyl is part of the thyroid-GRN and dynamically expressed during thyroid development

To further validate the predicted thyroid-GRN, we focused our interest on Heyl, a dynamic Notch signaling effector in development [45]. Heyl is moderately enriched in clusters 2 and 3 (Fig. 3d), constitutes the single most important regulator of Nkx2-1 (Fig. 3e), and encounters target gene associations in the same order of magnitude as Pax8. From the UMAP and dot plot it is evident that Heyl-expressing cells comprise a subpopulation of yet undifferentiated Nkx2-1^+^/Pax8^+^ progenitors (Fig. 3f, g). Moreover, as compared to Heyl negative cells many but not all high-rank thyroid lineage drivers are enriched in Heyl+ cells (Supplementary dataset 3). *In vivo*, we find *Heyl* mRNA expression predominantly in cells facing the aortic sac to which the descending thyroid primordium adheres (Fig. 3h, i) and more broadly as the midline thyroid subsequently grows bilaterally (Supplementary Fig. 6a-a’’). By contrast, and consistent with scRNAseq data, the developing Ubb is entirely Heyl negative (Supplementary Fig. 6b-b’). At E15.5, the thyroid parenchyma shows weaker *Heyl* mRNA expression whereas, remarkably, strongly Heyl+ cells appear in the stromal compartment at a much higher frequency than in neighboring embryonic tissues (Supplementary Fig. 6c-c’). Heyl^high^ interstitial cells are occasionally present also in the adult thyroid and can be distinguished from the microvasculature (Supplementary Fig. 6d-d’’). Since these strongly Heyl+ cells are Nkx2-1 negative (Supplementary Fig. 6e) they are likely not identical to parafollicular C-cells. Finally, Heyl is upregulated in a mouse model of papillary thyroid cancer generated by mutant *Braf* (encoding BRAF^V600E^ oncoprotein) conditionally expressed under the thyroglobulin promoter (Supplementary Fig. 7a-g; [46]); the reduced expression of full-length Heyl and signs of altered Heyl turnover that accompanies redifferentiation of tumor cells in Braf mutant mice treated with a specific Braf kinase inhibitor support previous notions of Notch pathway involvement in BRAF^V600E^-induced thyroid tumorigenesis [47]. The nature of Heyl^+^/Nkx2-1^-^/Pax8^-^ cells emerging during thyroid organogenesis remains elusive.

### The predicted Ubb-GRN implicated in C-cell differentiation displays ancestral features of pharyngeal pouch development

Pathways implicated in neuroendocrine differentiation are strongly enriched in cluster 4 and to a lesser extent in clusters 6 and 8 confirming their Ubb/C-cell lineage identity (Fig. 4a, b). The predicted Ubb-GRN consists of 28 TFs of which some show a gradually increased expression in clusters 6→ 8→ 4 likely corresponding to the course of progenitor differentiation to C-cells (Table 1, right panel; Fig. 4a, b). Some Ubb-associated TFs are additionally upregulated in clusters 7 and 9 (Fig. 4c) supporting their Ubb relationship as revealed by pseudotime cluster currents (Fig. 2a-c, f). Besides Nkx2-1, only four Ubb-TFs (Id3, Nfib, Hoxb2 and Foxp2) are shared with the thyroid-GRN indicating high lineage specificity. The number of predicted Nkx2-1 target genes is higher (88) than in thyroid clusters, and although 25 are shared many of them are differentially regulated in Ubb vs thyroid lineage cells (Supplementary Table 2 and 3).

**Fig. 4.**
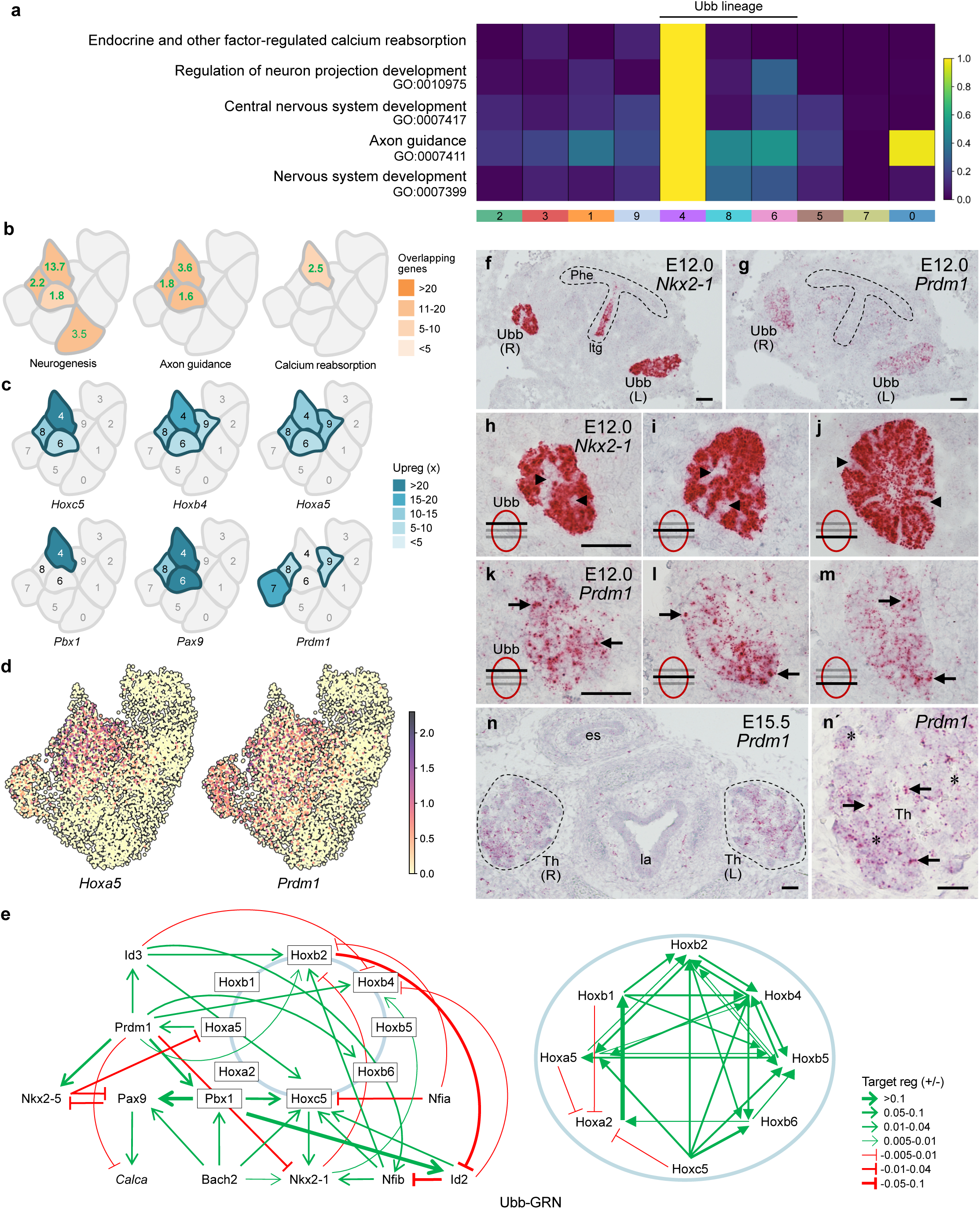
Gene enrichment and predicted gene regulatory network implicated in neuroendocrine differentiation of thyroid C-cells. **a** Heatmap of a curated list of pathways associated with neuronal development and neuroendocrine differentiation. Heat color intensity indicates the row normalized −log10 adjusted p-values (Benjamini–Hochberg) obtained from an enrichment analysis (with EnrichR) of the genes differentially upregulated (adjusted p-value < 0.01, log2-fold change > 1.0) in each cluster over a background comprising the remaining clusters. Clusters are indicated by numbers and colors according to Fig. 1i. **b** Cluster-specific enrichment of Ubb pathway-associated genes curated from GO Biological Process with −log10 adjusted p-values indicated (green). Color intensity corresponds to number of genes overlapping between the list of genes enriched in the pathway and the upregulated genes (adjusted p-value <0.01, log2-fold change >1.0) in the corresponding cluster. **c** Selection of genes encoding transcription factors (TFs) primarily identified in the predicted gene regulatory network of the Ubb lineage (Ubb-GRN; see below) and being variably upregulated in Ubb clusters 4, 6 and 8 (Table 1b). Curated from lists comprising all enriched genes per cluster. **d** UMAPs of indicated genes enriched in Ubb clusters. **e** Schematic gene regulatory network in Ubb lineage cells (Ubb-GRN) predicted by CellOracle network inference from multiomic profiles. Sharp (green) or blunt (red) arrows indicate source-target direction and arrow thickness represents positive or negative average cluster-specific GRN TF-target interaction scores of up- and downregulated genes. Circular connections indicate the predicted Hox gene subnetwork detailed in the right panel. **f-n’** RNAscope of *Nkx2-1* and *Prdm1* expression in the Ubb. **f, g** Overviews with endoderm outlined. **h-m** Details from serial sections. Arrowheads indicate Nkx2-1 negative cells; arrows indicate *Prdm1*^high^ cells. **n** *Prdm1* expression in thyroid lobes (encircled). Close-up of right lobe portion is shown in (n’). Arrows and asterisks indicate Prdm1^low^ and Prdm1^high^ cells, respectively. Phe Pharyngeal endoderm, ltg laryngotracheal grove, Ubb ultimobranchial body, Th thyroid, L left, R right, es esophagus, la larynx. Scale bars: 100 µm.

Consistent with a documented role in the development of inferior pharyngeal pouches [48, 49], numerous Hox genes as well as the heterodimerizing Hox partner Pbx1 are strongly upregulated in the Ubb lineage (Table 1, right panel) and also encountered among the strongest predicted Ubb lineage drivers (rank order): Hoxc5 (1), Hoxb4 (6), Pbx1 (11), Hoxb6 (17), Hoxb5 (34) and Hoxa5 (38) (Supplementary Dataset 2). Some Hox family members display a broader expression pattern arguing for cluster 9 belonging to the Ubb lineage (Fig. 4c, d) and interact broadly in the predicted Ubb-GRN (Fig. 4e). Being predominantly expressed in cluster 4 (Table 1, right panel), Pbx1 upregulates Hoxc5 which in turn target Nkx2-1 and Pax9, the only lineage-specific TF predicted to stimulate calcitonin expression (Fig. 4e). Altogether this forms a plausible regulatory subnetwork of C-cell differentiation.

Prdm1, a developmental TF of importance to caudal pharyngeal development [50, 51], is enriched in clusters 7-9 (Fig. 4c) and accounts for no less than 20% of Ubb target gene associations. According to the proposed cluster transition of the Ubb/C-cell lineage, Prdm1 is likely downregulated as C-cells differentiate (Fig. 4c; Table 1, right panel), confirmed by Prdm1 is the only predicted suppressor of *Nkx2-1* and *Calca* expression (Fig. 4e). Other major Prdm1 targets are Pbx1 and Nkx2-5 which in turn act antagonistically on Pax9 (Fig. 4e), further suggesting that Prdm1 is involved in keeping Ubb cells in an undifferentiated state. Notably, the tissues that express Prdm1 the most of all embryonic neck structures are the Ubbs (Fig. 4f-m); the expression is however heterogenous, more than that of Nkx2-1. Later, in the prospective thyroid lobe, Prdm1 is weakly expressed throughout the parenchyma although more strongly in scattered cells of presumed Ubb origin (Fig. 4n-n’). Collectively, this validates Prdm1 as a new biomarker of the Ubb/C-cell lineage confined to a subpopulation of immature cells which likely comprise C-cell precursors.

### Distinct growth kinetics of embryonic thyroid and Ubb are reproduced by single-cell clustering

We next wanted to examine whether single-cell transcriptome profiling might recapitulate key features of thyroid and Ubb cell propagation *in vivo* intending to identify alterations of potential importance for a compound gland to be formed. From previous studies [15, 52], as illustrated (Supplementary Fig. 8a-c), it is known that the growth kinetics differ between thyroid and Ubb lineage cells likely depending on different ratios of growth-prone progenitors and differentiated cells ceasing to proliferate. In the computed model, we find the distribution of G1, S and G2M cells and *Mki67*^+^ cells essentially mimics the lineage-specific cell proliferation rates, decreasing in Ubb and increasing in thyroid clusters across Eday (Supplementary Fig. 8d-f). Differential regulation of growth is further suggested from the predicted regulatory subnetwork of cyclin D genes indicting downregulation in Ubb lineage cells only (Supplementary Fig. 8g).

Most Ubb cells thus exit cell cycle prior to contact with the midline thyroid primordium is established (Supplementary Fig. 8a, b). Before this occurs the Ubb converges from a pseudostratified epithelium to a solid spheroid with densely packed proliferating cells (Supplementary Fig. 9a-e).

Intercellular cohesiveness is presumably maintained by the appearance of E-cadherin/Cdh1-based focal adhesions (Supplementary Fig. 6a, d). Many Ubb cells reverse polarity as evidenced by basal translocation of centrioles (Supplementary Fig. 6b-b’, c-c’), accompanied by lumen involution, constriction of the apical cytoplasm, and disassembly of Muc1 lining the apical surface (Supplementary Fig. 9e-e’’, f). Oriented cell division likely contributes to epithelial multilayering (Supplementary Fig. 9g-g’). During the process of major epithelial remodeling the Ubbs maintain cohesiveness and a compacted structure, which is evident also after being completely enclosed by overgrowing thyroid lineage cells (Supplementary Fig. 9h-h’).

### Basement membrane reorganization associated with differential expression of extracellular matrix constituents precede Ubb merging with the thyroid primordium

Attempting to elucidate the molecular signature and putative mechanisms underlying morphogenesis into a compound organ, we first examined clusters 0-9 broadly for possible differential gene expression of extracellular matrix (ECM) and basement membrane (BM) constituents. UMAP embedding of Gene Ontology panels showed that the collective scores of ECM- and BM-associated genes (GO:00311012, n=530; GO:0005604, n=123) differ only weakly (Supplementary Fig. 10a, b). Further, while differentially upregulated genes in each cluster are enriched for various ECM-associated ontology terms, we do not observe clear lineage differences apart from clusters 8 and 9 being almost blank (Supplementary Fig. 10c). Likewise, laminin complex genes show generally low expression levels with minor variations among clusters, the only remarkable exception being Pmp22 which is strongly upregulated in clusters 0-3 but barely detectable in clusters with Ubb identity (Fig. 5a). Pmp22 is a multifunctional protein mostly known as laminin binding partner in peripheral nerves [53], but is also a tight junction constituent and regulator [54, 55]. During embryogenesis, Pmp22 is expressed in mesodermal derivatives and the primitive gut tube [56]. Pmp22 expression prevailing across Eday specifically in the embryonic thyroid is validated by RNAscope (Supplementary Fig. 11a-d).

**Fig. 5.**
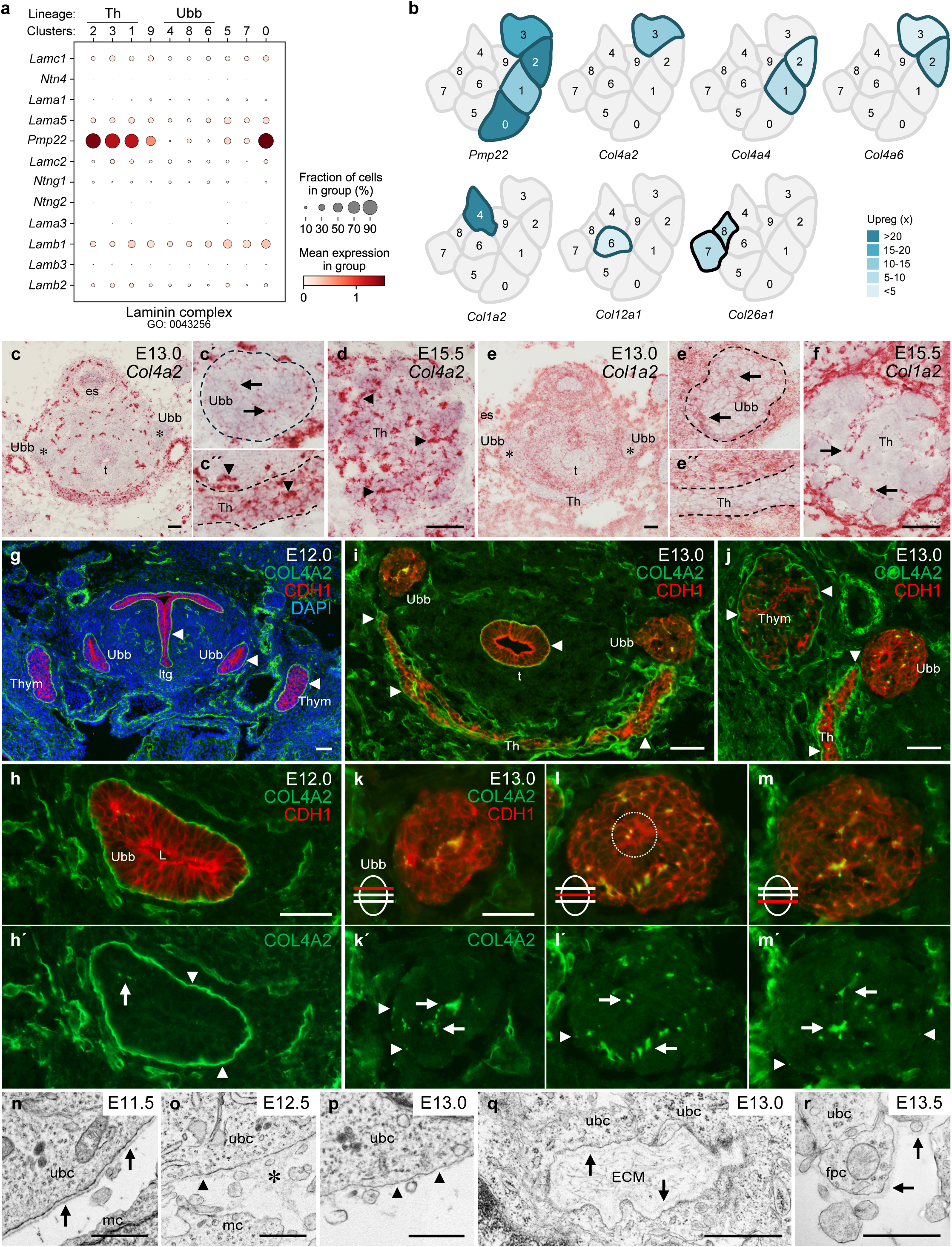
Differential expression of collagens and basement membrane dynamics in the developing thyroid and Ubb. **a** Dot plot displaying the expression pattern of laminin complex genes across clusters. Dot size indicates the fraction of cells in the cluster expressing the gene, and the color indicates the mean log2-normalized expression of the gene. **b** Cluster-specific enrichment of *Pmp22* and collagen I and IV genes. Curated from lists comprising all enriched genes per cluster. Corresponding data on *Col12a1* and *Col26a1* are shown to illustrate differential expression pattern. **c-f** Expression pattern of *Col4a2* (**c, d**) and *Col1a2* (**e, f**) at E13.0 (c, e) and E15.5 (d, f) revealed by RNAscope (overviews of both gene transcripts are shown in Supplementary Fig. 8). Ubb and midline thyroid are shown at high power and outlined in (c’-c’’, e’-e’’). Asterisks in (c, e) mark Ubb. Arrowheads indicate Col4a2^+^ capillary endothelium. Arrows indicate Col4a2^+^ cells in (c’) and Col1a2^+^ cells in (e’, f). **g-m** Double immunofluorescence of COL4A2 and CDH1/E-cadherin; except for (g), DAPI nuclear stain was omitted for improved resolution of collagen distribution. Overviews (g, i, j) and high-power images (h, k-m) at E12.0 and E13.0, respectively; (k-m) are serial sections of the same specimen. Single channel (green) images are shown in (h’, k’-m’). Arrowheads indicate contiguous collagen IV^+^ BMs. Arrows in (k’-m’) indicate collagen IV deposits. Outer border of Ubb is outlined in (k’-m’). **n-r** BM alternations revealed by transmission electron microscopy. **n** Intact Ubb-BM at E11.5. **o** Ubb-BM in dissolution at E12.5. **p** Ubb-BM missing at E13.0. **q** Ubb internal deposits of extracellular fibrillar material. **r** Reformed BM inclosing both Ubb and thyroid lineage cells at E13.5. Arrows and arrowheads indicate BM-covered and -denuded areas, respectively. Asterisk indicates electron-dense material of disintegrated BM. Th thyroid (primordium or lobe), fpc follicular progenitor cell, Ubb ultimobranchial body, ubc ultimobranchial body cells, es esophagus, t trachea, Thym thymus, ltg laryngotracheal grove, L lumen (encircled in (l)), mc mesenchymal cell. Scale bars: 100 (c-g, i-j), 50 (h, k-m) and 0.5 (n-r) µm.

Trajectory analysis identified Pmp22 ranking 190 of all thyroid lineage drivers (Supplementary Dataset 1) whereas it takes second place of genes negatively correlating with Ubb development (Supplementary Dataset 2). Surprisingly, Col1a2 ranks at 10^th^ place, and third place of non-TFs, among Ubb lineage drivers (Supplementary Dataset 2). This made us investigate the possibility of cluster-specific differences in collagen gene expression. Five out of six collagen type IV subfamily members are variably upregulated along with Pmp22 in thyroid lineage cells only (Fig. 5b, upper panel; omitted data: Col4a3 and Col4a5 are moderately enriched in clusters 1 and 3, respectively). By contrast, Ubb clusters exclusively show increased transcripts of *Col1a2* transcript in cluster 4 and *Col12a1* and *Col26a1* in clusters 6-8 (Fig. 5b, lower panel). Confirming *in silico* observations, the embryonic thyroid is absent of Col1a2 but show strong Col4a2 expression maintained across Eday in both parenchyma and microvasculature (Fig. 5c-f; Supplementary Fig. 12). On the other hand, compared to other pharyngeal derivatives, mature Ubb cells are nearly devoid of *Col4a2* whereas *Col1a2*^+^ cells of presumed Ubb origin is occasionally present in the prospective thyroid lobe (Fig. 5c-f; Supplementary Fig. 12). Lineage specificity is further suggested by Pax9 and Prdm1 stimulates whereas Pax8 inhibits collagen type 1 expression in the predicted subnetworks (Supplementary Fig. 13a).

Poor expression of type 4 collagen in the developing Ubb was confirmed by immunostaining. Remarkably, Col4a2 is entirely lost from the Ubb-BM between E12.0 and E13.0 i.e. prior to close contact with thyroid primordium is established (Fig. 5g-j). Additionally, Col4a2 appears multifocally in the interior of the Ubb (Fig. 5h-m, h’-m’), Preserved Col4a2 expression in adjacent endoderm-derived organ including the thyroid primordium indicates collagen IV alterations are Ubb-specific, altogether suggesting that redistribution of BM constituents from outside to inside might play a role in Ubb epithelial remodeling.

### Breakdown of laminin in the Ubb basement membrane is cell-autonomous, Nkx2-1 dependent and facilitates mixing of Ubb and thyroid lineage cells

Electron microscopy revealed that the entire Ubb-BM is gradually disintegrated (Fig. 5n-p) and that fibrillar ECM material occasionally accumulates in between Ubb cells (Fig. 5q), suggesting a more general mechanism involving disassembly of possibly all BM constituents. Indeed, although laminin expression does not significantly differ among clusters and across Eday (Fig. 5a) and only few laminin genes are predicted Ubb-TF targets (Supplementary Fig. 13b), starting at E12.5 by discrete breaches in the Ubb-BM laminin mimics the redistribution of Col4a2 (Fig. 6a, b) whereas in earlier developmental stages the laminin envelope is intact in both Ubb and thyroid primordia (Supplementary Fig. 14). Notably, the Ubb is devoid of vascularization (Fig. 6b, c), indicating gaps in the Ubb-BM do not come about by invading microvessels, as simultaneously occurs in the midline thyroid (Fig. 6c, d). Ubb-laminin breakdown accelerates between E12.5 and E13.0 (Fig. 6e). Reduced lumen size without accompanying changes in Ubb volume (Fig. 6f, g) rule out the possibility of mechanical breaching of the Ubb-BM. Interestingly, laminin occasionally appear in the shrinking lumen (Fig. 6h-h’’), which suggests that apical secretion of laminin might trigger reverse polarization as a means to reshape the Ubb epithelium [57].

**Fig. 6.**
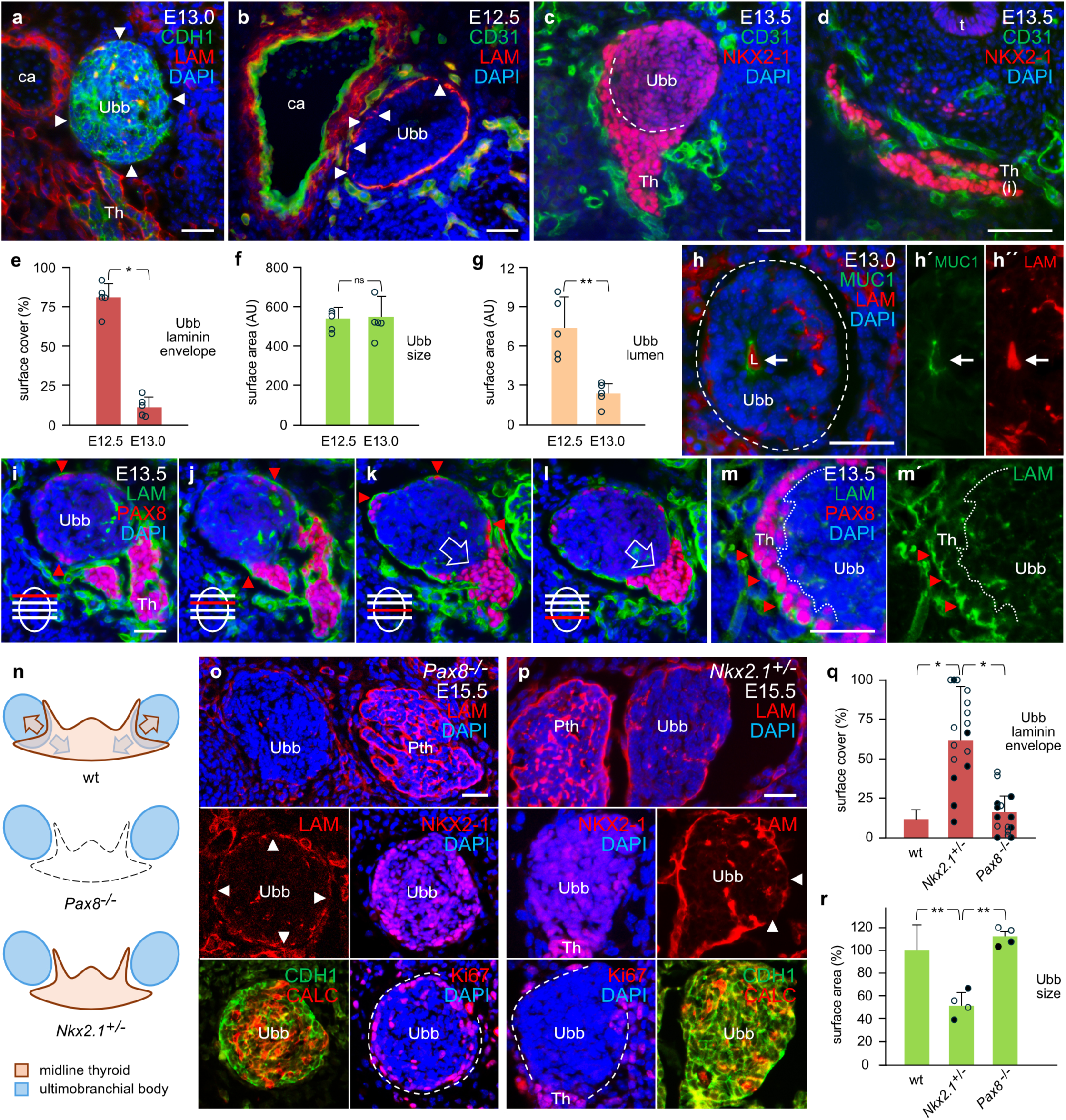
Disintegration and reassembly of laminin in thyroid primordia of wildtype, Pax8 null and Nkx2-1 heterozygous mice. **a-g** Gradual disappearance of basement membrane (BM) surrounding the Ubb. Double immunofluorescence of laminin (LAM), E-cadherin/CDH1, CD31 and NKX2-1 with DAPI nuclear staining. **a** Ubb devoid of laminin envelope at E13.0. **b** Gaps in the Ubb laminin envelope at E12.5 Arrowheads indicate denuded Ubb surface. For laminin distribution before E12.5, see Supplementary Fig. 10. **c, d** Vascularization of midline thyroid but not Ubb tissues. Interface between Nkx2-1^high^ thyroid cells and Nkx2-1^low^ Ubb cells outlined in (c). **e-g** Morphometric quantification of Ubb features at E12.5 and E13.0: surface cover of laminin (**e**), total volume (**f**) and lumen size (**g**); mean ± SEM (n=5; open circles indicate individual values): * p<0.0001; ** p<0.002. AU arbitrary units. **h** Ubb internal laminin deposits. Double immunofluorescence of LAM and MUC1. Arrows indicate luminal colocalization of markers in Ubb (outlined). **i-m** Reformation of BM around a common compartment. Double immunofluorescence of LAM and PAX8 with DAPI staining of nuclei. Serial imaging of the thyroid–Ubb fusion process (**i-l**) and closeup of laminin localization (**m**). Open arrows in (k, l) point at the BM-free interface of juxtapositioned thyroid and Ubb cells. Arrowheads (red) indicate laminin deposits associated with peripheral PAX8^+^ cells. Single channel motifs are shown in (h’-h’’ and m’) for improved resolution. Dotted lines in (m, m’) delineates the residual border between cell lineages. **n** Thyroid phenotypes of *Pax8* null (athyreotic) and *Nkx2.1* heterozygous (deficient fusion to Ubb) knockout (ko) mice (summarized from: [11, 16]). **o, p** Ubb features in *Pax8^-/-^* (**o**) and *Nkx2.1^+/-^* (**p**) E15.5 embryos. Immunofluorescence of LAM, NKX2-1, CDH1, calcitonin (CALC) and Ki67 with DAPI staining. Arrowheads indicate laminin^+^ and laminin-denuded surface areas. Dashed lines indicate Ubb perimeter. For further documentation of ko phenotypes, see Supplementary Figs. 11 and 12. **q, r** Morphometric quantification of laminin^+^ Ubb-BM (**q**) and Ubb size (**r**). Mean ± SEM (n=4 Ubb pairs from 2 mice for each mutant; * p<0.0005; ** p<0.05. Open and filled circles represent scores/section of the left and right Ubb). Comparison to corresponding mean wildtype (wt) percentual measurements obtained at E13.0 in (e, f). Th thyroid, Th(i) prospective isthmus, Ubb ultimobranchial body, t trachea ca carotid artery, L lumen, Pth parathyroid. Scale bars: 25 µm.

As the thyroid and Ubb gradually merge a new BM is formed, eventually investing the entire prospective thyroid lobe, in common to both lineage cells. Initially, this takes place by overgrowth of Pax8^+^ cells that eventually cover the entire Ubb surface. Serial sectioning revealed laminin deposits constituting the new BM is synthesized by the outermost yet incomplete Pax8^+^ cell layer (Fig. 6i-l), also confirmed by electron microscopy (Fig. 5r). At the same time, a limited stretch of the original thyroid BM is lost allowing direct contact of the two cell types precisely at the thyroid-Ubb interface (Fig. 6k-m).

Dynamic BM changes during this stage of thyroid organogenesis bringing cells of different embryonic origins together are obviously fundamental for the subsequent colonization and parafollicular positioning of C-cells. We used *Pax8* null athyreotic mice (Fig. 6n; [16]) to investigate whether breakdown of the Ubb-BM requires interaction with the thyroid primordium. Mutant mice were collected at E15.5 and serially sectioned to ensure Ubb identification by co-expression of Nkx2-1 and calcitonin. This showed that of all epithelial tissues in *Pax8^-/-^* embryos only the Ubb is denuded of laminin to the same extent as occurs in E13.0 wildtype mice (Fig. 6o, p; Supplementary Fig. 15).

Notably, calcitonin^+^/Nkx2-1^low^/Ki67^−^ cells accumulate in the Ubb center whereas calcitonin^−^/Nkx2-1^high^/Ki67^+^ cells attain a peripheral position (Figs. 6o). Altogether, this indicates that Ubb-BM degradation is a cell-autonomous process likely accomplished by undifferentiated Ubb cells.

Previous reports [11, 39, 58] have shown that *Nkx2-1^+/-^* mice are only mildly hypothyroid although the thyroid shows follicular abnormalities and is lacking C-cells due to a thyroid-Ubb fusion defect (Fig. 6n). The mechanism by which Nkx2-1 haploinsufficient C-cells fail entering the embryonic thyroid has not been elucidated. At E15.5, in heterozygous *Nkx2-1* mutants the Ubbs retain a spherical shape and cohesiveness of cells (Supplementary Fig. 16), although the thyroid lobes are rudimentary and much smaller than in wildtype embryos of the same age (compare size with wildtype in Fig. 8a). The Ubbs are also smaller, they do not contain any Nkx2-1^high^/Ki67^+^ cells, and the laminin envelope remains largely intact (Fig. 6p-r). Nkx2-1 gene dosage thus appears critical for efficient Ubb-laminin breakdown. This further infers that a preserved Ubb-BM will likely prevent thyroid- and Ubb-derived cells from getting in contact, which probably is required for their coordinated growth, thereby delaying thyroid organogenesis, as evident in heterozygous *Nkx2-1* null mice (Supplementary Fig. 16).

### Gene expression profiling of C-cells and C-cell precursors in the developing Ubb

In situ hybridization confirmed that calcitonin expression as confined to cluster 4 and E12.5 cells in the computed model is specifically expressed in the Ubb in mice of the corresponding age (Fig. 7a). From the heterogeneous expression pattern it is evident that only a few Ubb cells are yet committed to undergo neuroendocrine differentiation (Fig. 7b), confirmed by absence of calcitonin immunoreactivity at E12.5 and exceedingly few calcitonin^+^ cells present at E13.5 (Fig. 7c, d). We used the scRNAseq data to differentiate genes enriched in *Calca^+^*versus *Calca^−^* cells present in clusters 4, 6 and 8 at E12.5.

**Fig. 7.**
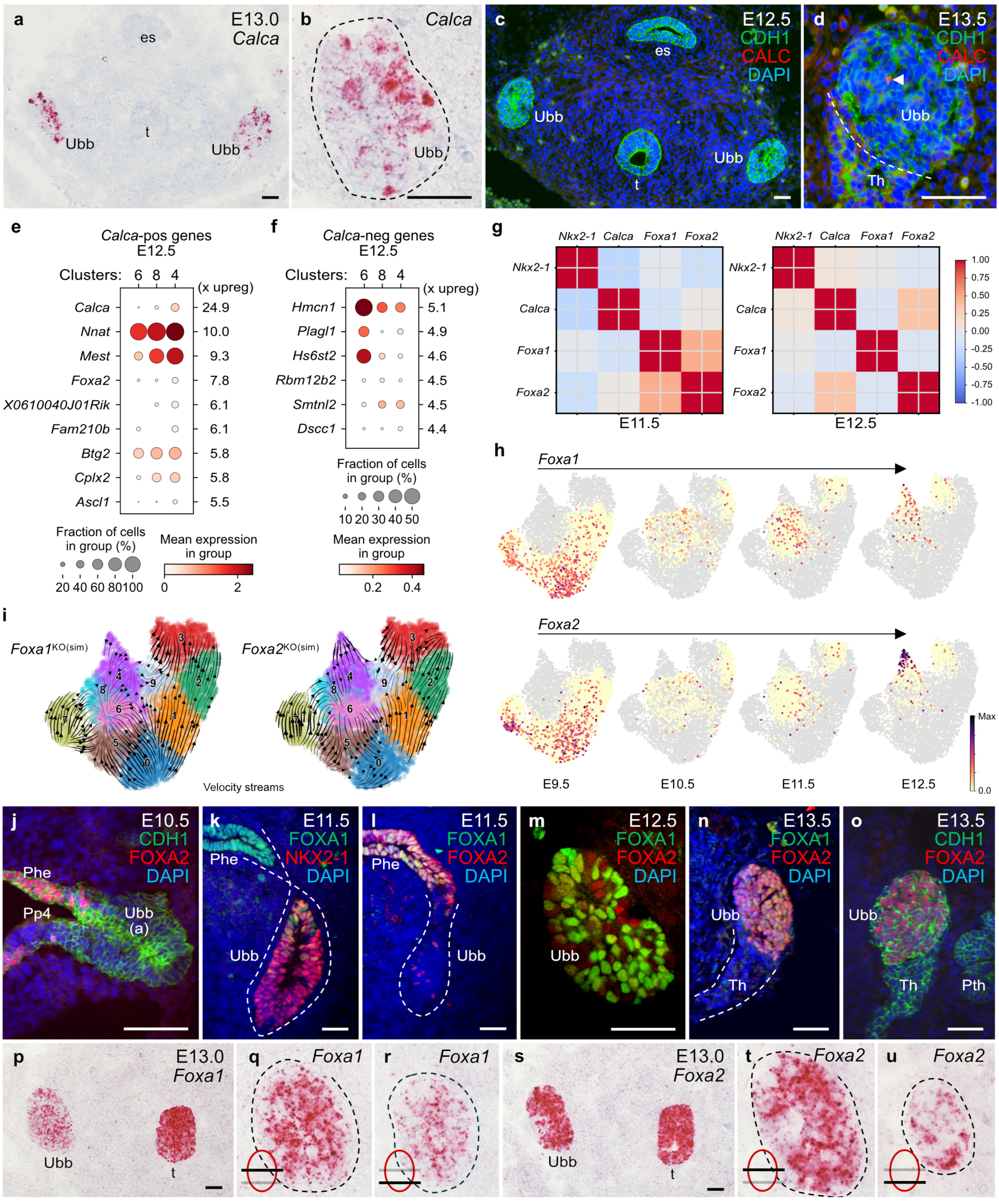
Ubb/C-cell lineage subpopulations distinguished by gene expression profiling related to differentiation state. **a, b** *Calca* mRNA expression in the mature Ubb as revealed by RNAscope. Outer Ubb border outlined in (b). **c, d** Immunofluorescence of calcitonin (CALC) prior to thyroid–Ubb fusion. Arrowheads in (d) indicate a CALC^+^ cell; border of primordia outlined. **e, f** Dot plots displaying upregulated genes (adjusted p-value < 0.01, log2-fold change >1.0) in the subset of E12.5 cells of the Ubb lineage clusters (4, 6 and 8) in *Calca* positive cells compared to *Calca* negative cells (**e**) and *Calca* negative cells compared to *Calca* positive cells (**f**). *Calca* positive cells are defined as those having normalized expression of *Calca* >0. The size of the dot indicates the fraction of cells from the subset in the cluster expressing the marker, and the color indicates the mean log2-normalized expression of the marker. Scores indicate the z-scores for each gene in the differential comparison between *Calca* positive and negative cells and vice versa. Enriched gene characteristics are shown in Supplementary Table 4. **g** Pearson correlation plots of *Nkx2-1*, *Calca*, *Foxa1* and *Foxa2* expression collectively confined to Ubb clusters 4, 6 and 8 at E11.5 (left panel) and E12.5 (right panel). **h** UMAPs visualizing differential expression of *Foxa1* (top panel) and *Foxa2* (bottom panel) across Eday. **i** *In silico* knockout simulations (ko(sim) of *Foxa1* (left panel) and *Foxa2* (right panel) based on GRN inference with CellOracle and predicted cell-state changes. Changes in gene expression between simulated knockouts and original cell states are displayed as velocity streams on the UMAP embedding of clusters 0-9 using scVelo. **j-o** Expression patterns of Foxa1 and Foxa2 across Eday. Ubb or thyroid primordium are outlined in (k, l and n). Double immunofluorescence of FOXA1/2, CDH1 or NKX2-1 with DAPI nuclear staining. **p-u** In situ hybridization of *Foxa1* (**p-r)**) and *Foxa2* (**s-u**) in mature Ubb. From serial sections with outer Ubb border outlined high power images. Overview *Foxa1/2* RNAscope images are shown in Supplementary Fig. 13. Ubb ultimobranchial body (a, anlage), Th thyroid, t trachea, es esophagus, Phe pharyngeal endoderm, Pp4 fourth pharyngeal pouch, Pth parathyroid. Scale bars: 100 (all except p) and 50 (p) µm.

Strongly up-regulated genes accompanying calcitonin expression are known or suggested neuronal and neuroendocrine markers (Fig. 7e; Supplementary Table 4, upper panel). Of particular interest, *Nnat* encoding neuronatin and ranking 42 among Ubb lineage drivers (Supplementary Dataset 2) is increasingly expressed along with the deduced cluster transition (clusters 6→ 8→ 4) of differentiating Ubb cells (Fig. 7e). Nnat is an imprinted gene mainly involved in brain development although regulation of insulin secretion [59] suggests a wider potentially neuroendocrine function, reinforced by recent findings of dysregulated *Nnat* in MTC tumor cells [60, 61].

In *Calca^−^* cells, enriched genes are predominantly found in cluster 6 some of which have documented or suggested roles in morphogenesis (Fig. 7f; Supplementary Table 4, lower panel). *Hmcn1* encoding hemicentin 1 is strongly enriched in cluster 6 and highest ranked among genes expressed in *Calca* negative cells (Fig. 7f). Interestingly, Hmcn1 belongs to the fibulin family of ECM proteins involved in BM assembly of type IV collagen [62] and maintenance of BM interacting with laminin [63]. Downregulation of Hmcn1 as observed along with C-cell differentiation (Fig. 7f) might thus be part of the mechanism that dismantles the Ubb-BM. Gene profiling of *Calca*^+^ and *Calca*^-^ Ubb cells further suggests that cluster 8 corresponds to an intermediate stage in Ubb/C-cell lineage development.

Foxa2 is enriched but Foxa1 is not (or is low) in *Calca*^+^ cells largely confined to Ubb cluster 4 (Fig. 7e), which is consistent with a documented role of Foxa2 enhancing calcitonin expression [64, 65]. Accordingly, *Foxa2* but not *Foxa1* correlates with *Calca* expression in E12.5 cells (Fig. 7g).

Both Fox genes are nonetheless crucial for organogenesis from foregut endoderm as signified by strong expression in pharyngeal derivatives with the notable exception of the thyroid primordium (Fig. 1e; Fig. 2f; Supplementary Fig. 17a-d); and [5, 17, 66]. However, although largely co-expressed at E9.5-10.5 i.e. accompanying formation of the pharyngeal pouch from which Ubb derives, UMAP embeddings across Eday show that Foxa1^+^ and Foxa2^+^ cells are thereafter differently distributed with only small overlap (Fig. 7h), as also revealed by progressively diminished Pearson correlation scores (Fig. 7g). We previously showed that Foxa1 but not Foxa2 is co-expressed by Ki67+ Ubb cells [5]. The computed model thus confirms that Foxa1 rather than Foxa2 regulates propagation of immature Ubb cells corresponding to C-cell precursors.

*In silico* knockouts of *Foxa1* and *Foxa2* simulate relocation of both targets from cluster 4 to cluster 6 representing a more immature state in Ubb development (Fig. 7i). Interestingly, velocity streams to cluster 6 also account clusters 0, 5 and 7, which plausibly simulates pharyngeal pouch development featured by transient downregulation of Foxa1 and Foxa2 in the prospective Ubb (Fig. 7j-l), as previously reported [5]. It is noteworthy that *Foxa1* and *Foxa2* knockout simulations also predict velocity streams from cluster 0 to clusters 1 and 2 (Fig. 7i), inferring a possibility that loss of Foxa1/Foxa2 expression might trigger endodermal cells to adopt a thyroid fate. Indeed, absence of Foxa2 is required for induction of thyroid competence in anterior foregut endoderm derived from mouse pluripotent stem cells [66]. Similarly, velocity streams from cluster 9 to cluster 3 suggests the intriguing possibility that progenitor cells with putative dual lineage competence might be segregated into this cluster.

Consistent with present UMAP datasets and confirming previous reports [5], Foxa1 and Foxa2 are differentially expressed in the developing Ubb although most cells eventually co-express both (Fig. 7m-o). Foxa1/2 heterogeneity is nonetheless evident at the transcriptional level among Ubb cells approaching the time point of thyroid-Ubb fusion (Fig. 7p-u). GRN modeling reveals that Foxa1 and Foxa2 are differentially regulated (Supplementary Fig. 18). Pax9, which is a predicted transcriptional regulator of calcitonin (Fig. 4e), paradoxically downregulates Foxa2. Pax8 is the only predicted TF suppressing Foxa2 expression in thyroid lineage cells, which *in vivo* is evident already from the placode stage [17].

### C-cell precursors undergo epithelial-to-mesenchymal transition and acquire a Foxa1^+^/Cdh2^+^ migratory phenotype correlating with intrathyroidal dissemination

A remaining question is how thyroid follicular cells and C-cells become fully integrated in the compound gland. In the prospective thyroid lobe, propagation of the thyroid lineage occurs by branching morphogenesis during which proliferating cells differentiate and reorganize into follicles (Fig. 8a-c) [15]. Follicular cells producing thyroglobulin (Tg), the prohormone to thyroxin, can be identified at a pre-follicular stage by ubiquitous expression of Muc1 (Fig. 8b) which redistributes to the apical membrane and thus colocalize with luminal Tg in newly formed follicles (Fig. 8c-c’’). Concurrently, the Ubb rudiment consisting of Nkx2-1^low^/Muc1^-^ cells remains centrally positioned while some Ubb-derived cells begin to relocate outwards (Fig. 8a, b, d). Before follicles develop their own basal laminae both lineage cell types thus reside in the same compartment enclosed by a shared BM surrounding the entire lobe (Fig. 8b, e).

**Fig. 8.**
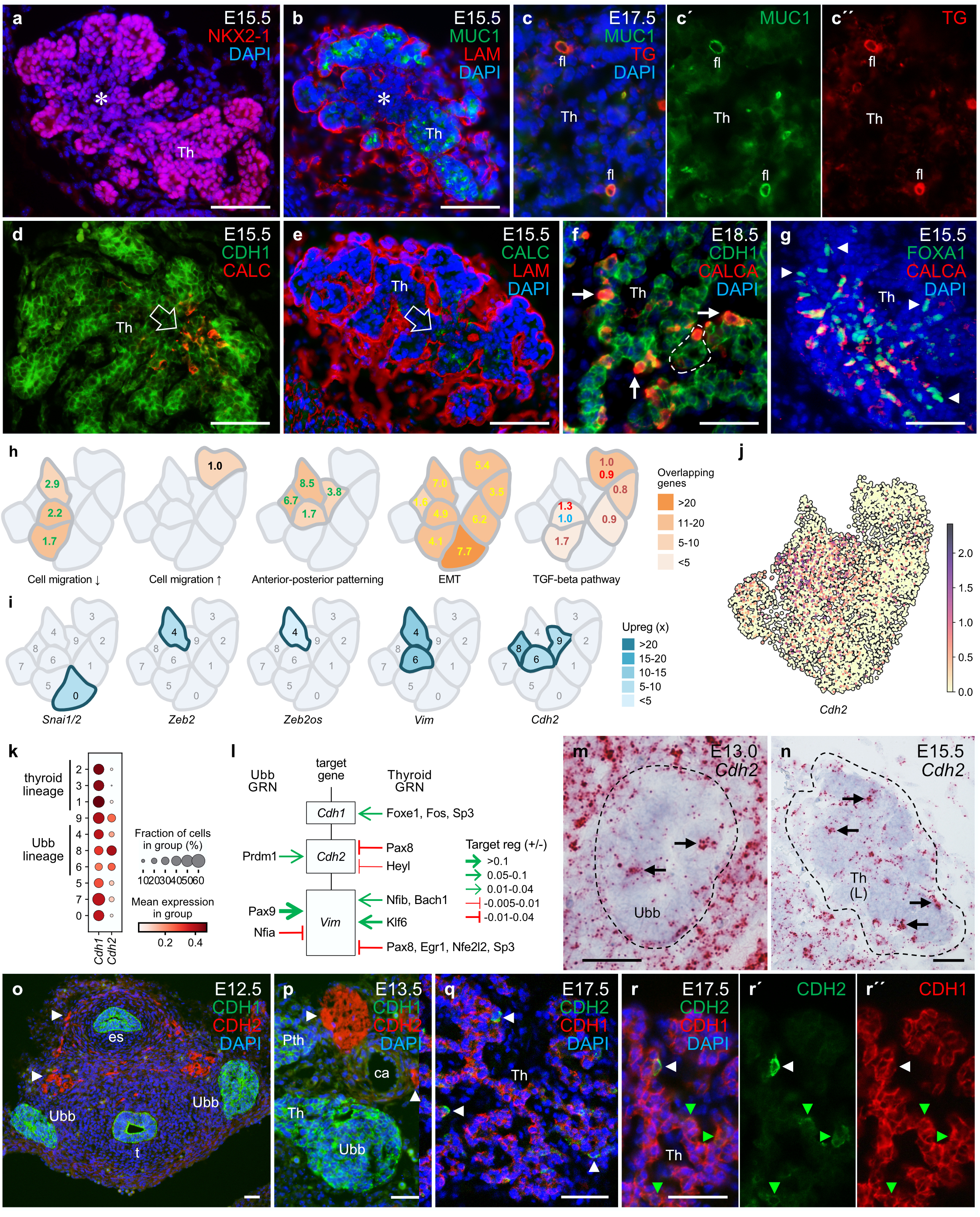
Post-fusion thyroid morphogenesis and upregulation of EMT genes in Ubb/C-cell lineage cells colonizing the embryonic thyroid. **a-g** Propagation of thyroid and Ubb lineage cells forming the final thyroid lobe histoarchitecture. Double immunofluorescence with DAPI counterstaining. **a, b** Branching morphogenesis of NKX2-1^high^ thyroid cells enclosed by a common laminin/LAM^+^ basement membrane. Asterisks mark residual Ubb featured by NKX2-1^low^/MUC1^-^ cells. **c** Colocalization of thyroglobulin (TG) and MUC1 in newly formed follicles; separate channels shown in (c’, c’’). **d, e** Embryonic C-cells starting to migrate; open arrows indicate central location of calcitonin/CALC^+^ cells. **f** Tissue distribution of C-cells. CALC^+^ cells indicated by arrows; a newly formed follicle with parafollicular C-cell is encircled. **g** Tissue distribution of C-cells and C-cell precursors. Arrowheads indicate FOXA1^+^/CALC^−^ cells. **h** Cluster-specific enrichment of the indicated process- or pathway-associated gene ontology terms curated from: GO_Biological_Process (green), GO Molecular Function (red), KEGG (blue), WikiPathway (brown), MSigDB_Hallmark (yellow) and Biocarta (black). Enrichment analysis (with EnrichR) was performed on the genes differentially upregulated (adjusted p-value < 0.01, log2-fild change >1.0) in each cluster over a background comprising the remaining clusters. Color intensity corresponds to the number of cluster enriched genes found in the corresponding ontology term with −log10 adjusted p-values (Benjamin-Hochberg). Related heatmap of EMT-associated pathways and processes is shown in Supplementary Fig. 15. **i** Cluster-specific enrichment of genes implicated in EMT. Curated from lists comprising all enriched genes (adjusted p-value <0.01, log2-fold change >1.0) per cluster. **j** UMAP embedding of *Cdh2* with cells outlined and colored by their log2-normalized expression. **k** Dot plot of *Cdh1* and *Cdh2* expression across clusters. Dot size indicates the fraction of cells expressing the marker and the color indicates their mean log2-normalized expression. **l** Transcriptional regulation of *Cdh1*, *Cdh2* and *Vim* predicted by CellOracle. Up- versus downregulation are indicated by sharp (green) or blunt (red) arrows and arrow thickness representing mean lineage-specific GRN TF-target gene scores. m, n *Cdh2* expression by RNAscope in Ubb (**m)** and thyroid lobe (**n**). Arrows indicate scattered *Cdh2^+^* cells within Ubb/thyroid tissues (encircled). Overviews of *Cdh2*-expressing embryonic tissues are shown in Supplementary Fig. 13. **o-r** Cdh1/Cdh2 expression by immunofluorescence. **o, p** In Ubb at E12.5 (o) and E13.5 (p). Arrowheads indicate Cdh2^+^ nerves. **q, r** In thyroid lobe, overview (**q**) and close-up (**r**); r’-r’’ show separate green and red channels for improved visualization. Arrowheads indicate CDH1^low^/CDH2^high^ (white) and CDH1^high^/CDH2^low^ (green) cells. Th thyroid, L left (lobe), fl follicle lumen, Ubb ultimobranchial body, Pth parathyroid, es esophagus, t trachea, ca carotid artery. Scale bars: 100 (a, b, d, e, m-p), 50 (c, f) and 25 (g, q) µm.

From previous mouse studies we know that intrathyroidal dissemination of C-cells is a regulated process involving heterotypic cell-cell interactions [67]. Embryonic C-cells seemingly migrate alongside the branching parenchyma and attain one by one or in small groups a parafollicular position as new follicles are about to develop (Fig. 8f). Immunofluorescence reveals Foxa1^+^/Calcitonin^−^ C-cell precursors are more frequent in the forefront of migration streams (Fig. 8g), suggesting that the capability to migrate might be constrained as cells undergo neuroendocrine differentiation and become resident. The likelihood of counterbalancing anti- and promigratory mechanisms featuring the mature Ubb is suggested by enrichment of genes associated with inhibited cell motility in clusters with Ubb identity (Fig. 8h). In fact, only cluster 3 shows a pathway signature consistent with increased cell migration, which *in vivo* corresponds to bilateral growth of progenitor cells forming the thyroid isthmus. Moreover, genes implicated in cell positioning along the anterior-posterior axis are selectively upregulated in all Ubb clusters including cluster 9 (Fig. 8h). Presumably, this relates to Hox gene predominance in the Ubb-GRN (Fig. 4e), potentially mediating positional information that might prevail in the Ubb rudiment residing in the prospective thyroid lobe.

We next asked whether epithelial-to-mesenchymal transition (EMT) might be involved in embryonic C-cell migration. Surprisingly, EMT-associated pathways are enriched in nearly all clusters and transforming growth factor-beta (TGF-beta) signaling scores in both thyroid and Ubb/C-cell lineages (Fig. 5e, right panel), Heatmaps from curated lists of ontology terms suggest different EMT-associated pathways are involved (Supplementary Fig. 19). Accordingly, vimentin ranking high (98) among Ubb lineage drivers (Supplementary Dataset 2) is strongly enriched in clusters 4 and 6 (Fig. 8i). That Ubb cells likely are committed to undergo EMT is further supported by enrichment of *Zeb2* and *Zeb2os* in cluster 4 (Fig. 8i). Interestingly, Zeb2 is a repressor of Nkx2-1 [68], which is downregulated in mature Ubb cells (Fig. 8a), and Zeb2os, a long non-coding RNA, facilitates Zeb2 expression and function [69]. Based on enrichment data, Snai genes might have a role in early Ubb development but are likely not involved in Ubb maturation (Fig. 8i).

N-cadherin/Cdh2, another high ranked (119) Ubb lineage driver (Supplementary Dataset 2) and well-known EMT effector primarily by modifying intercellular adhesion [70, 71], is enriched in Ubb clusters although less in cluster 4 (Fig. 8i, j). Dot plot comparison indicates that *Cdh1* and *Cdh2* are reciprocally up- and downregulated, Cdh2>Cdh1 in cluster 8 and Cdh1>Cdh2 in cluster 4 with equal transcript levels in cluster 6 (Fig. 8k), suggesting cadherin expression is dynamically altered as the Ubb matures, consistent with highest *Chd2* transcript levels in C-cell precursors. Moreover, cluster-specific GRN inference predicts Cdh1 and Cdh2 are differentially regulated (Fig. 8l), which infer upregulation of Cdh2 by Prdm1 and vimentin by Pax9, conceivably corresponding to their individual expression pattern (Fig. 4c, Fig. 8i). *In vivo*, *Cdh2* is widely expressed in neuronal and mesenchymal embryonic tissues but absent in epithelia of endoderm origin with the exception of Ubb, prospective thyroid lobe and laryngotracheal epithelium (Fig. 8m, n; Supplementary Fig. 17e, f). Similar to calcitonin, no N-cadherin/Cdh2 immunoreactivity is found in the Ubb before merging with the thyroid primordium (Fig. 8o, p). Moreover, scattered Cdh2^+^ cells are present in the thyroid lobe, but their numbers are generally lower than the calcitonin positive cells (Fig. 8q; compare with Fig. 8f). Separate visualization of fluorophores revealed that cells expressing high Cdh2 levels have much reduced immunostaining of Cdh1 whereas Cdh2^low^ cells, hardly detectable in superimposed images, co-express E-cadherin at similar levels to that of thyroid lineage cells (Fig. 8r-r’’), thus reproducing the corresponding scRNAseq findings (Fig. 8k).

In summary, the computed model of C-cell development, validated by expression analysis in wildtype and mutant embryos, suggests that Foxa1^+^ C-cell precursors exhibit a promigratory phenotype that once deployed will pave the way for C-cells colonizing the gland. In this process, direct contact with thyroid lineage cells is required to unleash their full migrating potential for which cadherin switch – downregulation of E-cadherin/Cdh1 and upregulation of N-cadherin/Cdh2 – appears important.

### Invasive tumor cells possess C-cell precursor features in thyroid carcinoma of dual lineage origin

In the adult thyroid, Cdh2^+^ cells equals in number that of C-cells and are similarly distributed in the lobe center holding a parafollicular or intraepithelial position (Fig. 9a, b). Inferring N-cadherin as a new biomarker of the terminal neuroendocrine phenotype, Cdh2 is co-expressed with Cdh1, Foxa1 and Foxa2 in the TT cell line of human MTC origin (Fig. 9c). In general, MTC is a silent disease due to slow progression and often diagnosed at an advanced stage difficult to treat, hence disease-related death is significant. C-cell-derived malignant cells are arguably more invasive and metastatic than most follicular cell-derived tumors. We therefore wanted to investigate whether MTC tumor cells might recapitulate some developmental features that evidently differ between Ubb/C-cell and thyroid lineage cells.

**Fig. 9.**
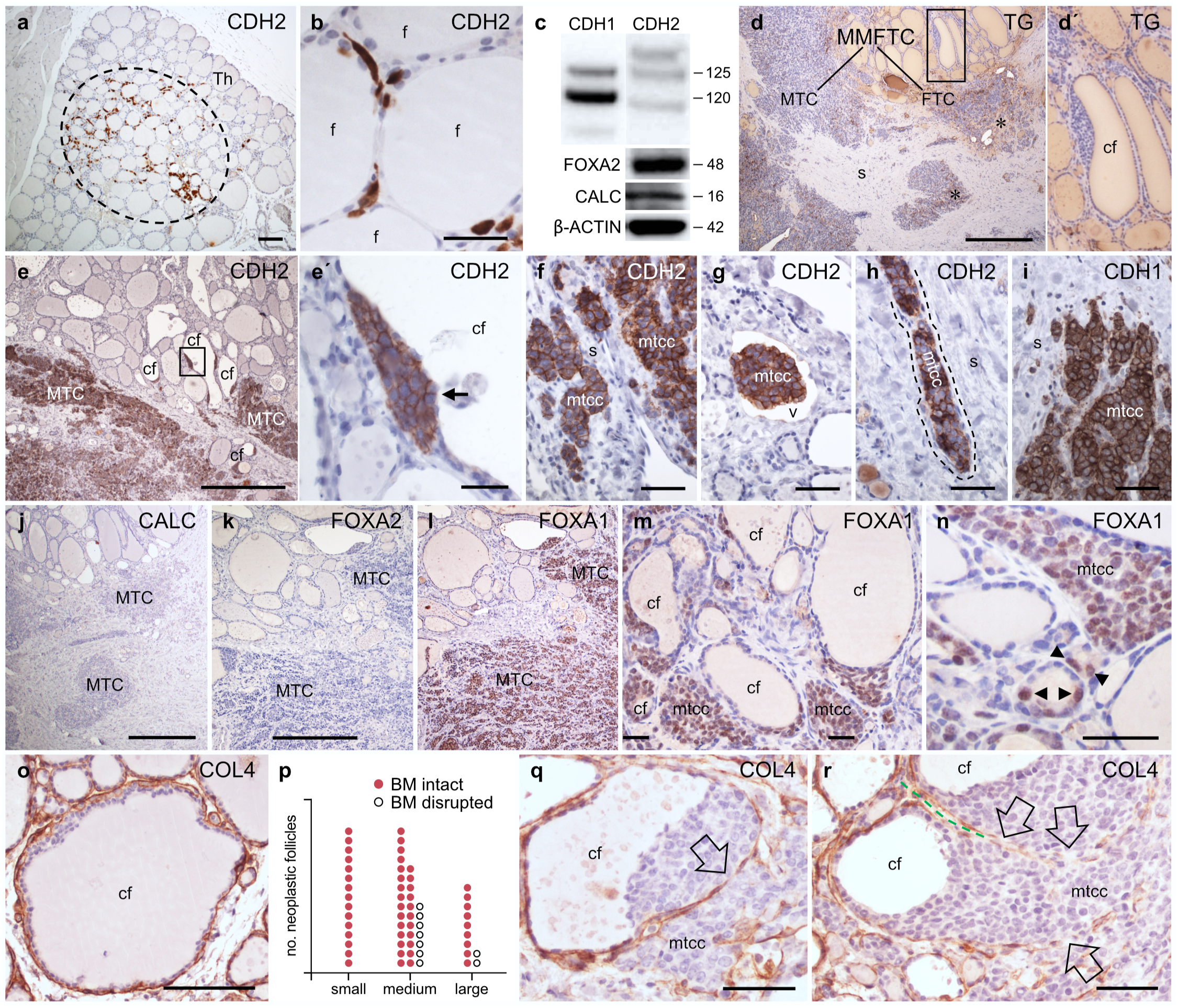
Expression of N-cadherin/Cdh2, Foxa1/2 and collagen IV in medullary thyroid carcinoma cells related to tumor invasiveness. Immunohistochemical (IHC) staining of formalin-fixed paraffine-embedded tissue sections and Western blot analysis. **a, b** Distribution of CDH2^+^ cells in mouse thyroid gland, overview (**a**) and detail (**b**). **c** Co-expression of CDH1/2, FOXA1/2 and calcitonin/CALC in TT cell line; uncropped gel data are provided in Supplementary Dataset 3. **d-p** Histological features of mixed medullary-follicular thyroid carcinoma (MMFTC). **d** Immunostaining of thyroglobulin/TG; **d’** show high power of boxed area in (d). Asterisks indicate TG^+^ MTC tumor portions. **e** CDH2 expression in MTC tumor and compound neoplastic follicles; **e’** shows high power of boxed area in (e) with arrow indicating CDH2^+^ cells facing the follicle lumen. **f-h** Stromal and intravasal infiltration of CDH2^+^ medullary thyroid carcinoma cells (mtcc); microvessel wall outlined in (h). **i** CDH1 expression in invasive tumor cells. **j, k** The MTC tumor component is lacking calcitonin/CALC (**i**) and FOXA2 (**j**) immunoreactivity. **l-n** FOXA1 expression in MTC tumor and compound neoplastic follicles, overview (**l**) and close-ups (**l, m**). Arrowheads in (m) indicate FOXA2**^+^** cells in microfollicles. **o-r** COL4^+^ basement membrane (BM) alterations in compound neoplastic follicles. **o** Large compound follicle with preserved BM. **p** Incidence of follicles with intact and disrupted BM; encountered from x20 images of four different areas of the follicular tumor and distributed in three arbitrary follicle size categories, each dot representing a single follicle profile. **q** Tumor cells breaking through the disrupted BM. **r** Completely disintegrated BM with massive tumor cell invasion; open arrows indicate loss of COL4^+^ BM segments with the interspace of adjacent follicles outlined in green for clarity. MMFTC mixed medullary follicular thyroid carcinoma, MTC medullary thyroid carcinoma, FTC follicular thyroid carcinoma, Th thyroid (lobe), f follicle, cf compound follicle, mtcc medullary thyroid carcinoma cells, s stroma, v vessel. Scale bars: 500 (d, i), 100 (a, e, j-l, n) 50 (f-h, p) and 25 (b, e’, m, o) µm.

In a recent case of mixed medullary and follicular cell-derived carcinoma (MMFTC), a rare tumor entity [72, 73], the primary tumor is composed of synchronously developed compound neoplastic follicles although the neuroendocrine component is conspicuously more invasive than the follicular counterpart (Fig. 9d-d’). The follicular tumor consists of differentiated cells revealed by Tg positivity, whereas the neuroendocrine tumor is calcitonin negative fulfilling the criteria of a hormone-negative MTC [74]. Further phenotyping showed strong CDH2 immunostaining confined to the neuroendocrine tumor (Fig. 9e). Intriguingly, compound follicles display clusters of Cdh2^+^ tumor cells that occupy a continuum of parafollicular or intraepithelial positions facing both lumen and extrafollicular space (Fig. 9e’). Invasive cells express both Cdh2 and Cdh1 and retain cohesiveness indicating local spreading occurs by collective migration (Fig. 9f-i). Consistent with possible parallels to C-cell precursor features, the neuroendocrine tumor cells are generally Foxa1 positive but completely lacking calcitonin and Foxa2 immunoreactivity (Fig. 9j-l). In compound neoplastic follicles, Foxa1^+^ cells show the same distribution as Cdh2^+^ cells indicating they are identical (Fig. 9m). The occurrence of small, newly formed follicles consisting of both Foxa1^+^ and Foxa1^-^ cells (Fig. 9n) strongly suggests that tumorigenesis is oligoclonal and that neoplastic follicles develop synchronously by cells with different clonality and possibly of different lineage origins. Independently of size, most compound follicle profiles display an intact BM (Fig. 9o, p), indicating that collagen IV synthesis and assembly is not compromised. However, in occasional follicles with more neuroendocrine cells accumulated within it is evident that the Col4+ BM is disrupted allowing tumor cells transmigrate extrafollicularly (Fig. 9q). Moreover, complete resolution of the follicular BM correlates with massively invasive tumor growth (Fig. 9r).

We previously showed that metastatic MTC maintains Foxa1 expression but transiently downregulate Foxa2 and E-cadherin [5]. The present findings indicate that dedifferentiated MTC tumor cells adopt C-cell precursor-like properties and infer a potential role of N-cadherin/Cdh2 in tumor invasiveness. Since released type IV collagen and laminins trigger EMT and thereby promote cell migration [75–77], it is possible that disintegration of the follicular BM might be a pathogenetic factor for MMFTC attaining an invasive phenotype. Therefore, in embryonic development, elucidating the mechanism by which Ubb/C-cell lineage cells maintain cohesion and do not precociously migrate despite a degraded BM might reveal means and a potential strategy counteracting tumor invasiveness in MTC.

## Discussion

This study provides the first comprehensive analysis of mammalian thyroid development linking global single-cell molecular signatures of embryonic lineage trajectories to organogenesis of the composite thyroid gland. The predicted transcriptional regulation of lineage propagation across Eday corresponding to *in vivo* morphogenesis of organ primordia involves unique constellations of networking TFs and lineage-specific target genes. Both thyroid and neuroendocrine differentiation commence before primordial tissues merge. Transcriptome profiling confirms a neuron-like phenotype of the Ubb/C-cell lineage, which firmly establishes the endodermal origin of thyroid C-cells previously indicated by lineage tracing [5]. The long-standing misconception of a neural crest ancestry [4], essentially based on quail-chick xenotransplant studies [78, 79], is now also disproven by recent findings of a universal pharyngeal endoderm origin of calcitonin-producing neuroendocrine cells throughout the animal kingdom down to the protochordate endostyle [80].

In thyroid lineage cells committed to a follicular fate, we identified several enriched TFs with no previously identified functions in thyroid development, and which are either transiently upregulated in the more immature cell stages (Crem, Elf3, Klf6, Nfe2l2, Nr4a2, Tbx3) or accompanying terminal differentiation (Fos, Jun > Bcl11b, Egr1, Fosb, Hivep3, Junb, Maf, Nr3c1, Sox9, Tcf4 > Arntl, Elk3, Ets1, Heyl, Nfic, Rorc). Most of these TFs participate in the predicted GRN subnetwork targeting or being targeted by one or more of the classical four thyroid TFs – Pax8, Nkx2-1, Hhex and Foxe1 – which are also upregulated in progenitor cells as they differentiate and acquire competence to synthesize thyroid hormone. Considering causal mechanisms of thyroid dysgenesis are unknown for the majority of patients diagnosed with congenital hypothyroidism (inactivating mutations of known developmental genes as *Nkx2-1* are infrequent), the present *in silico* observations open avenues searching for novel mutations or epigenetic dysregulation in patient cohorts and to functionally explore candidate genes with perturbation experiments. This was recently done for Sox9 [81], a newly discovered thyroid TF [15] which we also found enriched in thyroid lineage cells undergoing differentiation. The predicted thyroid-GRN, to some extent validated by functional experiments, and the multitude of associated candidate target genes should be regarded as a comprehensive framework of thyroid development, which likely will stimulate future in-depth studies. While *in silico* GRNs and perturbation experiments have limitations, they pave the way for further experimental investigation. As suggested from our present findings of Nkx2-1 regulating the expression of Pax8, monitoring subnetwork alterations across Eday will be required to fully comprehend how TFs interact and synergize in the developing thyroid, and potentially uncover key regulatory mechanisms that govern the function of thyroid lineage cells with distinct gene expression profiles. Recently, cell diversity was demonstrated for the Pax8 orthologous gene *Pax2a* by single-cell transcriptome analysis in zebrafish thyroid [82], and we found similar cell heterogeneities in the computed model of thyroid development, confirmed for Nkx2-1, Pax8 and Heyl by RNAscope double fluorescent staining of intact tissues. The dynamic expression of Heyl, a downstream target of Notch signaling [83–86], in undifferentiated embryonic cells and Braf mutant adult cells is of particular interest considering the newly discovered importance of the Notch pathway for maintaining thyroid endocrine function [87].

We identified several new candidate genes and validated the expression of some potentially involved in embryogenesis of thyroid C-cells. These genes comprise both TFs and putative target genes that are differentially expressed during Ubb development (early: e.g. Hoxa2, Meis2, Ripply3, Hmcn1); intermediate: e.g. Hoxb1, Hoxb5, Prdm1, Cdh2; and late: e.g. Hoxb4, Hoxb6, Hoxc5, Pbx1, Foxa2, Ascl1, Ret, Mest, Nnat, Zeb2, Col1a2). Yet other genes specifically identified in the Ubb/C-cell lineage i.e. Pax9, Foxa1, Foxp2, Hoxa5, Id2, Meis3, Meox1, Nkx2-5, Sox4 and Yxb1 are more stably expressed. Pax9, broadly expressed in pharyngeal endoderm [7] and instrumental for harvesting of embryonic cells subjected to single cell analysis [14], is thus enriched in both C-cell precursors and differentiated C-cells. Interpreted from the Ubb-GRN, Pax9 inhibits Foxa2 expression presumably in C-cell progenitors, upregulates vimentin and type I collagen in cells undergoing differentiation, and stimulates calcitonin expression suggesting Pax9 exerts multiple roles in different stages of Ubb development. This is conceptually similarly to Pax8 which is broadly expressed in all thyroid developmental stages and of crucial importance for both organogenesis and functional differentiation including generation of follicles [18, 35, 43]. Pseudotime analysis and knockout simulations essentially confirmed the expression pattern of Pax8 *in vivo* and thereby conceivably validated as a whole the predicted subnetworks of lineage-specific gene regulation.

We found that Prdm1, a transcriptional regulator of pharyngeal pouch development acting in concert with Tbx1 [51, 88], is specifically expressed in the Ubb and, moreover, that N-cadherin/Cdh2 is a predicted Prdm1 target in a subset of Ubb cells that possess C-cell precursor features. Further tracing of these cells suggested that N-cadherin/Cdh2 as part of an induced EMT program facilitate their dissemination into the embryonic thyroid. This process is evidently promoted by Nkx2-1, at least partly by degrading the Ubb-BM which does not happen in *Nkx2-1* heterozygous mutants. Since Prdm1 is predicted to strongly downregulate Nkx2-1, we hypothesize that reduced Nkx2-1 expression in the mature Ubb is required but not sufficient for cells to undergo EMT and disseminate. The developmental delay of both thyroid and Ubb in *Nkx2-1^+/-^*mice might thus simply reflect the natural process, the Ubb should not precociously disintegrate until contact with thyroid progenitors is established. Although we know from previous studies that Eph-ephrin signaling facilitates C-cells spreading within the thyroid [67], the precise mechanism unlocking Ubb cells from inhibited migration remains to be elucidated. As Prdm1 promotes cell migration both during embryogenesis and in tumors acting downstream of TGF-beta [89–91], it is possible that Prdm1 via Cdh2 might play a direct role for C-cell precursors to migrate and eventually become resident parafollicular C-cells. Since basically the reverse process takes place in MMFTC in which neuroendocrine tumor cells escape their initial parafollicular position and become invasive, it is plausible to assume that MTC tumor cells might adopt a developmental trait of partial EMT featuring embryonic C-cells and involving Cdh2 regulated by Prdm1.

Essentially validated by *in situ* hybridization, the computed model predicted Col1a2 is upregulated by Pax9 and Prdm1 suggesting that type I collagen might contribute to remodeling of the Ubb and the cells acquiring a promigratory phenotype. Previous cell line studies show that collagen I may trigger EMT by destabilizing E-cadherin at cell-cell contacts [92, 93] and upregulate N-cadherin/Cdh2 expression [94]. Type I collagen-induced EMT is mediated by binding to discoidin domain receptors Ddr1and Ddr2 [95, 96] and activation of the TGF-beta signaling pathway [97] in a ligand-independent manner [98]. Notably, Ddr stabilizes EMT by the activation of EMT-associated TFs and is also reciprocally regulated e.g. by Zeb1 [96, 99–102]. Although we did not find upregulation in any clusters and both Ddr genes ranked low among lineage drivers, it is yet possible that Col1a2 might act through Ddr1/2 influencing Ubb development.

Loss of major BM constituents during Ubb maturation similar in wildtype and athyreotic *Pax8* null mice indicates that Ubb-BM breakdown is attributed to a cell-autonomous mechanism likely programmed to forearm the process of merging with the thyroid primordium. Indeed, removal of the epithelial BM fence function is critical to bring the two lineage cell types in direct contact, a prerequisite for their mixing and tandem propagation in the prospective thyroid lobe. Differential expression of laminin complex member Pmp22 and several collagen genes particularly Col1a2 (in Ubb) and Col4a2-6 (in thyroid) and their corresponding GRN subnetworks indicate that ECM regulation fundamentally differs between lineages. It is previously known that biosynthesis of the follicular BM, which conceptually corresponds to renewal of the laminin^+^ BM we found here initially enclosing the entire lobe rudiment, is coordinated with vascularization whereby endothelial cells actively contribute to both folliculogenesis and recruitment of differentiated C-cells [103, 104]. Laminin-mediated polarization of thyrocytes, of fundamental importance to form follicles, is nonetheless intrinsically regulated by Pax8 [43]. The Ubb-GRN did not predict any negative regulation of collagen IV or laminin genes arguing that the Ubb-BM disappears at least partly by a proteolytic mechanism.

We did not particularly look for stem cell characteristics in the present work. However, cells in cluster 9 sorted out with some shared or intermediate expression patterns that might constitute a small pool of progenitors committed to differentiate into both lineages. Alternatively, this cluster consists of heterogeneous cells with distinct lineage identity although sharing transcriptional profiles sufficient to co-cluster. The possible existence of stem cells acquiring alternative thyroid cell fates rests on in vivo observations of Tg-positive follicles in the ultimobranchial body remnant and C-cell development in ectopic thyroids where direct contact between primordia predestined to either lineage is physically impossible ([3] and refs therein). A side-population of cells with a suggested dual lineage potential has been identified in adult mouse thyroid [105, 106]. Presence of cells fated to share some thyroid and Ubb lineage features might putatively explain occurrence of a second type of follicles of ultimobranchial origin in the thyroid gland [107–110] and rare thyroid tumors with mixed phenotypes in humans [3, 111, 112], as the one studied here. Whether MMFTC might develop from bipotent ancestral stem/progenitor cells, at difference with interfacing collision MTC/FTC tumors that display distinct mutation profiles [113], is yet an unproven concept.

In all vertebrates except mammals, plasma Ca^2+^ levels are tightly controlled by two major calcium-regulating endocrine cell types confined to the parathyroid and ultimobranchial glands, the latter principally constituting free Ubbs independent of any physical or functional connections to the thyroid gland of the same species. The physiological role of calcitonin in humans has been controversial [114–116], fueled by the fact that calcitonin is not required to substitute after thyroidectomy. Indeed, ultimobranchial calcitonin is arguably more important for systemic calcium metabolism in aquatic vertebrates as amphibians [117, 118]. However, there is also ample evidence in the literature indicating that calcitonin secreted by thyroid C-cells facilitates net transfer of calcium to bone in response to postprandial hypercalcemia thereby suppressing bone resorption. Using skeleton, the primary tissue target of circulating calcitonin, as a buffer for maintaining calcium homeostasis probably was evolutionarily advantageous at the transition from aquatic to terrestrial life [119]. It is nonetheless difficult to find a rationale why mammalian C-cells do not stay settled in the Ubb but integrate with thyroid cells if were it not for the benefit of coordinated regulation of hormone secretion, as suggested by the pioneering work of Eladio Nunez and coworkers on thyroid alterations in hibernating bats [120]. Although today receiving poor attention among scientists and endocrinologists, there are several independent reports in support of intrathyroidal regulation by reciprocal paracrine interactions [121–125]. The present study reveals embryonic thyroid-Ubb fusion is a tightly regulated process that does not happen by chance but involves both lineages of thyroid endocrine cell types intricately.

## Methods

### Mouse strain breeding, collection of embryos and experiments

Wildtype mouse embryos, generated by breeding of C57BL/6J mice housed at the animal core facility Experimental Biomedicine (EBM), University of Gothenburg, were dissected at E9.5-E18.5, where the morning of plug detection was considered E0.5, and saved for preparation of frozen sections and immunofluorescence analysis. As previously described [14], the scRNA-seq pharyngeal atlas was generated from heterozygous *Pax9^VENUS^* embryos obtained by crossing *Pax9^VENUS^*reporter male mice with C57BL/6J females at the animal facility of the Massachusetts Chan Medical School. *Nkx2-1^+/-^* and *Pax8^-/-^* embryos were generated by breeding of null heterozygous mice of C57BL/6J background available at the animal facility of University of Naples Federico II and collected at E15.5 for the same purpose. Heterozygous *Nkx2.1-CreERT2* mice carrying the *Nkx2.1-CreER* knock-in/knock-out allele (RRID:IMSR_JAX:014552; The Jackson Laboratory), which disrupts one *Nkx2-1* allele [38], were alternatively used to investigate the Nkx2-1 heterozygous phenotype. Serving as controls to induced activation of an irrelevant floxed allele, both *Nkx2.1-CreERT2^+/-^* and age-matched wildtype mice were injected with tamoxifen intraperitoneally (10 mg/ml; 50 µl) daily x4 ten days before sacrifice. *Tg-CreERT2;Braf^CA/+^* mutants were generated by recombination of heterozygous *Tg-CreERT2* in which the *thyroglobulin* (*Tg*) promoter acts as Cre driver (RRID:IMSR_JAX:030676; Jackson) and *Braf^CA^* mice in which the floxed human Braf oncogene encodes BRAF^V600E^ (RRID:IMSR_JAX:017837; Jackson). Non-induced *Tg-CreERT2;Braf^CA/+^* mice represent a model of sporadic thyroid cancer characterized by multifocal development of papillary thyroid carcinoma due to spontaneous, stochastic activation of Cre recombinase [46]. Oral PLX4720 (417 ppm; provided by Plexxikon), a prodrug to vemurafenib, and control dietary pellets were continuously supplied during treatment period, as indicated. In general, strains were backcrossed with C57BL/6J female mice (B6-F; Taconic Biosciences) 10 times before recombination. Ear punch biopsies were sampled for genotyping with PCR. Animal experiments were approved by: the regional ethical committee (Approval No 26-2013 and 5.8.18-03925/2018) according to European standards and national regulations provided by the Swedish Agriculture Agency; the local ethical committee “Comitato Etico per la Sperimentazione Animale” (CESA) in accordance with the regulations and guidelines of Italy and the European Union; the Institutional Animal Care and Use Committee (Approval protocol #2384) according to the regulatory standards defined by the National Institutes of Health and the University of Massachusetts.

### Human tumor samples

Fresh thyroid tumor samples subjected to immunohistochemical staining of FFPE tissue sections were obtained at neck surgery of a 69 years-old female with a neuroendocrine metastatic malignancy previously included in an approved patient-derived xenograft study [126]. The likelihood of a primary thyroid tumor origin is inferred by histopathological diagnosis of a *Ret* mutation negative mixed medullary-follicular thyroid carcinoma (MMFTC). As revealed by immunohistochemistry, the neuroendocrine tumor was positive for Nkx2-1/TTF1, chromogranin A and synaptophysin but negative of calcitonin, which was also not detected in serum. Radiology confirmed widespread metastases in lungs, mediastinum and skeleton.

### Computational analysis of datasets with thyroid and Ubb lineage identities from single-cell RNA-seq and ATAC-seq pharyngeal atlas

All information on general procedures including cell isolation, library preparation, sequencing, alignment, quantification, quality control and filtering, and the final data processing and global analysis of single-cell transcriptomes and chromatin landscapes obtained from *Pax9^+^* mouse pharyngeal endoderm are previously published [14] and deposited resources available in the Gene Expression Omnibus (GEO) database under accession codes GSE182135 (scRNA atlas) and GS182134 (scATAC atlas).

### Single-cell RNAseq data analysis

Analysis was performed using scanpy 1.8.2 [127]. Single-cell RNA (scRNA) counts from cells in clusters 12, 13, 17, and 22 of the scRNA atlas (Supplementary Fig. 1a; Magaletta et al [14]) were extracted leaving 5904 cells from 10 samples across 4 Edays as follows – 3121 from E9.5, 912 from E10.5, 1186 from E11.5, and 685 from E12.5. This dataset had been previously filtered for contaminants including cells with neuronal and neural crest signatures, as described [14]. Genes occurring in less than 2 cells were removed leaving 18081 genes. Counts per million normalized counts were log transformed. The top 30 principal components were used to obtain a nearest neighbor graph and perform Louvain clustering at resolution = 0.5. Size factors were computed using scran’s [128] computeSumFactors with min.mean = 0.1 and using the Louvain clusters. Cell raw counts were normalized using these size factors and log-transformed using scanpy.pp.log1p. Sample batch correction was performed in scanpy using combat with default parameters. Cell cycle phase scores and predicted phase for each cell were computed using scanpy’s score_genes_cell_cycle with lists of genes in S and G2M phases from Table S2 in: [129]. 2450 highly variable genes were obtained after excluding mitochondrial, ribosomal protein (Rpl and Rps), and mitochondrial ribosomal protein (Mrpl and Mrps) genes, using scanpy’s highly_variable_genes with default parameters. Correction for counts, mitochondrial content, and phase was performed using scanpy’s regress_out. The top 35 principal components were used to obtain a nearest neighbor graph. Leiden clustering at resolution 1.2 gave 10 clusters labelled 0-9 in decreasing order of number of cells in the cluster.

Cells were visualized on a UMAP embedding obtained with scanpy.tl.umap and default parameters. UMAPs were visualized with scanpy.pl.umap and dotplots with scanpy.pl.dotplot.

Normalized and scaled number of cells in each category were visualized with stacked barplots using pandas.plot with “kind=bar” and stacked=True.

Differential expression analysis was performed using scanpy’s rank_genes_groups with adjusted p-value < 0.01, log2-fold change > 1.0 and otherwise default parameters. For the heatmap of cluster specific genes, besides the p-value and log2foldchange cutoff, additional constraints were set, namely that the gene must be expressed in > 20% of the cells in that cluster. Further, a specificity score for each gene in a cluster was computed as the ratio of the total normalized expression in that cluster by the total normalized expression in the dataset. For each cluster, genes with specificity in the top 40 percentile of that cluster were retained. Genes with the top 10 z-scores from scanpy rank_gene_groups scores were visualized on the heatmap in scanpy with clusters ordered using hierarchical clustering with scanpy’s dendrogram. These z-scores were also used to discuss gene enrichment in the results.

For the differential expression analysis between *Calca* positive and negative cells in the ubb at E12.5, cells in clusters 4, 6 and 8 at E12.5 were used. Cells with normalized expression of *Calca* > 0 were labelled *Calca* positive, and the rest *Calca* negative. For the differential analysis between thyroid and Ubb, cells in clusters 4, 6 and 8 were used for the Ubb, and cells in clusters 1, 2 and 3 for the thyroid. The p-value cutoff was 0.01, log2foldchange cutoff was 1.0.

Enrichment analysis of differentially expressed genes, obtained using scanpy’s rank_genes_groups with adjusted p-value < 0.01, log2-fold change > 1.0 and otherwise default parameters, was performed using gget enrichr [130–133] with adjusted p-value < 0.1. A list of terms was curated from the resulting enriched terms and the negative log binomial FDR value was plotted on a scaled heatmap generated using scanpy.pl.heatmap with standard_scale=’var’.

To obtain scores associated with specific terms, gene lists associated with the corresponding Gene Ontology (GO) [134, 135] terms were obtained from the Gene Ontology Browser at Mouse Genome Informatics, The Jackson Laboratory, Bar Harbor, Maine. World Wide Web (URL: http://www.informatics.jax.org). scanpy.tl.score_genes was used to score them and the resulting per-cell scores were visualized on UMAP embeddings.

Venn diagrams of genes in all 9 clusters were generated using BioVenn [136]. A compound diagram mirroring the topography of the unbiased reclustering of UMAP cluster 12, 13, 17 and 22 from Magaletta et al [14]was manually assembled.

### Pseudotime and trajectory inference

To visualize developmental trajectories, we generate a force directed embedding by stitching adjacent timepoints together with Harmony [21]. Next, we use CellRank 2 [137] to perform pseudotime and trajectory inference with two complementary approaches – Palantir [22] and moscot [24, 25].

#### Integrating timepoints with Harmony

Starting from the raw UMI counts, perform normalization using scanpy normalize_total with total_counts = 10000 and the resulting log-transformed pseudo-counts were obtained using the scanpy log1p function. For feature selection, ribosomal (Rps and Rpl), mitochondrial(mt-), mitochondrial-ribosomal (Mtrpl and Mtrps) transcripts were excluded. Genes associated with cell cycle from Table S8 in Park et al. [138] were also excluded and the highly variable genes were obtained using scanpy.highly_variable_genes with min_mean=0.0125, max_mean=3, and min_disp=0.5. The top 100 principal components were computed on this set of highly variable genes using scanpy.pca with svd-solver=’arpack’.. Harmony from Nowotschin et al [21] was used to obtain an augmented affinity matrix with additional connections between consecutive time-points. Random number generators were seeded for repeatability (np.random.seed(0)). The augmented affinity matrix was used as input for a force-directed graph layout using ‘harmony.plot.force_directed_layout’.

#### Pseudotime inference with Palantir

Using Palantir [22], run_diffusion_maps was run on the augmented affinity matrix from above to obtain the diffusion map which was used as input to palantir.utils.determine_multiscale_space to get the multi-scale distance. Since the force directed embedding in Fig. 2a was constructed taking the embryonic day into account with Harmony as described above, we used it to define an early cell (from which Palantir selects a start state) and terminal states for Palantir. Early cell was defined as the cell with the lowest Y-coordinate. Two terminal states were defined from among E12.5 cells belonging to cluster 3 (thyroid) and cluster 4 (Ubb) and having the largest X-coordinate in Fig. 2a. Palantir was run with palantir.core.run_palantir, early cell and terminal states and otherwise default parameters to obtain pseudotime and fate probabilities in Fig. 2a, c; palantir.utils.run_magic_imputation was used to impute gene expression with MAGIC [139] on the diffusion map obtained as described above; palantir.presults.select_branch_cells was run to identify cells most likely to follow each of the two lineages. The MAGIC imputed expression was plotted for key markers with palantir.presults.compute_gene_trends. Lineage drivers were inferred with compute_lineage_drivers from CellRank 2 [137]. Heyl+ cells were defined as those with scran normalized expression of *Heyl* > 0. Since lack of expression of *Heyl* could be the effect of dropout, gene expression imputation was performed with MAGIC and cells with *Heyl* < 0.02 (chosen with a cutoff at the lowest mode of the imputed expression distribution) were counted as Heyl-. The Heyl+/- subset of cells in clusters 1, 2, and 3 were retained with a total of 882 Heyl- and 150 Heyl+ cells.

#### Pseudotime inference with moscot

We adopted a complementary strategy to infer cell trajectories. Ribosomal (Rps and Rpl), mitochondrial(mt-), mitochondrial-ribosomal (Mtrpl and Mtrps) transcripts and genes associated with cell cycle from Table S8 in Park et al. [138] were excluded. Principal components and nearest neighbors were identified with scanpy.pp.pca and scanpy.pp.neighbors with random_state = 0. moscot.TemporalProblem was run to set up the problem. Next, marginals were adjusted to account for cellular growth and death with score_genes_for_maginals with mouse gene sets for proliferation and apoptosis. The OT problem was prepared and solved with prepare (time_key=eday) and solve (epsilon=1e-3, tau_a=0.95, scale_cost="mean"). epsilon and tau_a control the amount of entropic regularization (speeding up and stabilizing the solution) and unbalancedness on the source marginal (accounting for cell growth and death), respectively. Next, CellRank 2 [137] RealTimeKernel was set up with RealTimeKernel.from_moscot and the transition matrix computed with compute_transition_matrix(self_transitions="all", conn_weight=0.2, threshold="auto"). To automatically identify initial and terminal states, we initialize the Generalized Perron Cluster Cluster Analysis (GPCCA) estimator in CellRank 2 with the RealTimeKernel from above. We fit two states with fit(), predict terminal states with predict_terminal_states() and plot them with plot_macrostates(which=”terminal”).

### GRN inference from single-cell RNA-seq and ATAC-seq pharyngeal atlas

The CellOracle [25] inferred GRNs for each of the clusters in Magaletta et al [14] (Supplementary Fig. 1a) were used. The inference apporach is restated here for clarity. Base GRN construction Peaks and peak-to-peak co-accessibility were obtained in ArchR. A peak was associated with a target gene if it overlapped with the TSS (TSS peak, co-accessibility = 1) or if it had a co-accessibility ≥0.5 with a TSS peak. Peaks were scanned for motifs using CellOracle’s scan function which uses the gimmemotifs motif scanner [140] (background_length = 200, fpr = 0.02, default motif database = gimme.vertebrate.v5.0 using binding and inferred motifs and cumulative binding score cutoff = 10) to generate an annotated peak-motif binary matrix which is the base GRN in CellOracle.

Using CellOracle, the base GRN was refined using the atlas scRNA data to form cell-type-specific GRN’s and simulate the TF knockouts *in silico*. 10,000 genes were selected by requiring at least 3 counts and by using scanpy’s preprocessing utility ‘scanpy.pp.filter_genes_dispersion’ with ‘flavor = ‘cell_ranger’’ and ‘n_top_genes = 10,000’. The selected genes included all 1119 variable genes used in the atlas scRNA analysis. Normalized (using sc.pp.normalize_per_cell) count data was imputed in a 50 principal components subspace using CellOracle’s balancedKNN implementation (k = 54 nearest neighbors, b_sight = 54*8, b_maxl = 54*4). Cell-type-specific GRN’s were trained using CellOracle’s default procedure: for each target gene and each cell type, bagging ridge regression was run (bagging_number = 20, alpha = 10) using connections determined by the base GRN. Edges were preserved with edge p-value ≤ 0.001 where the source node was in the top differentially expressed genes of that cluster as determined by Seurat’s FindAllMarkers (log2-fold change threshold = 0.25, return.thresh = 0.01) on the scRNA atlas and the edge weight was >0.005 leaving us with a trained GRN. Network statistics including betweenness centrality were computed for each GRN using the CellOracle get_score function. GRNs were re-trained on the preserved edges using the same parameters as before. The resulting TF-target gene links, average cell-type specific GRN bagging ridge model edge weights (“coef_mean”) and associated p-values, were extracted from the “filtered_links” attribute in the CellOracle object.

### *In silico* knockout simulations using GRNs

CellOracle simulateshift was used for the knockout simulations of *Nkx2-1*, *Pax8*, *Foxa1* and *Foxa2*, respectively. In brief, for each of the GRNs containing the respective TF, the TF expression was set to 0 and propagated through the network up to a depth of 5. To investigate the shift in gene expression in the presence of the *in silico* knockout simulations on the data subset used in this study, we predicted and then visualized the direction and magnitude of the simulated knockout using stream plots in scVelo 0.2.5 [26]. The difference of simulated knockout and imputed counts (delta_X in CellOracle) was defined as velocity and the imputed counts obtained according to the CellOracle pipeline in the original study were defined as the spliced moment Ms in scvelo. Either all cells in the scRNAseq atlas from Magaletta et al were used ([14]: Figs. 2g and h, left panels) or the cells were subset to the ubb and thyroid lineages as defined in this study (Figs. 2g and h, right panels; Fig. 7i). For each of the cases, the transition probabilities were computed on the corresponding cells using scvelo velocity_graph function with default parameters. Magnitude of the velocity, which in this context represents the per cell magnitude of the change in expression between the cells before and after the *in silico* knockout simulation, is obtained from the ‘velocity_length’ field after running scvelo velocity_confidence with default parameters. The velocities are projected onto the scRNA atlas UMAP embedding using scvelo velocity_embedding_stream function with lower velocities filtered out (min_mass = 3). The per-cell change in expression (“velocity_length”) is visualized on the UMAP embedding with scvelo pl.scatter.

### Orthologous comparison of transcriptomes

The enriched transcripts in cluster 0-9 or intersects of select clusters (data set A) were compared to transcripts previously found to be enriched (fold change > 3) in the mouse E10.5 thyroid bud by microarray (Affymetrix Mouse 430 2.0 Genome Array) analysis of micro-dissected midline primordia (data set B) [17]. Comparisons were performed in two ways. Firstly, it was investigated which of the 50 transcripts in each cluster (0-9) with the highest score within data set A that were present within the entire set of enriched transcripts in data set B. Genes absent from the Mouse 430 2.0 Genome Array were excluded from calculations. The median fold change of concordant transcripts retrieved from data set B was calculated. Secondly, it was investigated which of the 50 transcripts in data set B with the highest fold change that were present within the clusters or intersects of data set B.

### *In situ* hybridization

Spatial detection of target RNA expression within intact tissues was performed on formalin-fixed paraffine embedded (FFPE) specimens (both embryonic and adult) or frozen sections (embryonic only) using RNAscope assays according to manufacturer’s instruction (Advanced Cell Diagnostics/ACD). Excised embryos were fixed in 4 % formaldehyde for 24 h, processed for FFPE or transferred to 30 % sucrose in phosphate buffered saline (PBS) overnight, embedded in OCT Tissue-Tek (Sakura) and frozen at -80 °C. Transverse serial sections (thickness 10 µm) of the neck including regions of interest (thyroid primordium and Ubb) were cut on a microtome (FFPE) or a Leica cryostat. Sections collected on SuperFrost Plus® Slides (Fisher Scientific; Cat. No. 1255015) were hybridized at 40°C for 2 h with the following RNAscope™ Probes from ACD (Catalog number-Channel): Mm-Nkx2.1 (434721-C1 and 434721-C2), Mm-Pax8 (574431-C1 and 574431-C2), Mm-Foxa1 (492431-C1), Mm-Foxa2 (409111-C1), Mm-Heyl (446881-C1), Mm-Prdm1 (441871-C1), Mm_Tcf4 (423691-C1), Mm-Pmp22 (505171-C2), Mm-Pecam1 (316721-C2) Mm-Calca (578771-C1), Mm-Col1a2 (585461-C1), Mm-Col4a2 (544351-C1), Mm-Cdh2 (489571-C1) and as negative control DapB (310043). Hybridization signals were amplified and visualized with RNAscope 2.5 HD Detection Kit RED (322360) or RNAscope™ Multiplex Fluorescent detection Reagent Kit v2 (323110). Digital whole slides were obtained using a wide-field Zeiss Axioskop2 plus microscope equipped with a Nikon DS-Qi1Mc camera used for imaging.

### Immunofluorescence and immunohistochemistry

Embryos were fixed and handled as detailed above for ISH. Serial sections collected on parallel poly-L-lysine coated glass slides (Vector Laboratories) for multiple immunostainings of the same specimens were permeabilized in Triton X-100 (0.1 %) in PBS for 20 min, washed in PBS for 5 min, and incubated for one hour with normal donkey serum (2 %) (Jackson Immunoresearch) dissolved in PBS. Sections were then double-labelled overnight in 4 °C with primary antibodies (Catalog number/dilution): rabbit anti-NKX2-1/TTF-1 (PA0100/1:1000; Biopat; ab227652/1:100; Abcam), rabbit anti-PAX8 (No 10336-1-AP/1:2000; Proteintech Europe), guinea pig anti-FOXA1 (kindly provided by Jeffrey Whitsett, Cincinnati Children’s Hospital, OH, USA; 1:2000; applied for mouse tissue only), mouse anti-FOXA1 (No WMAB-2F83/1:1000; Seven Hills Bioreagents; applied for human specimens only), rabbit anti-FOXA2 (WRAB-FOXA2/1:2000; Seven Hills), rat anti-E-cadherin/CDH1 (13-1900/1:4000; Novex/Life Technologies), rabbit anti-N-cadherin/CDH2 (ab18203/1:500 for IHC, 1:200 for IF; Abcam), rabbit anti-laminin/LAM (L9393/1:500; Sigma-Aldrich), chicken anti-laminin/LAM (ab14055/1:500; Abcam), rabbit anti-COL4A1 (ab6586/1:400; Abcam), rat anti-CD31 (550274/1:250; Pharmingen), rabbit anti-thyroglobulin/TG (A0251/1:5000; Agilent), rabbit anti-calcitonin/CALC (102480/1:500; Agilent), rabbit anti-pericentrin/PCNT (ab4448/1:1000; Abcam), rabbit anti-Ki67 (ab15580/1:500; Abcam), and Armenian hamster anti-mucin1/MUC1 (CT2 monoclonal against aa 239-255 (SSLSYTNPAVAATSANL) of the cytoplasmic tail of MUC1, 1:1000; gift from Cathy Madsen at Sandra Gendler Lab, Mayo Clinic; also available at ThermoFischer MA5-11202). The two antibodies against LAMININ were tested by double labeling of the same sections and showed identical staining. Secondary Rhodamin Red^TM^-X-conjugated donkey-anti-rabbit IgG (Jackson Immunoresearch) and biotin-conjugated donkey-anti-rat or anti-chicken IgGs (Jackson Immunoresearch) were applied for one hour followed by Streptavidin-FITC (Agilent) for 30 min at 4 °C. CT2 staining used a FITC-conjugated goat anti-Armenian hamster-IgG (1:200; Jackson Immunoresearch). All incubation steps were followed by washing 3x5 min in Triton X-100 (0.1 %) in PBS and finally with PBS only. Sections were incubated with DAPI nuclear stain (Sigma-Aldrich) and mounted with cover glass with fluorescence mounting medium (Agilent). A wide-field Zeiss Axioskop2 plus microscope equipped with a Nikon DS-Qi1Mc camera was used for imaging. The NIS Elements Imaging Software was used for image processing.

FFPE tissue samples (adult mouse thyroid resected en bloc; human thyroid tumor tissues freshly obtained at surgery) were subjected to immunohistochemical labeling of deparaffinized sections with cross-reacting antibodies against (sources indicated above): thyroglobulin, calcitonin, Foxa1, Foxa2, Cdh1, Cdh2 and collagen type IVa1. PT Link (Agilent) was used for antigen retrieval and quenching of endogenous peroxidase activity prior to immunostaining optimized with the Dako EnVision system (Agilent) or the ImmPress system (VectorLabs) depending on primary antibody.

Sections were viewed and imaged in an Olympus BX45TF microscope equipped with a Nikon DS-U2 camera.

### Western blot analysis

Lysates from subconfluent TT cells cultured according to manufacturer’s instructions (CRL-1803^TM^; ATCC, Manassas, VA) and wildtype and Braf mutant mice were extracted using RIPA buffer containing protease. Protein concentrations were quantified using a BCA protein assay kit (ThermoFisher Scientific). Lysate proteins were separated by 6–15% sodium dodecyl sulfate–polyacrylamide gel electrophoresis and transferred to a polyvinylidene difluoride membrane, followed by blocking with 5% skim milk for 1 h. Membranes were incubated overnight at 4 °C with antibodies (catalog number/dilution) against: Cdh1 (610181/1:5000; BD Transduction Laboratories), Cdh2 (ab18203/1:1000; Abcam), Foxa1 (WMAB-2F83/1:1000; Seven Hills), Foxa2 (WRAB-FOXA2/1: 2000; Seven Hills), calcitonin (A0576/1: 500; DAKO), Heyl (15679-1-AP/1:1000; ThermoFisher) and β-actin (P07437/1:1000; Abways). Labeled membranes were incubated with the appropriate secondary antibodies (anti-rabbit: 31460; or anti-mouse: 31430 (both Invitrogen) for 1 h at room temperature. Specific bands were detected by enhanced chemiluminescence. Protein levels were analyzed by densitometry; representative data are mean ± SEM from n=3 per group. *p < 0.05, **p < 0.01, ***p< 0.001. Supplementary Dataset 4 and 5 provide the uncropped blots from all included experiments.

### Morphometry

Neck serial sections of wildtype embryos collected at E12.5 and E13 and *Pax8^-/-^* and *Nkx2.1^+/-^* mutant embryos both collected at E15.5 (*n* for each timepoint and genotype is indicated in legend to Fig. 6) were alternatingly immunostained for NKX2-1/CDH1 and LAM/CDH1 and subjected to morphometric measurements of (bilateral) Ubb total size, Ubb lumen size and and percentual laminin coating of the Ubb surface on images captured with the Zeiss Axioscope 2 Plus fluorescence microscope equipped with a Nikon DS-Qi1Mc camera. Statistical analysis of morphometric data was performed using student’s *t* test with a *p*-value <0.05 considered significant.

### Transmission electron microscopy

Mouse embryos of age E11.5, E12.5, E13.0 and E13.5 were fixed with 2 % formaldehyde and 2.5 % glutaraldehyde dissolved in 0.05 M sodium cacodylate buffer containing 0.02 % sodium azide for 24 h. An oscillation tissue slicer was used to produce, from the embryo’s neck, 150 µm transverse sections that were postfixed in 1 % osmium tetroxide with 1 % potassium hexacyanoferrate for 2 h at 4°C and thereafter contrasted *en bloc* with 0.5 % uranyl acetate for 1 h in the dark at room temperature. After stepwise dehydration in alcohol series and acetone, specimens were embedded in Agar 100 Resin according to standard procedures. Sections of 1 µm were cut on a Leica EM UC6 ultramicrotome and stained with azure methylene blue for light microscopic identification of the thyroid primordium and Ubb; selected sections were subjected to ultrathin sectioning. Sections with a thickness of approximately 60 nm were contrasted with uranyl acetate and lead citrate, coated with carbon by evaporation and examined in a LEO 912AB Omega transmission electron microscope.

## Acknowledgements

This work was supported by grants from the Swedish Cancer Society (22-2426 and 25-4799 to MN), the Swedish Research Council (2016-02360 to MN; 2017-06278 to CG), the US-Israel Binational Science Foundation (to RM), the University of Pennsylvania Orphan Disease Center in partnership with the Hypopara Research Foundation (to RM), the Gothenburg Medical Society (to ES and JD), Assar Gabrielsson Research Foundation (to ES), the Swedish state under the ALF-agreement between the Swedish government and the county councils (associated with clinical research position of HF).

## Author Contributions

MN conceptualized the study and wrote the manuscript draft. ML, EJ, SK, CG, HF, RM and MN edited the manuscript. ML, EJ, SK, CG, HF, RM and MN designed, analyzed and interpreted the experiments. ML, EJ, SK, ES, IA, SL, TC, BRJ, PM, MDF and JD collected tissues and performed experiments. ML performed all bioinformatic data processing and analyses.

## Materials & Correspondence

Correspondence and material requests should be addressed to

## Data availability

This paper analyzes existing publicly available data in the Gene Expression Omnibus (GEO) database under accession codes “GSE182135” (scRNA atlas) and “GSE182134” (scATAC atlas).

## Code availability

Code used for scRNAseq/bioinformatic analysis will be released at https://github.com/maehrlab/ThyroidUBB_Lineage upon acceptance of the manuscript.

For the purpose of review, code to reproduce analyses and figures is available on Dropbox at https://www.dropbox.com/scl/fo/qsdz34ayu11s2ojy827a9/AIMOQzrQW_ZvaV7O4nx-qfI?rlkey=y6171v7th61wvh8o26f1gm1tk&st=k5z7pjg5&dl=0

## References

1. Nilsson, M. and H. Fagman, Development of the thyroid gland. Development, 2017. 144(12): p. 2123–2140.

2. Kameda, Y., Morphological and molecular evolution of the ultimobranchial gland of nonmammalian vertebrates, with special reference to the chicken C cells. Dev Dyn, 2017. 246(10): p. 719–739.

3. Nilsson, M. and D. Williams, On the Origin of Cells and Derivation of Thyroid Cancer: C Cell Story Revisited. Eur Thyroid J, 2016. 5(2): p. 79–93.

4. Adams, M.S. and M. Bronner-Fraser, Review: the role of neural crest cells in the endocrine system. Endocr Pathol, 2009. 20(2): p. 92–100.

5. Johansson, E., et al., Revising the embryonic origin of thyroid C cells in mice and humans. Development, 2015. 142(20): p. 3519–28.

6. Kameda, Y., Cellular and molecular events on the development of mammalian thyroid C cells. Dev Dyn, 2016. 245(3): p. 323–41.

7. Peters, H., et al., Pax9-deficient mice lack pharyngeal pouch derivatives and teeth and exhibit craniofacial and limb abnormalities. Genes Dev, 1998. 12(17): p. 2735–47.

8. Manley, N.R. and M.R. Capecchi, The role of Hoxa-3 in mouse thymus and thyroid development. Development, 1995. 121(7): p. 1989–2003.

9. Liao, J., et al., Full spectrum of malformations in velo-cardio-facial syndrome/DiGeorge syndrome mouse models by altering Tbx1 dosage. Hum Mol Genet, 2004. 13(15): p. 1577–85.

10. Fagman, H., et al., The 22q11 deletion syndrome candidate gene Tbx1 determines thyroid size and positioning. Hum Mol Genet, 2007. 16(3): p. 276–85.

11. Kusakabe, T., N. Hoshi, and S. Kimura, Origin of the ultimobranchial body cyst: T/ebp/Nkx2.1 expression is required for development and fusion of the ultimobranchial body to the thyroid. Dev Dyn, 2006. 235(5): p. 1300–9.

12. Krude, H., et al., Choreoathetosis, hypothyroidism, and pulmonary alterations due to human NKX2-1 haploinsufficiency. J Clin Invest, 2002. 109(4): p. 475–80.

13. Mio, C., F. Baldan, and G. Damante, NK2 homeobox gene cluster: Functions and roles in human diseases. Genes Dis, 2023. 10(5): p. 2038–2048.

14. Magaletta, M.E., et al., Integration of single-cell transcriptomes and chromatin landscapes reveals regulatory programs driving pharyngeal organ development. Nat Commun, 2022. 13(1): p. 457.

15. Liang, S., et al., A branching morphogenesis program governs embryonic growth of the thyroid gland. Development, 2018. 145(2).

16. Mansouri, A., K. Chowdhury, and P. Gruss, Follicular cells of the thyroid gland require Pax8 gene function. Nat Genet, 1998. 19(1): p. 87–90.

17. Fagman, H., et al., Gene expression profiling at early organogenesis reveals both common and diverse mechanisms in foregut patterning. Dev Biol, 2011. 359(2): p. 163–75.

18. Haerlingen, B., et al., Mesodermal FGF and BMP govern the sequential stages of zebrafish thyroid specification. Development, 2023. 150(10).

19. Okubo, T., et al., Ripply3, a Tbx1 repressor, is required for development of the pharyngeal apparatus and its derivatives in mice. Development, 2011. 138(2): p. 339–48.

20. Shone, V., et al., Mode of reduction in the number of pharyngeal segments within the sarcopterygians. Zoological Lett, 2016. 2: p. 6.

21. Nowotschin, S., et al., The emergent landscape of the mouse gut endoderm at single-cell resolution. Nature, 2019. 569(7756): p. 361-367.

22. Setty, M., et al., Characterization of cell fate probabilities in single-cell data with Palantir. Nat Biotechnol, 2019. 37(4): p. 451–460.

23. Lange, M., et al., CellRank for directed single-cell fate mapping. Nat Methods, 2022. 19(2): p. 159–170.

24. Schiebinger, G., et al., Optimal-Transport Analysis of Single-Cell Gene Expression Identifies Developmental Trajectories in Reprogramming. Cell, 2019. 176(4): p. 928–943 e22.

25. Klein, D., et al., Mapping cells through time and space with moscot. Nature, 2025. 638(>8052): p. 1065–1075.

26. Lanigan, T.M., S.K. DeRaad, and A.F. Russo, Requirement of the MASH-1 transcription factor for neuroendocrine differentiation of thyroid C cells. J Neurobiol, 1998. 34(2): p. 126–34.

27. Kameda, Y., et al., Mash1 regulates the development of C cells in mouse thyroid glands. Dev Dyn, 2007. 236(1): p. 262–70.

28. Ciampi, R., et al., Genetic Landscape of Somatic Mutations in a Large Cohort of Sporadic Medullary Thyroid Carcinomas Studied by Next-Generation Targeted Sequencing. iScience, 2019. 20: p. 324–336.

29. Kamimoto, K., et al., Dissecting cell identity via network inference and in silico gene perturbation. Nature, 2023. 614(7949): p. 742-751.

30. Bergen, V., et al., Generalizing RNA velocity to transient cell states through dynamical modeling. Nat Biotechnol, 2020. 38(12): p. 1408–1414.

31. Bouchard, M., et al., Nephric lineage specification by Pax2 and Pax8. Genes Dev, 2002. 16(22): p. 2958–70.

32. Batista, M.F. and K.E. Lewis, Pax2/8 act redundantly to specify glycinergic and GABAergic fates of multiple spinal interneurons. Dev Biol, 2008. 323(1): p. 88–97.

33. Bouchard, M., et al., Pax2 and Pax8 cooperate in mouse inner ear morphogenesis and innervation. BMC Dev Biol, 2010. 10: p. 89.

34. Johansson, E., et al., Asynchrony of Apical Polarization, Luminogenesis, and Functional Differentiation in the Developing Thyroid Gland. Front Endocrinol (Lausanne), 2021. 12: p. 760541.

35. Fernández, L.P., A. López-Márquez, and P. Santisteban, Thyroid transcription factors in development, differentiation and disease. Nat Rev Endocrinol, 2015. 11(1): p. 29–42.

36. Antonica, F., et al., Generation of functional thyroid from embryonic stem cells. Nature, 2012. 491(7422): p. 66–71.

37. Parlato, R., et al., An integrated regulatory network controlling survival and migration in thyroid organogenesis. Dev Biol, 2004. 276(2): p. 464–75.

38. Taniguchi, H., et al., A resource of Cre driver lines for genetic targeting of GABAergic neurons in cerebral cortex. Neuron, 2011. 71(6): p. 995–1013.

39. Kusakabe, T., et al., Thyroid-specific enhancer-binding protein/NKX2.1 is required for the maintenance of ordered architecture and function of the differentiated thyroid. Mol Endocrinol, 2006. 20(8): p. 1796–809.

40. Fagman, H., et al., Expression of classical cadherins in thyroid development: maintenance of an epithelial phenotype throughout organogenesis. Endocrinology, 2003. 144(8): p. 3618–24.

41. Calì, G., et al., CDH16/Ksp-cadherin is expressed in the developing thyroid gland and is strongly down-regulated in thyroid carcinomas. Endocrinology, 2012. 153(1): p. 522–34.

42. de Cristofaro, T., et al., An essential role for Pax8 in the transcriptional regulation of cadherin-16 in thyroid cells. Mol Endocrinol, 2012. 26(1): p. 67–78.

43. Koumarianou, P., G. Goméz-López, and P. Santisteban, Pax8 controls thyroid follicular polarity through cadherin-16. J Cell Sci, 2017. 130(1): p. 219–231.

44. Silberschmidt, D., et al., In vivo role of different domains and of phosphorylation in the transcription factor Nkx2-1. BMC Dev Biol, 2011. 11: p. 9.

45. Nandagopal, N., et al., Dynamic Ligand Discrimination in the Notch Signaling Pathway. Cell, 2018. 172(4): p. 869–880 e19.

46. Schoultz, E., et al., Tissue architecture delineates field cancerization in BRAFV600E-induced tumor development. Dis Model Mech, 2022. 15(2).

47. Traversi, F., et al., BRAF(V600E) Overrides NOTCH Signaling in Thyroid Cancer. Thyroid, 2021. 31(5): p. 787–799.

48. Manley, N.R. and M.R. Capecchi, Hox group 3 paralogs regulate the development and migration of the thymus, thyroid, and parathyroid glands. Dev Biol, 1998. 195(1): p. 1–15.

49. Manley, N.R., et al., Abnormalities of caudal pharyngeal pouch development in Pbx1 knockout mice mimic loss of Hox3 paralogs. Dev Biol, 2004. 276(2): p. 301–12.

50. Wilm, T.P. and L. Solnica-Krezel, Essential roles of a zebrafish prdm1/blimp1 homolog in embryo patterning and organogenesis. Development, 2005. 132(2): p. 393–404.

51. Robertson, E.J., et al., Blimp1 regulates development of the posterior forelimb, caudal pharyngeal arches, heart and sensory vibrissae in mice. Development, 2007. 134(24): p. 4335–45.

52. Fagman, H., L. Andersson, and M. Nilsson, The developing mouse thyroid: embryonic vessel contacts and parenchymal growth pattern during specification, budding, migration, and lobulation. Dev Dyn, 2006. 235(2): p. 444–55.

53. Amici, S.A., et al., Peripheral myelin protein 22 is in complex with alpha6beta4 integrin, and its absence alters the Schwann cell basal lamina. J Neurosci, 2006. 26(4): p. 1179–89.

54. Notterpek, L., et al., Peripheral myelin protein 22 is a constituent of intercellular junctions in epithelia. Proc Natl Acad Sci U S A, 2001. 98(25): p. 14404–9.

55. Roux, K.J., et al., Modulation of epithelial morphology, monolayer permeability, and cell migration by growth arrest specific 3/peripheral myelin protein 22. Mol Biol Cell, 2005. 16(3): p. 1142–51.

56. Baechner, D., et al., Widespread expression of the peripheral myelin protein-22 gene (PMP22) in neural and non-neural tissues during murine development. J Neurosci Res, 1995. 42(6): p. 733–41.

57. O’Brien, L.E., et al., Rac1 orientates epithelial apical polarity through effects on basolateral laminin assembly. Nat Cell Biol, 2001. 3(9): p. 831–8.

58. Pohlenz, J., et al., Partial deficiency of thyroid transcription factor 1 produces predominantly neurological defects in humans and mice. J Clin Invest, 2002. 109(4): p. 469–73.

59. Millership, S.J., et al., Neuronatin regulates pancreatic β cell insulin content and secretion. J Clin Invest, 2018. 128(8): p. 3369–3381.

60. Oczko-Wojciechowska, M., et al., Differences in the transcriptome of medullary thyroid cancer regarding the status and type of RET gene mutations. Sci Rep, 2017. 7: p. 42074.

61. Shen, C., et al., Comprehensive DNA Methylation Profiling of Medullary Thyroid Carcinoma: Molecular Classification, Potential Therapeutic Target, and Classifier System. Clin Cancer Res, 2024. 30(1): p. 127–138.

62. Gianakas, C.A., et al., Hemicentin-mediated type IV collagen assembly strengthens juxtaposed basement membrane linkage. J Cell Biol, 2023. 222(1).

63. Zhang, J.L., et al., Vertebrate extracellular matrix protein hemicentin-1 interacts physically and genetically with basement membrane protein nidogen-2. Matrix Biol, 2022. 112: p. 132–154.

64. Viney, T.J., et al., Regulation of the cell-specific calcitonin/calcitonin gene-related peptide enhancer by USF and the Foxa2 forkhead protein. J Biol Chem, 2004. 279(48): p. 49948–55.

65. Suzuki, K., et al., Thyroid transcription factor 1 is calcium modulated and coordinately regulates genes involved in calcium homeostasis in C cells. Mol Cell Biol, 1998. 18(12): p. 7410–22.

66. Dame, K., et al., Thyroid Progenitors Are Robustly Derived from Embryonic Stem Cells through Transient, Developmental Stage-Specific Overexpression of Nkx2-1. Stem Cell Reports, 2017. 8(2): p. 216–225.

67. Andersson, L., et al., Role of EphA4 receptor signaling in thyroid development: regulation of folliculogenesis and propagation of the C-cell lineage. Endocrinology, 2011. 152(3): p. 1154–64.

68. McKinsey, G.L., et al., Dlx1&2-dependent expression of Zfhx1b (Sip1, Zeb2) regulates the fate switch between cortical and striatal interneurons. Neuron, 2013. 77(1): p. 83–98.

69. Vivinetto, A.L., et al., Zeb2 Is a Regulator of Astrogliosis and Functional Recovery after CNS Injury. Cell Rep, 2020. 31(13): p. 107834.

70. Radice, G.L., N-cadherin-mediated adhesion and signaling from development to disease: lessons from mice. Prog Mol Biol Transl Sci, 2013. 116: p. 263–89.

71. Gheldof, A. and G. Berx, Cadherins and epithelial-to-mesenchymal transition. Prog Mol Biol Transl Sci, 2013. 116: p. 317–36.

72. Papotti, M., et al., Thyroid carcinomas with mixed follicular and C-cell differentiation patterns. Semin Diagn Pathol, 2000. 17(2): p. 109–19.

73. Sadow, P.M. and J.L. Hunt, Mixed Medullary-follicular-derived carcinomas of the thyroid gland. Adv Anat Pathol, 2010. 17(4): p. 282–5.

74. Sobol, R.E., V. Memoli, and L.J. Deftos, Hormone-negative, chromogranin A-positive endocrine tumors. N Engl J Med, 1989. 320(7): p. 444–7.

75. Horejs, C.M., Basement membrane fragments in the context of the epithelial-to-mesenchymal transition. Eur J Cell Biol, 2016. 95(11): p. 427–440.

76. Horejs, C.M., et al., Biologically-active laminin-111 fragment that modulates the epithelial-to-mesenchymal transition in embryonic stem cells. Proc Natl Acad Sci U S A, 2014. 111(16): p. 5908–13.

77. Walter, C., et al., Physical defects in basement membrane-mimicking collagen-IV matrices trigger cellular EMT and invasion. Integr Biol (Camb), 2018. 10(6): p. 342–355.

78. Le Douarin, N., J. Fontaine, and C. Le Lièvre, New studies on the neural crest origin of the avian ultimobranchial glandular cells--interspecific combinations and cytochemical characterization of C cells based on the uptake of biogenic amine precursors. Histochemistry, 1974. 38(4): p. 297–305.

79. Polak, J.M., et al., Immunocytochemical confirmation of the neural crest origin of avian calcitonin-producing cells. Histochemistry, 1974. 40(3): p. 209–14.

80. Rees, J.M., et al., A pre-vertebrate endodermal origin of calcitonin-producing neuroendocrine cells. Development, 2024. 151(20).

81. López-Márquez, A., et al., Sox9 is involved in the thyroid differentiation program and is regulated by crosstalk between TSH, TGFβ and thyroid transcription factors. Sci Rep, 2022. 12(1): p. 2144.

82. Gillotay, P., et al., Single-cell transcriptome analysis reveals thyrocyte diversity in the zebrafish thyroid gland. EMBO Rep, 2020. 21(12): p. e50612.

83. Leimeister, C., et al., Analysis of HeyL expression in wild-type and Notch pathway mutant mouse embryos. Mech Dev, 2000. 98(1-2): p. 175–8.

84. Katoh, M. and M. Katoh, Integrative genomic analyses on HES/HEY family: Notch-independent HES1, HES3 transcription in undifferentiated ES cells, and Notch-dependent HES1, HES5, HEY1, HEY2, HEYL transcription in fetal tissues, adult tissues, or cancer. Int J Oncol, 2007. 31(2): p. 461–6.

85. Han, L., et al., The Notch pathway inhibits TGFbeta signaling in breast cancer through HEYL-mediated crosstalk. Cancer Res, 2014. 74(22): p. 6509–18.

86. Bodas, M., et al., The NOTCH3 Downstream Target HEYL Is Required for Efficient Human Airway Basal Cell Differentiation. Cells, 2021. 10(11).

87. Mosteiro, L., et al., Notch signaling in thyrocytes is essential for adult thyroid function and mammalian homeostasis. Nat Metab, 2023. 5(12): p. 2094–2110.

88. Vincent, S.D., et al., Prdm1 functions in the mesoderm of the second heart field, where it interacts genetically with Tbx1, during outflow tract morphogenesis in the mouse embryo. Hum Mol Genet, 2014. 23(19): p. 5087–101.

89. Romagnoli, M., et al., Epithelial-to-mesenchymal transition induced by TGF-β1 is mediated by Blimp-1-dependent repression of BMP-5. Cancer Res, 2012. 72(23): p. 6268–78.

90. Huang, T.F., et al., BLMP-1/Blimp-1 regulates the spatiotemporal cell migration pattern in C. elegans. PLoS Genet, 2014. 10(6): p. e1004428.

91. Lee, H., et al., Blimp-1 Upregulation by Multiple Ligands. Front Pharmacol, 2022. 13: p. 763678.

92. Menke, A., et al., Down-regulation of E-cadherin gene expression by collagen type I and type III in pancreatic cancer cell lines. Cancer Res, 2001. 61(8): p. 3508–17.

93. Koenig, A., et al., Collagen type I induces disruption of E-cadherin-mediated cell-cell contacts and promotes proliferation of pancreatic carcinoma cells. Cancer Res, 2006. 66(9): p. 4662–71.

94. Shintani, Y., et al., Collagen I promotes metastasis in pancreatic cancer by activating c-Jun NH(2)-terminal kinase 1 and up-regulating N-cadherin expression. Cancer Res, 2006. 66(24): p. 11745–53.

95. Shintani, Y., et al., Collagen I-mediated up-regulation of N-cadherin requires cooperative signals from integrins and discoidin domain receptor 1. J Cell Biol, 2008. 180(6): p. 1277–89.

96. Walsh, L.A., A. Nawshad, and D. Medici, Discoidin domain receptor 2 is a critical regulator of epithelial-mesenchymal transition. Matrix Biol, 2011. 30(4): p. 243–7.

97. Imamichi, Y., et al., Collagen type I-induced Smad-interacting protein 1 expression downregulates E-cadherin in pancreatic cancer. Oncogene, 2007. 26(16): p. 2381–5.

98. DeMaio, L., et al., Ligand-independent transforming growth factor-β type I receptor signalling mediates type I collagen-induced epithelial-mesenchymal transition. J Pathol, 2012. 226(4): p. 633–44.

99. Zhang, K., et al., The collagen receptor discoidin domain receptor 2 stabilizes SNAIL1 to facilitate breast cancer metastasis. Nat Cell Biol, 2013. 15(6): p. 677–87.

100. Xie, B., et al., DDR2 facilitates hepatocellular carcinoma invasion and metastasis via activating ERK signaling and stabilizing SNAIL1. J Exp Clin Cancer Res, 2015. 34(1): p. 101.

101. Koh, M., et al., Discoidin domain receptor 1 is a novel transcriptional target of ZEB1 in breast epithelial cells undergoing H-Ras-induced epithelial to mesenchymal transition. Int J Cancer, 2015. 136(6): p. E508–20.

102. Mitchell, A.V., et al., DDR2 coordinates EMT and metabolic reprogramming as a shared effector of FOXQ1 and SNAI1. Cancer Res Commun, 2022. 2(11): p. 1388–1403.

103. Hick, A.C., et al., Reciprocal epithelial:endothelial paracrine interactions during thyroid development govern follicular organization and C-cells differentiation. Dev Biol, 2013. 381(1): p. 227–40.

104. Villacorte, M., et al., Thyroid follicle development requires Smad1/5- and endothelial cell-dependent basement membrane assembly. Development, 2016. 143(11): p. 1958–70.

105. Hoshi, N., et al., Side population cells in the mouse thyroid exhibit stem/progenitor cell-like characteristics. Endocrinology, 2007. 148(9): p. 4251–8.

106. Murata, T., et al., An Adult Mouse Thyroid Side Population Cell Line that Exhibits Enriched Epithelial-Mesenchymal Transition. Thyroid, 2017. 27(3): p. 460–474.

107. Harach, H.R., Mixed follicles of the human thyroid gland. Acta Anat (Basel), 1987. 129(1): p. 27–30.

108. Wollman, S.H. and P. Nève, Ultimobranchial follicles in the thyroid glands of rats and mice. Recent Prog Horm Res, 1971. 27: p. 213–34.

109. Wollman, S.H. and S.R. Hilfer, Embryologic origin of the various epithelial cell types in the second kind of thyroid follicle in the C3H mouse. Anat Rec, 1978. 191(1): p. 111–21.

110. Kameda, Y., Follicular cell lineage in persistent ultimobranchial remnants of mammals. Cell Tissue Res, 2019. 376(1): p. 1–18.

111. Harach, H.R., Thyroglobulin in human thyroid follicles with acid mucin. J Pathol, 1991. 164(3): p. 261–3.

112. Baloch, Z.W., et al., Overview of the 2022 WHO Classification of Thyroid Neoplasms. Endocr Pathol, 2022. 33(1): p. 27–63.

113. Volante, M., et al., Mixed medullary-follicular thyroid carcinoma. Molecular evidence for a dual origin of tumor components. Am J Pathol, 1999. 155(5): p. 1499–509.

114. Davey, R.A. and D.M. Findlay, Calcitonin: physiology or fantasy? J Bone Miner Res, 2013. 28(5): p. 973–9.

115. Hirsch, P.F., G.E. Lester, and R.V. Talmage, Calcitonin, an enigmatic hormone: does it have a function? J Musculoskelet Neuronal Interact, 2001. 1(4): p. 299–305.

116. Hirsch, P.F. and H. Baruch, Is calcitonin an important physiological substance? Endocrine, 2003. 21(3): p. 201–8.

117. Robertson, D.R., Endocrinology of amphibian ultimobranchial glands. J Exp Zool, 1971. 178(1): p. 101–4.

118. Stiffler, D.F., Amphibian calcium metabolism. J Exp Biol, 1993. 184: p. 47–61.

119. Kurokawa, K., How is plasma calcium held constant? Milieu interieur of calcium. Kidney Int, 1996. 49(6): p. 1760–4.

120. Nunez, E.A. and M.D. Gershon, Cytophysiology of thyroid parafollicular cells. Int Rev Cytol, 1978. 52: p. 1–80.

121. Nunez, E.A. and M.D. Gershon, Thyrotropin-induced thyroidal release of 5-hydroxytryptamine and accompanying ultrastructural changes in parafollicular cells. Endocrinology, 1983. 113(1): p. 309–17.

122. Tamir, H., et al., Expression and development of a functional plasmalemmal 5-hydroxytryptamine transporter by thyroid follicular cells. Endocrinology, 1996. 137(10): p. 4475–86.

123. Ahrén, B., Effects of calcitonin, katacalcin, and calcitonin gene-related peptide on basal and TSH-stimulated thyroid hormone secretion in the mouse. Acta Physiol Scand, 1989. 135(2): p. 133–7.

124. Martín-Lacave, I., et al., C cells evolve at the same rhythm as follicular cells when thyroidal status changes in rats. J Anat, 2009. 214(3): p. 301–9.

125. Morillo-Bernal, J., et al., Functional expression of the thyrotropin receptor in C cells: new insights into their involvement in the hypothalamic-pituitary-thyroid axis. J Anat, 2009. 215(2): p. 150–8.

126. Schoultz, E., et al., Involvement of KEAP1/NRF2 pathway in non-BRAF mutated squamous cell carcinoma of the thyroid. J Pathol, 2025. 266(4-5): p. 481–494.

127. Wolf, F.A., P. Angerer, and F.J. Theis, SCANPY: large-scale single-cell gene expression data analysis. Genome Biol, 2018. 19(1): p. 15.

128. Lun, A.T., D.J. McCarthy, and J.C. Marioni, A step-by-step workflow for low-level analysis of single-cell RNA-seq data with Bioconductor. F1000Res, 2016. 5: p. 2122.

129. Macosko, E.Z., et al., Highly Parallel Genome-wide Expression Profiling of Individual Cells Using Nanoliter Droplets. Cell, 2015. 161(5): p. 1202–1214.

130. Luebbert, L. and L. Pachter, Efficient querying of genomic reference databases with gget. Bioinformatics, 2023. 39(1).

131. Chen, E.Y., et al., Enrichr: interactive and collaborative HTML5 gene list enrichment analysis tool. BMC Bioinformatics, 2013. 14: p. 128.

132. Kuleshov, M.V., et al., Enrichr: a comprehensive gene set enrichment analysis web server 2016 update. Nucleic Acids Res, 2016. 44(W1): p. W90–7.

133. Xie, Z., et al., Gene Set Knowledge Discovery with Enrichr. Curr Protoc, 2021. 1(3): p. e90.

134. Ashburner, M., et al., Gene ontology: tool for the unification of biology. The Gene Ontology Consortium. Nat Genet, 2000. 25(1): p. 25–9.

135. Gene Ontology, C., The Gene Ontology resource: enriching a GOld mine. Nucleic Acids Res, 2021. 49(D1): p. D325–D334.

136. Hulsen, T., J. de Vlieg, and W. Alkema, BioVenn - a web application for the comparison and visualization of biological lists using area-proportional Venn diagrams. BMC Genomics, 2008. 9: p. 488.

137. Weiler, P., et al., CellRank 2: unified fate mapping in multiview single-cell data. Nat Methods, 2024. 21(7): p. 1196–1205.

138. Park, J.E., et al., A cell atlas of human thymic development defines T cell repertoire formation. Science, 2020. 367(6480).

139. van Dijk, D., et al., Recovering Gene Interactions from Single-Cell Data Using Data Diffusion. Cell, 2018. 174(3): p. 716–729 e27.

140. van Heeringen, S.J. and G.J. Veenstra, GimmeMotifs: a de novo motif prediction pipeline for ChIP-sequencing experiments. Bioinformatics, 2011. 27(2): p. 270–1.

