## Supplemental Figures 1-19 for "Resolving thyroid lineage cell trajectories merging into a dual endocrine gland in mammals"

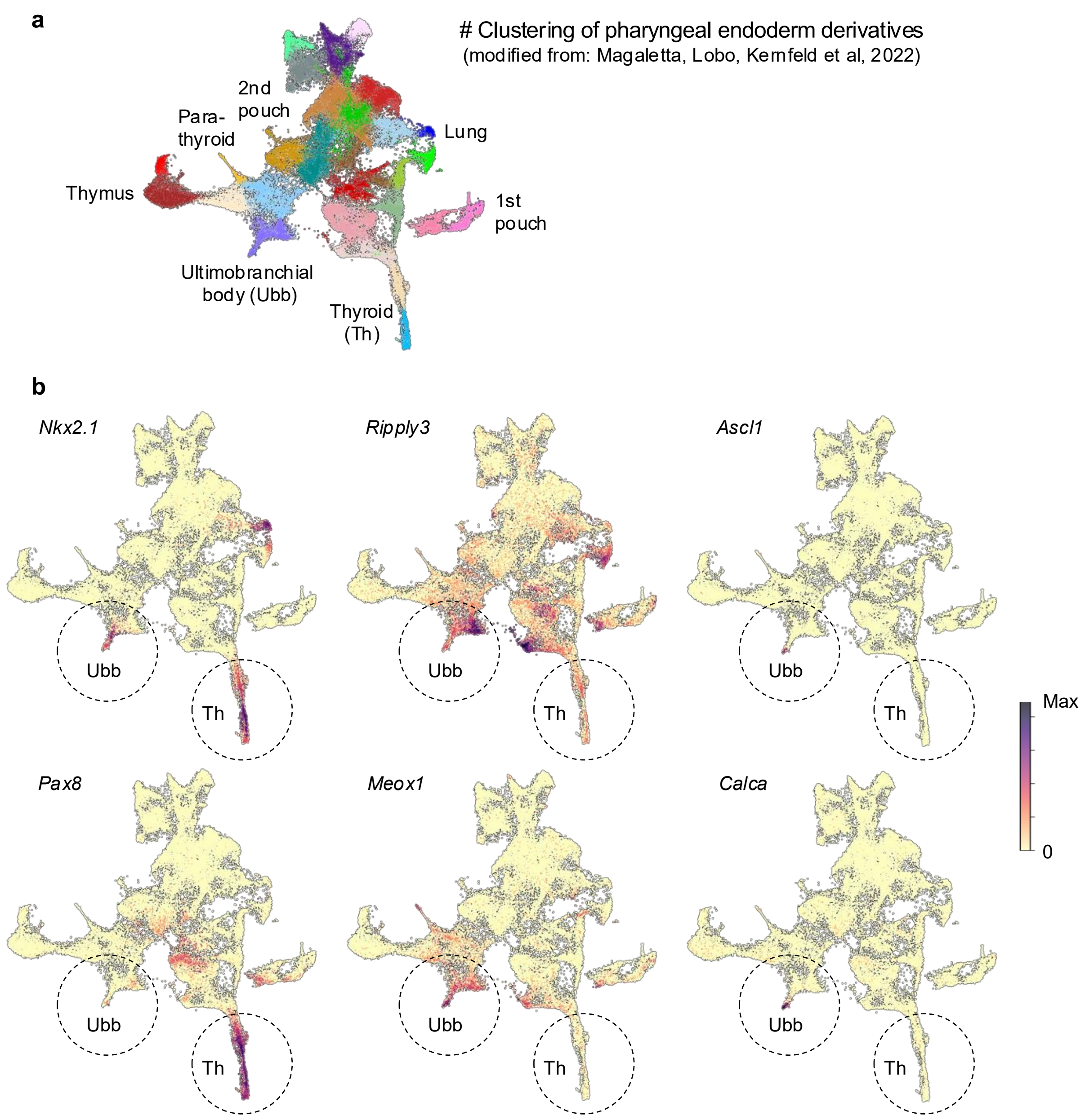


**Supplementary Fig. 1. UMAP visualization of select thyroid and ultimobranchial body marker genes**. Extracted from the single-cell transcriptome atlas previously generated from $Pax9^{VENUS}$ embryos harvested daily between embryonic days (E)9.5 and E12.5 (Magaletta et al [14]). Supplementary to Fig. 1g, h.
**a** Overview depicting unsupervised clustering with the earlier less mature stages being distributed centrally in the UMAP embedding whereas the more mature and organ-specific cells are found in the peripheral clusters, as indicated. **b** Cluster-specific enrichment of the indicated genes of interest. Each dot represents a single cell in the gobal transcriptome space colored by average log2 normalized expression. Thyroid (Th) and ultimobranchial body (Ubb) clusters are encircled. Scalebar indicates log2-normalized expression.

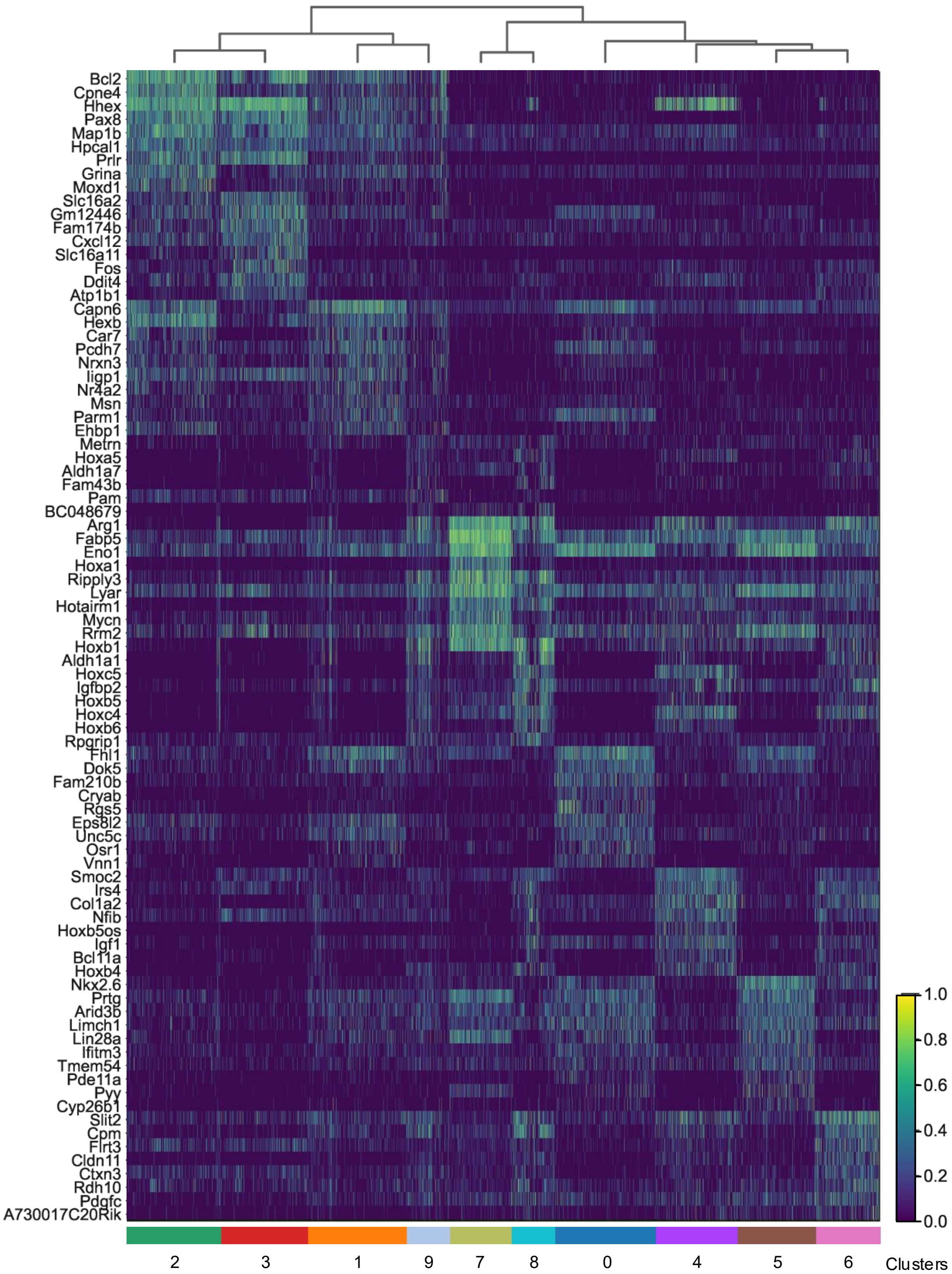


**Supplementary Fig. 2. Clusters with thyroid and ultimobranchial lineage identities distinguished by enriched genes**. Heatmap showing the scaled expression of the of top 10 upregulated genes in each of clusters 0-9 exclusively featuring an adjusted p-value <0.01, a log2-fold change >1.0, and being expressed in >20% of the cells in the cluster. Supplementary to Fig. 1i, k. Genes upregulated in multiple clusters are displayed corresponding to the cluster having maximum average log2-normalized expression of the gene. The dendrogram obtained from unsupervised hierarchical clustering is displayed. Scalebar indicates log2-normalized expression.

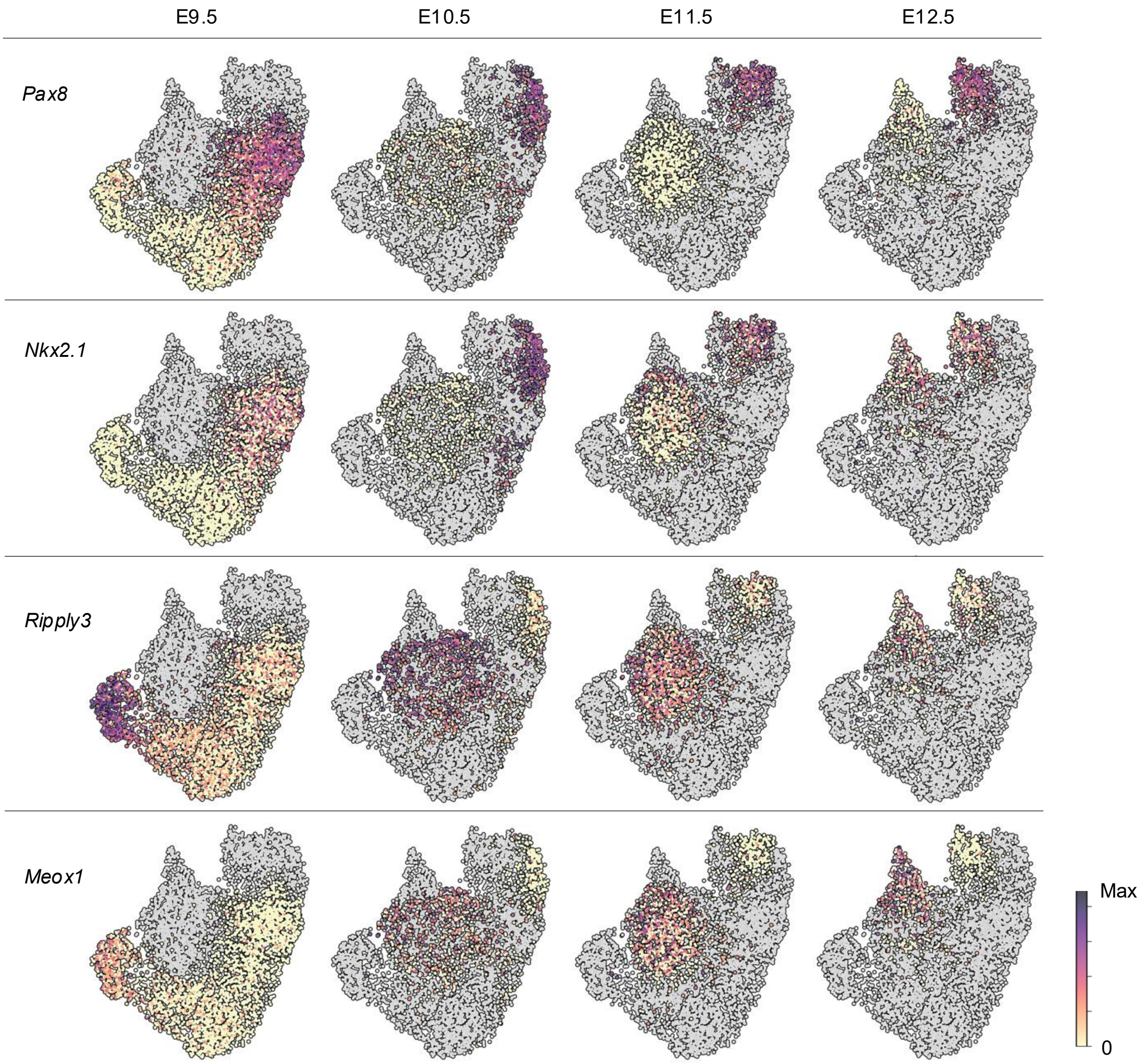


**Supplementary Fig. 3. Single-cell mRNA expression of thyroid and ultimobranchial lineage marker genes across Eday.** UMAP embedding showing expression patterns of *Pax8*, *Nkx2-1*, *Ripply3* and *Meox1* separately at each embryonic day between E9.5 and E12.5 (top panels). Supplementary to Fig. 1i-l. Cells are outlined and colored by the log2-normalized expression if they are present at the corresponding embryonic day. Gray outlined dots represent cells present in the dataset from timepoints other than the one indicated in the plot. Leiden cluster assignments with predicted thyroid lineage clusters 1-3 and Ubb clusters 4, 6 and 8, according to unsupervised global scRNAseq analysis, are shown for comparison (right panel). Scalebar indicates log2-normalized expression. Ubb ultimobranchial body.

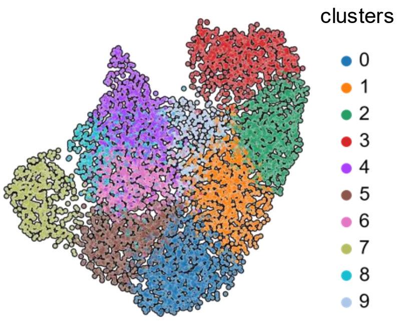

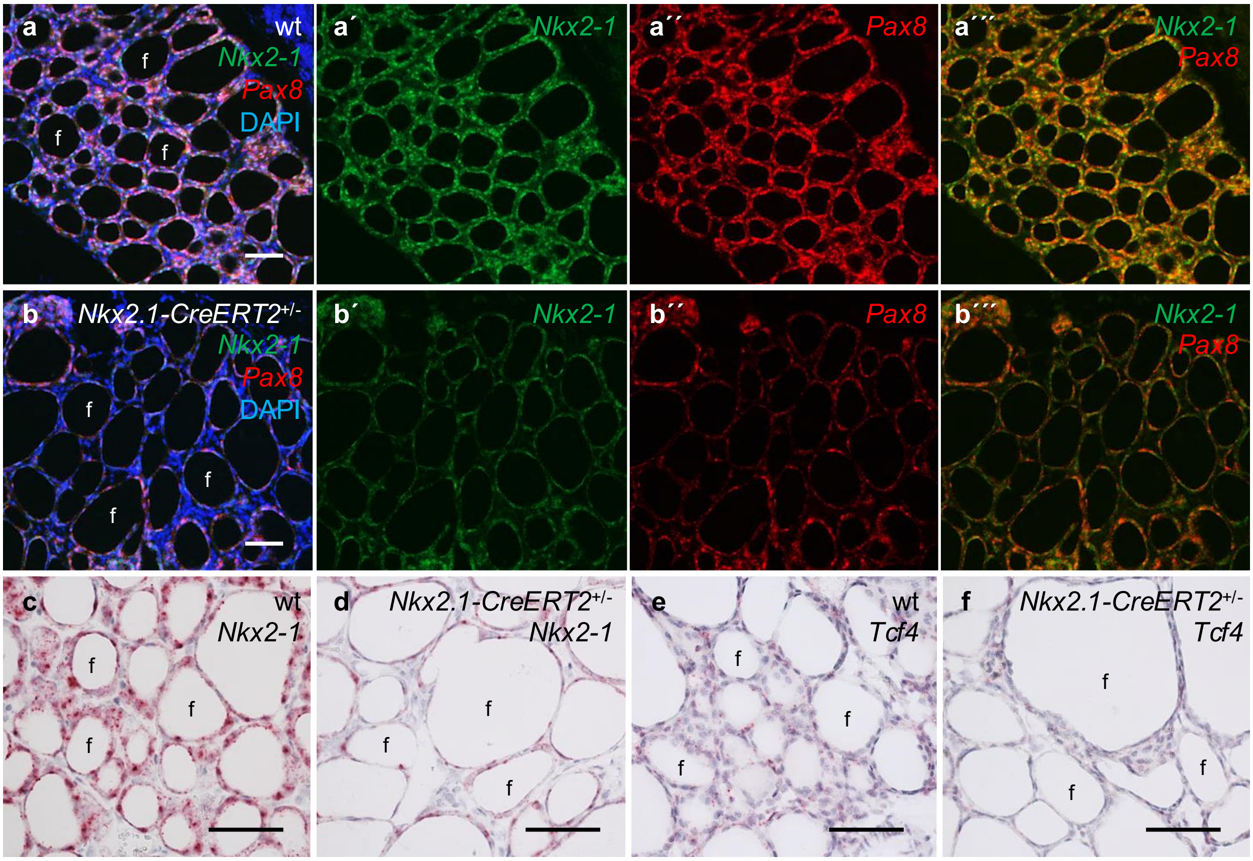


**Supplementary Fig. 4. Downregulation of Pax8 and Tcf4 in heterozygous *Nkx2-1* deficient mouse thyroid cells *in vivo*.** Thyroid FFPE sections of wildtype (wt) and heterozygous *Nkx2.1-CreERT2* mice (age: 6 months) were subjected to RNAscope analysis to validate the *Pax8* gene as a predicted target of Nkx2-1 according to Ordinary Fig. 3e and Supplementary Table 1. This mutant is equivalent to heterozygous Nkx2-1 null mice as evidenced by similarly altered follicular morphology. Multiplex fluorescence was applied for colocalization in (a, b) whereas histochemical detection allowed comparison of high and low abundance mRNA levels in (c-f). **a** Co-expression of *Nkx2-1* and *Pax8* in the majority of thyroid follicular cells. **b** Diminished expression of *Pax8* in cells with reduced *Nkx2-1* expression due induced inactivation of the Cre driver allele. **a´-a´´´, b´-b´´´** show single/double channel without DAPI nuclear staining for improved visualization. Note heterogenous expression of Nkx2-1 and Pax8 is evident among follicular cells in both normal and mutant thyroids. **c-f** Diminished expression of *Tcf4*, another predicted Nkx2-1 target gene (see Supplementary Table 1), in *Nkx2-1* haploinsufficient thyroid cells. f follicles. Scale bars: 100 µm.

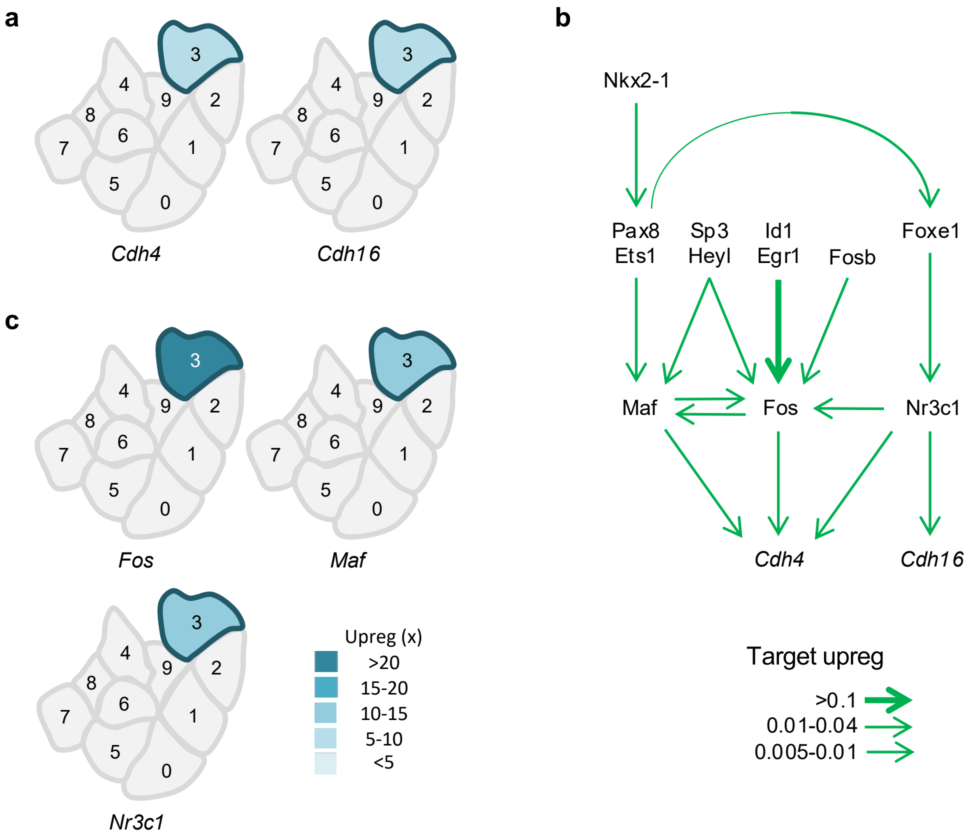


**Supplementary Fig. 5. Enriched expression of cadherin-4 (R-cadherin) and cadherin-16 (Ksp-cadherin) and their predicted transcriptional regulation in embryonic thyroid cells undergoing differentiation. a** Upregulation of *Cdh4* and *Cdh16* in cluster 3 only. Curated from lists comprising all enriched genes per cluster. **b** Predicted subnetwork of *Cdh4* and *Cdh16* regulation in thyroid lineage cells identified by CellOracle. Arrows indicate upregulation with arrow thickness representing mean cluster-specific GRN TF–target gene interaction scores. No TFs were found to downregulate these cadherins in the filtered thyroid and Ubb GRNs. **c** Cluster-specific upregulation of key TFs predicted to regulate *Cdh4* and *Cdh16* expression in cluster 3 cells. Supplementary to Fig. 3a-e. GRF gene regulatory network.

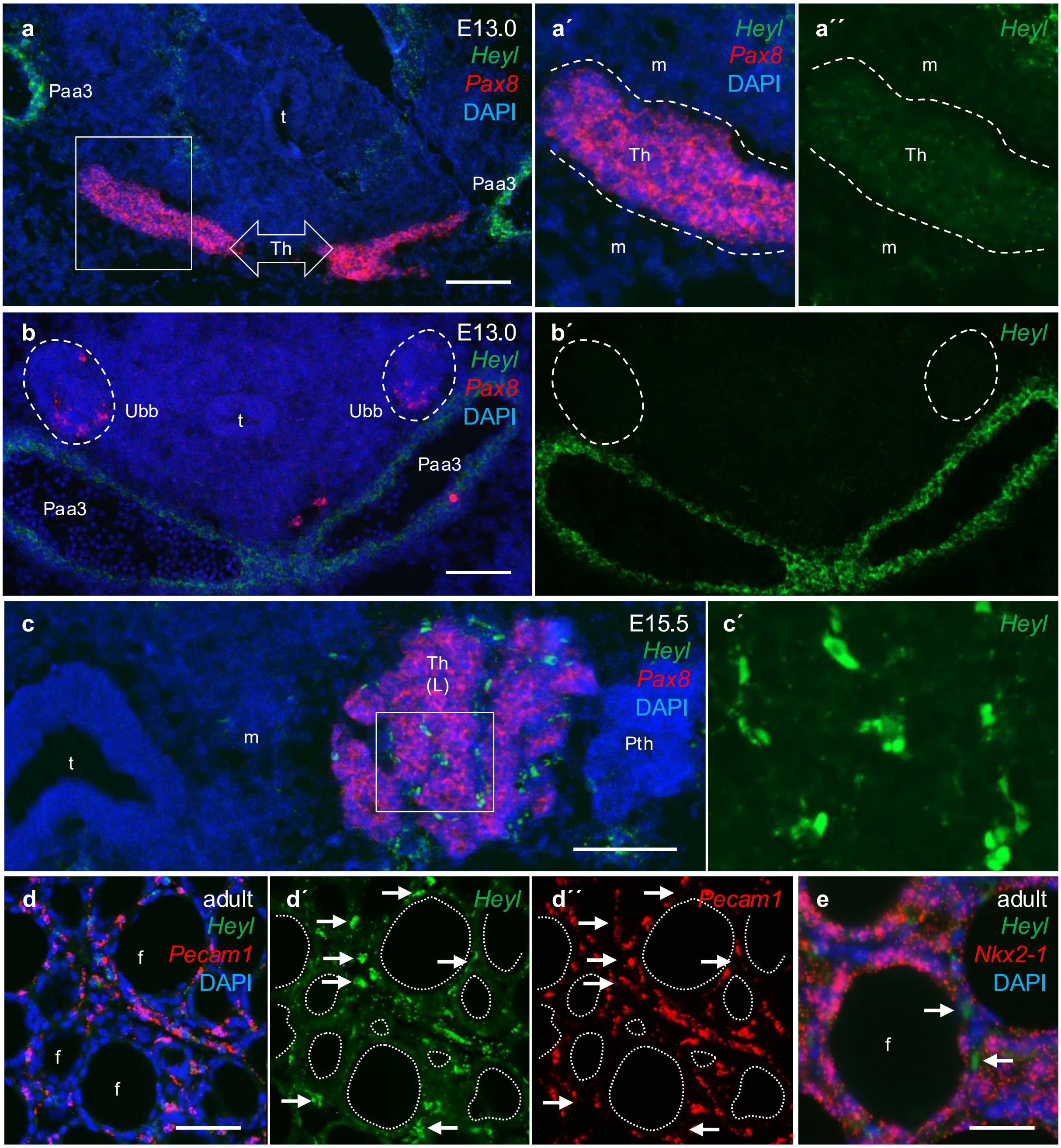


**Supplementary Fig. 6. Heyl expression pattern in the embryonic and adult mouse thyroid gland.** RNAscope Multiplex fluorescent analysis of *Heyl* vs *Pax8* (a-c), *Pecam1* (d) and *Nkx2-1* (e) expression was done to validate scRNAseq data presented in Ordinary Fig. 3. Single channels are shown in (a´´, b´, c´, d´, d´´) for improved clarity. **a** Bilateral growth of the midline thyroid primordium at E13.0; a´, a´´ show high power of boxed area in (a) with weak *Heyl*+ thyroid tissue encircled. **b** Ultimo-branchial bodies prior to fusion with the thyroid; b´ shows the Ubbs are uniformly *Heyl* negative. Note scattered *Pax8*+ Ubb cells confirming immunofluorescent data in Ordinary Fig. 2k. **c** Accumulation of *Heyl*+ interstitial cells in the embryonic thyroid lobe; c´ shows high power of boxed area in (c). **d** Differential *Heyl* expression in the adult thyroid gland: weak in follicular cells; moderate in microvessels; strong in scattered, *Pecam1* negative interstitital cells indicated by arrows in (d´-d´´) in which follicle lumina are outlined for clarity. **e** *Heyl*+ interstitial cells are Nkx2-1 negative (arrows). Th thyroid, Pth parathyroid, Ubb ultimobranchial body, Paa Pharyngeal arch artery, t trachea, m mesenchyme, f follicle, L left. Scale bars: 100 (a-d) and 50 (e) µm.

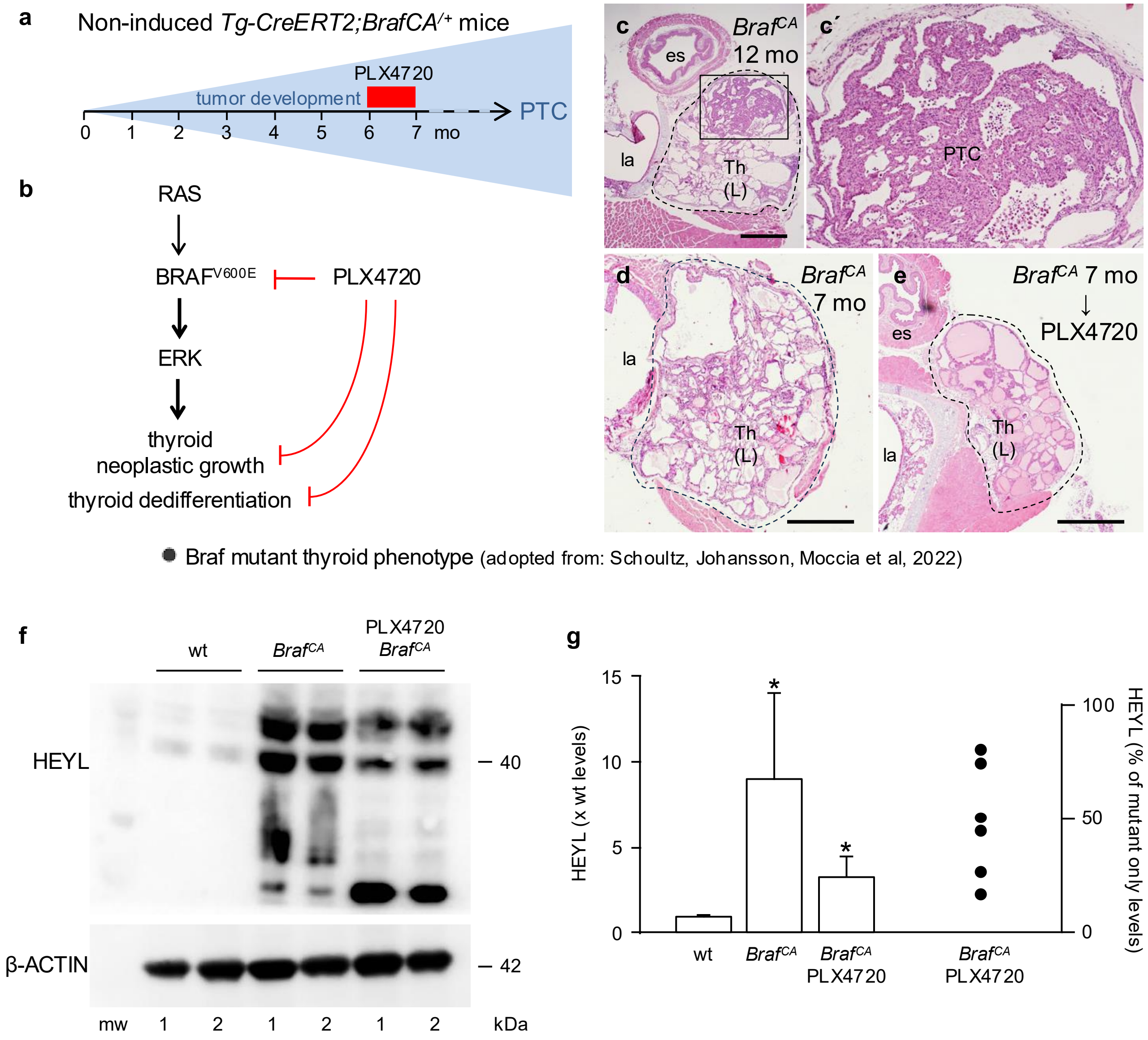


**Supplementary Fig. 7. Upregulation of Heyl expression in Braf mutant thyroid cells.** Data obtained from *Tg-CreERT2;Braf*$^{CA/+}$ mice in which a Braf mutant allele encoding BRAF$^{V600E}$ oncoprotein is conditionally expressed under the thyroglobulin (Tg) promoter. BrafCA activation occurred by spontaneous Cre-mediated recombination (non-induced conditions) conferring multifocal tumor development of papillary thyroid carcinoma (PTC) whereas most of the thyroid tissue remain follicular, as previously described (Schoultz et al [45]). Mutant mice are abbreviated "*Braf*$^{CA}$" throughout. **a** Mouse PTC model and drug treatment protocol. **b** Principal outcome of constitutive activation of the MAPK signaling pathway by BRAF$^{V600E}$ and effects of mutant Braf kinase inhibition in mouse thyroid. **c-e** Thyroid histology in *Braf*$^{CA}$ mice of different age without (c, d) or with (e) PLX4720 treatment (see Methods for details). c′ shows high power of boxed area in (c). Th(L) left thyroid lobe, la larynx, es esophagus, mo months. Scale bars: 500 µm. **f, g** Western blot analyzis of HEYL expression in 7 mo old wildtype (wt) and *Braf*$^{CA}$ mice with/without PLX4720 treatment as outlined in (a). Numbers (1, 2) in (f) indicate duplicate samples; mw molecular weight ladder. Altered HEYL expression in (g) is given relative to the mean wt level set to "1" (bars to the left) or the mean *Braf*$^{CA}$ level set to 100% (dots to the right). Mean±SD (n=6, double samples from 3 mice/group); *p<0.02 (t test) mutants vs wt. Included raw Western blots of HEYL aand the corresponding b-ACTIN blots are shown in Supplementary Dataset 4. These findings validate scRNAseq data provided in Ordinary Fig. 3 altogether indicating Heyl is a new regulated gene in mouse thyroid.

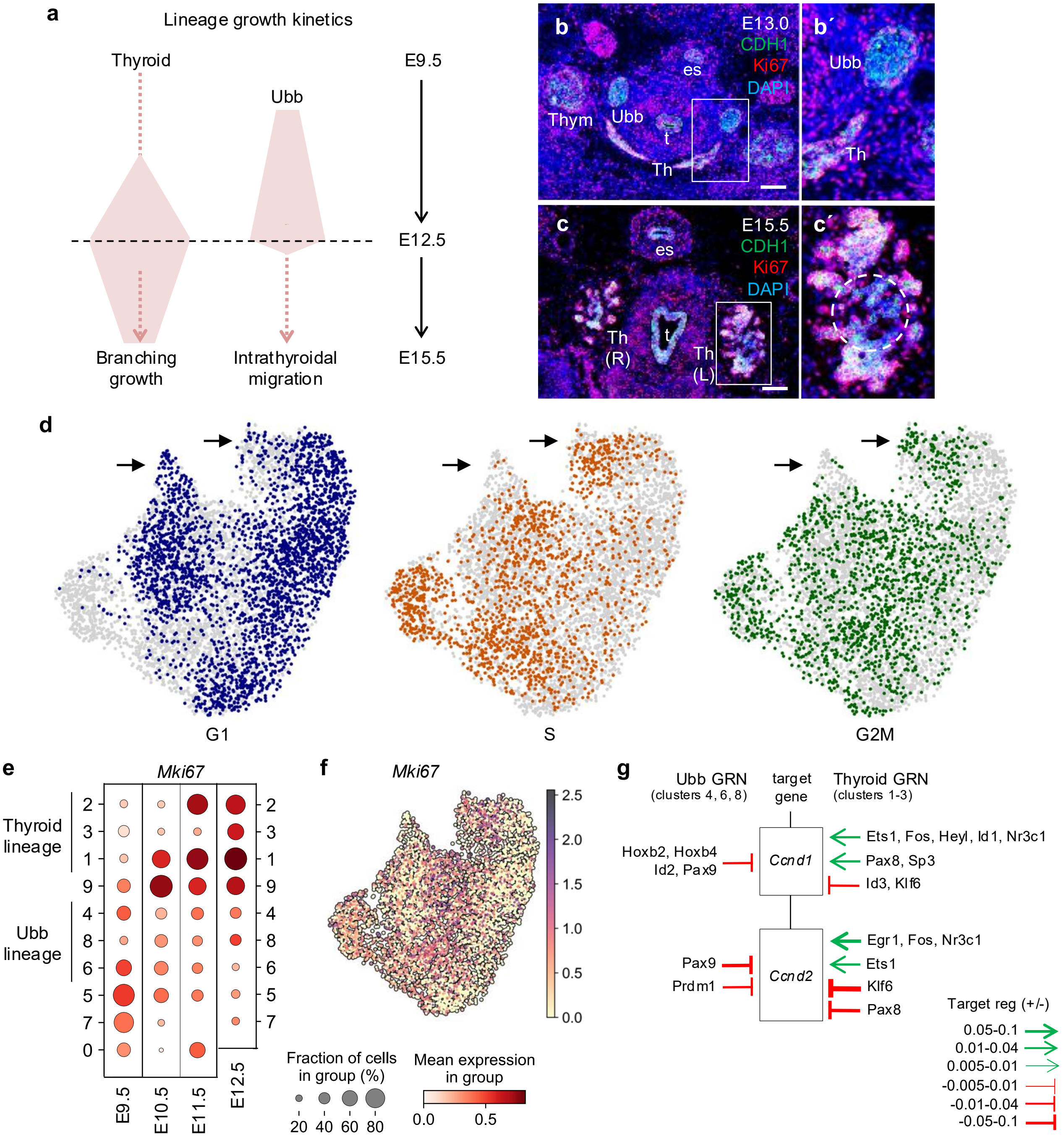


**Supplementary Fig. 8. Different growth dynamics of thyroid and Ubb lineages during development. a** Depicted differences in thyroid and Ubb cell proliferation across embryonic day (Eday) based on Ki67 expression patterns, as previously reported [15, 46]. **b, c** Ki67 expression before (b) and after (c) the paired Ubb merged with the midline thyroid bilaterally. Double immunofluorescence of Ki67 and CDH1/E-cadherin with DAPI nuclear staining. Boxed areas are shown at high power in (b´, c´) with Ubb rudiment encircled in (c´). Th thyroid (R, L right and left lobes), Ubb ultimobranchial body, t trachea, es esophagus, Thym thymus. Scale bars: 100 µm. **d** UMAP embeddings with cells colored according to whether they are predicted to be in cell cycle phase G1 (blue, left), S(orange, center) or G2M (green, right). Cells in each plot are colored gray if they are not in the corresponding phase. **e** Dot plots of cluster-specific *Mki67* expression across Eday. Dot size indicates the fraction of *Mik67*$^{+}$ cells whereas color indicates the mean log2-normalized expression. **f** UMAP embedding of *Mki67*. Scalebar indicates log2-normalized expression. **g** Predicted transcriptional regulation of *Ccnd1* and *Ccnd2* encoding cyclin D in Ubb and thyroid *in silico* GRNs identified by CellOracle. Magnitude of predicted up- and downregulation are indicated by sharp (green) or blunt (red) arrows and arrow thickness representing mean cluster-specific TF-target gene scores.

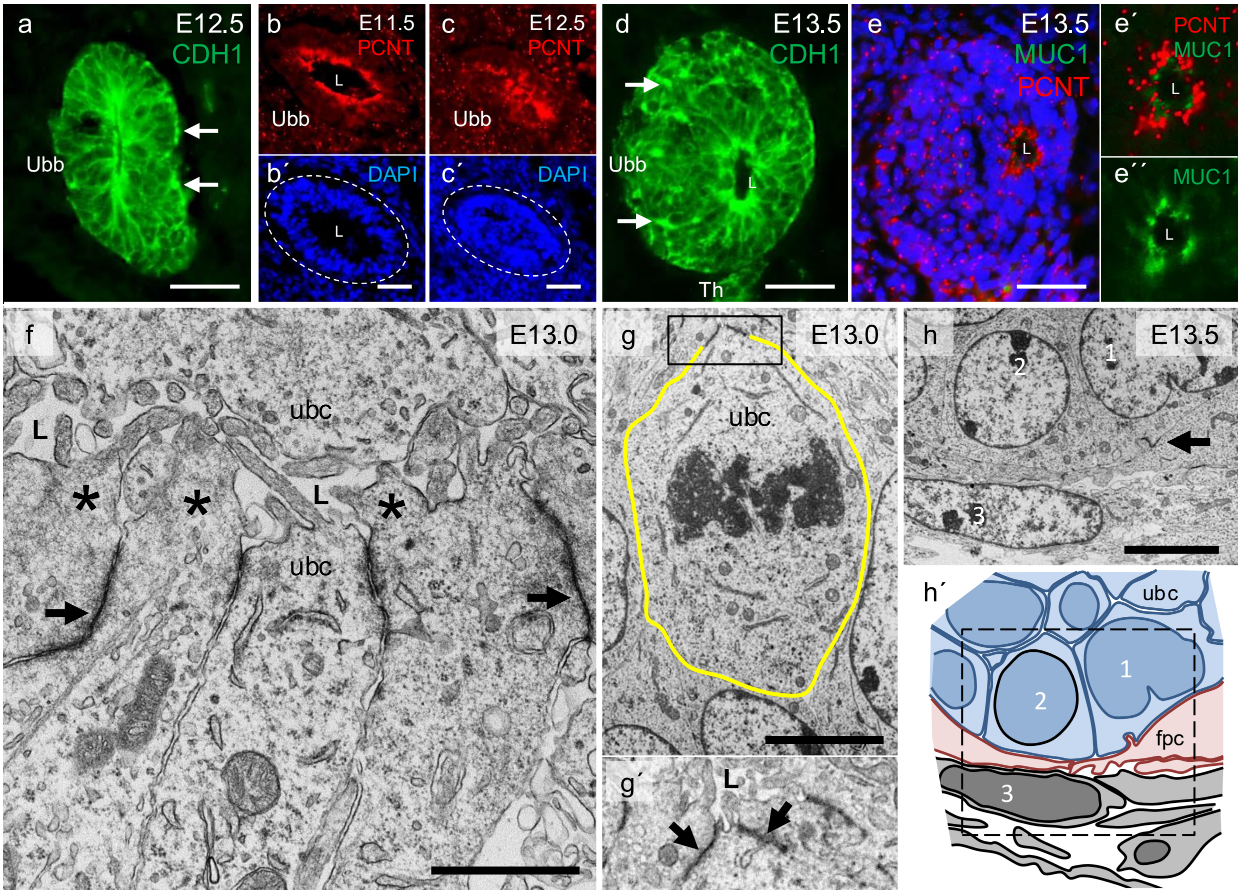


**Supplementary Fig. 9. Morphodynamic changes of the developing Ubb**. Immunofluorescence and transmission electron microscopy of wildtype ultimobranchial bodies (Ubb) between E11.5 and E13.5. **a-e** Lumen involution accompanying epithelial multilayering. Redistribution of CDH1/E-cadherin (a, d), PCNT/pericentrin (b-c´) and MUC1/mucin-1 (e-e´´). Arrows indicate basally located CDH1$^{+}$ foci. **f** Apical constriction of Ubb cells. Arrows indicate elongated adherens junctions; asterisks mark protrusions of apical cytoplasm. **g** Ubb cell (outlined) undergoing oriented cell division signified by mitotic spindle perpendicular to the apical-basal axis. **g´** High power of boxed area in (g) with a much narrowed apical surface of the same mitotic cell. **h** Peripheral portion of Ubb partly invested by cells derived from the midline thyroid primordium. Arrows indicate focal adhesions established between the two cell types. **h´** Cartoon reproducing the motif in (h) (framed) with the deduced cell identities in *blue*: Ubb cells; in *pink*: thyroid cells; in *gray*: mesenchymal cells. Some corresponding cells are numbered 1-3 for clarity. L lumen, ubc ultomobranchial body cell, fpc folliclar progenitor cell. Scale bars: 50 (a-e), 5 (g, h) and 2 (f) µm.

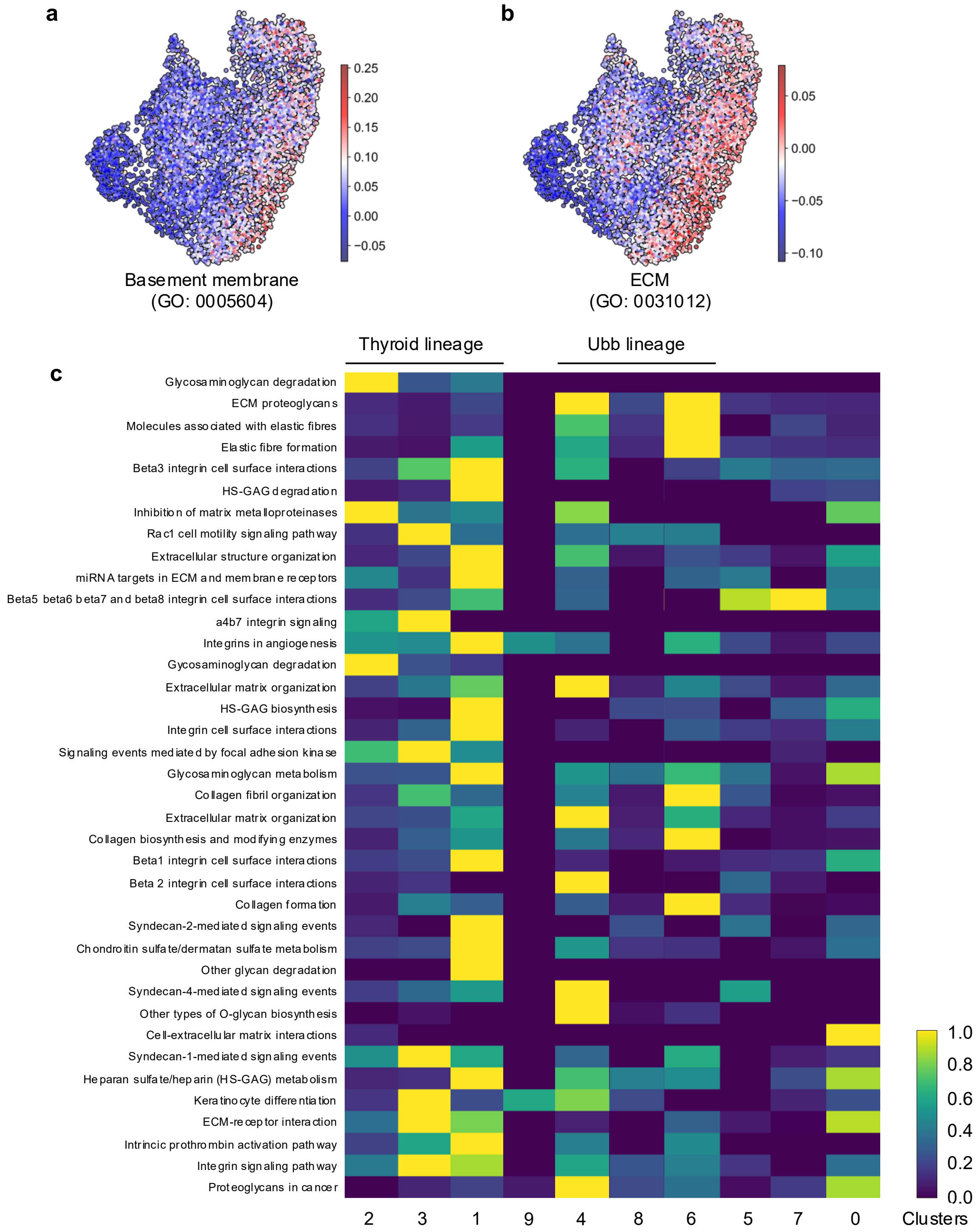


**Supplementary Fig. 10. Differential expression of genes involved in turnover and regulation of extracellular matrix constituents during thyroid and ultimobranchial lineage development. a, b** UMAPs displaying the per cell scores of basement membrane (a) and extracellular matrix (b) genes collectively identified obtained from the Gene Ontology database. Scores are defined as the average log2-normalized expression of the set of genes minus the average log2-normalized expression of a background set. Ubb, ultimobranchial body. Scale bar indicates gene set score. **c** Heatmap of a curated list of pathways from Gene Ontology, Reactome and WikiPathway databases associated with extracellular matrix (ECM) with heat indicating the row normalized −log10 adjusted p-values (Benjamini–Hochberg) obtained from an enrichment analysis (with EnrichR) of the genes differentially upregulated (adjusted p-value < 0.01, log2-fold change > 1.0) in each cluster over a background comprising the remaining clusters. Scale bar indicates scaled –log10 (adjusted value). Supplementary to Fig. 5.

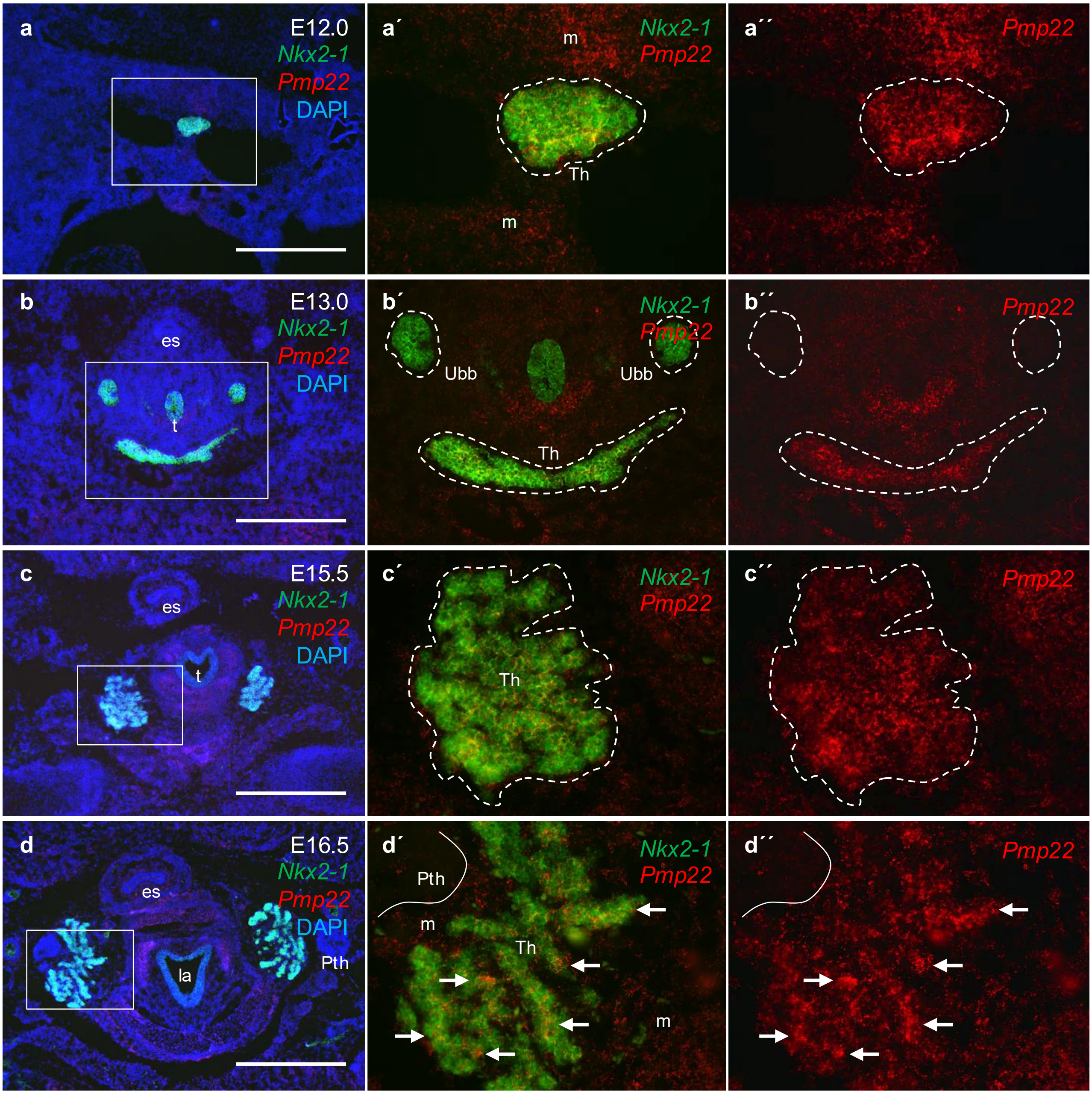


**Supplementary Fig. 11. Pmp22 expression in the developing mouse thyroid.** RNAscope Multiplex fluorescent analysis of *Pmp22* vs *Nkx2-1*. Middle and right panels show high power of the corresponding boxed areas to the left, without DAPI nuclear stain or with red channel only for improved clarity. **a** Midline thyroid primordium at E12.0. **b** Thyroid primordium growing bilaterally at E13.0. **c** Prospective thyroid lobes at E15.5. **d** Thyroid lobes undergoing branching growth at E16.5. Thyroid and ultimobranchial bodies are encircled, parathyroid perimeter outlined. Arrows indicate *Pmp22*+ branching parenchyma. Th thyroid, Pth parathyroid, Ubb ultimobranchial body, t trachea, la larynx, m mesenchyme. Scale bars: 500 µm.

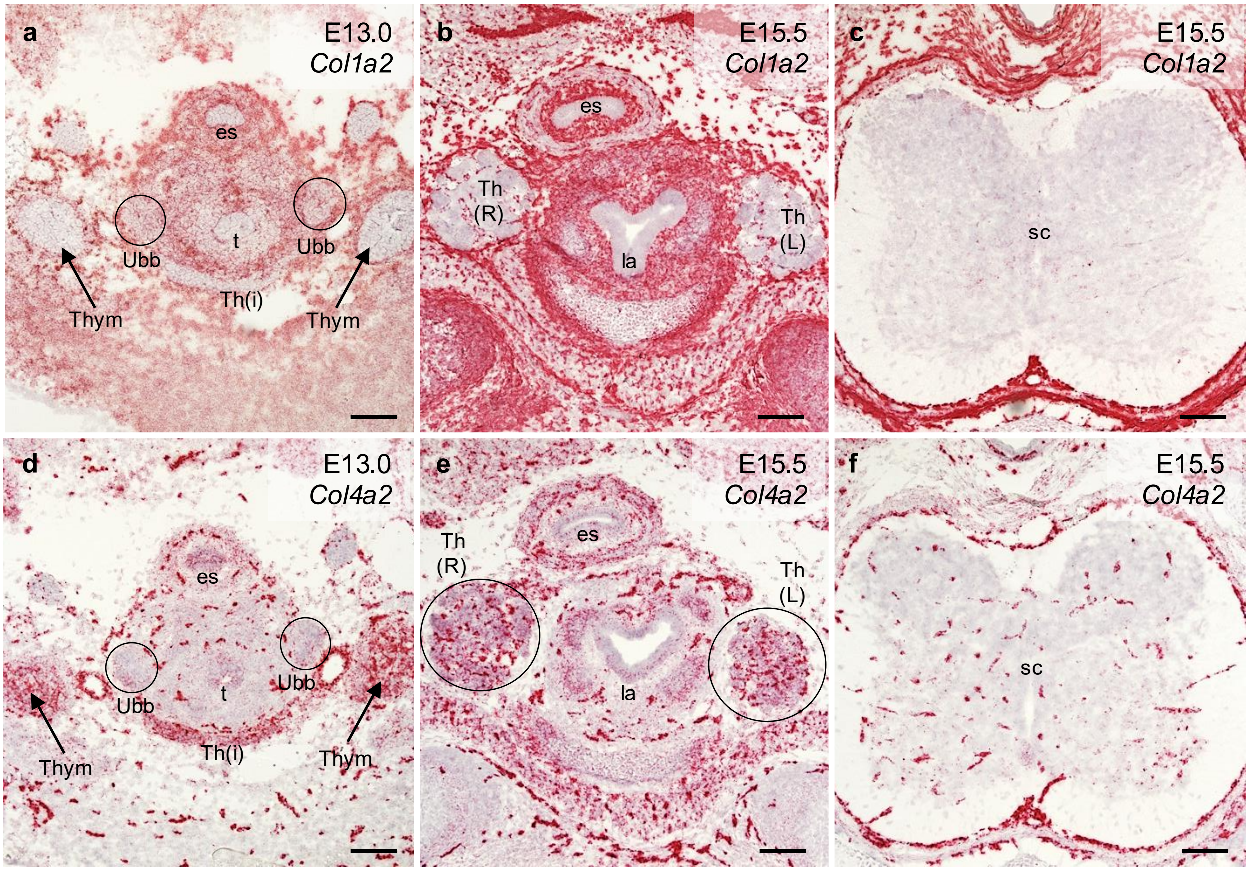


**Supplementary Fig. 12. Differential expression of collagens type I and IV in embryonic neck tissues**. Overview RNAscope images of *Col1a2* (**a-c**) and *Col4a2* (**d-f**) mRNA expression supplementary to Fig. 5c-f. Probes were hybridized onto parallel transverse sections obtained from paraformaldehyde-fixed/frozen E13.0 and E15.5 embryos. Ubb is encircled in (a, d) and right (R) and left (L) thyroid lobes are encircled in (e) för clarity. For comparison, collagen I-negative/collagen IV-positive microvessels are visualized in nervous tissue in (c, f). Th thyroid, Th(i) presumptive isthmus portion of thyroid, Ubb ultimobranchial body, Thym thymus, la larynx, t trachea, es esophagus, sc spinal cord. Scale bars: 100 µm.

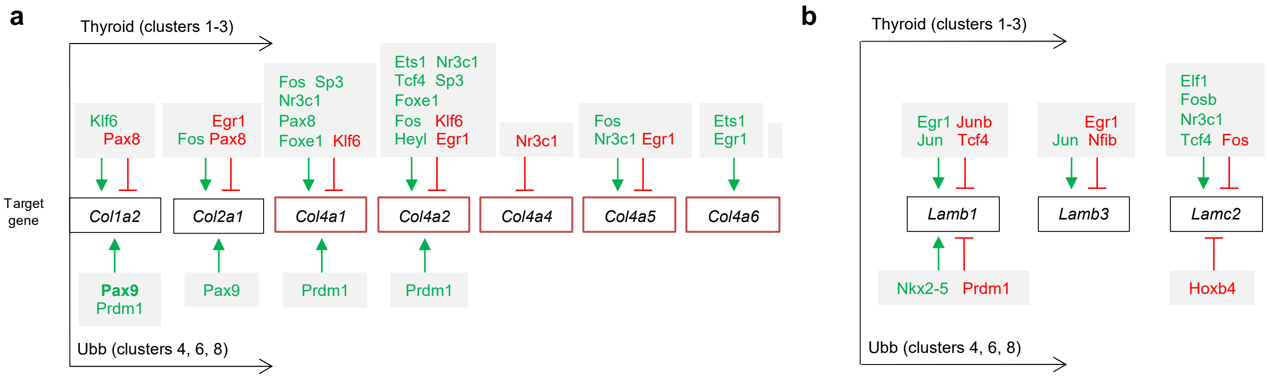


**Supplementary Fig. 13. Inferred regulatory networks of collagen and laminin genes differ between thyroid and Ubb lineage cells.** Schematic subnetworks comprising collagen (*Col*) I and IV genes (**a**) and laminin (*Lam*) genes (**b**) identified by CellOracle in clusters with thyroid and ultimobranchial lineage identities. Genes encoding basement membrane-specific collagen type IV alpha-chains are boxed in red. Predicted up- versus downregulation by the indicated transcription factors (TFs) are colored green or red and by corresponding sharp or blunt arrows. No information is provided on strength of betweenness designating importance of a single TF among others in a given subnetwork. Supplementary to Fig. 5 and 6.

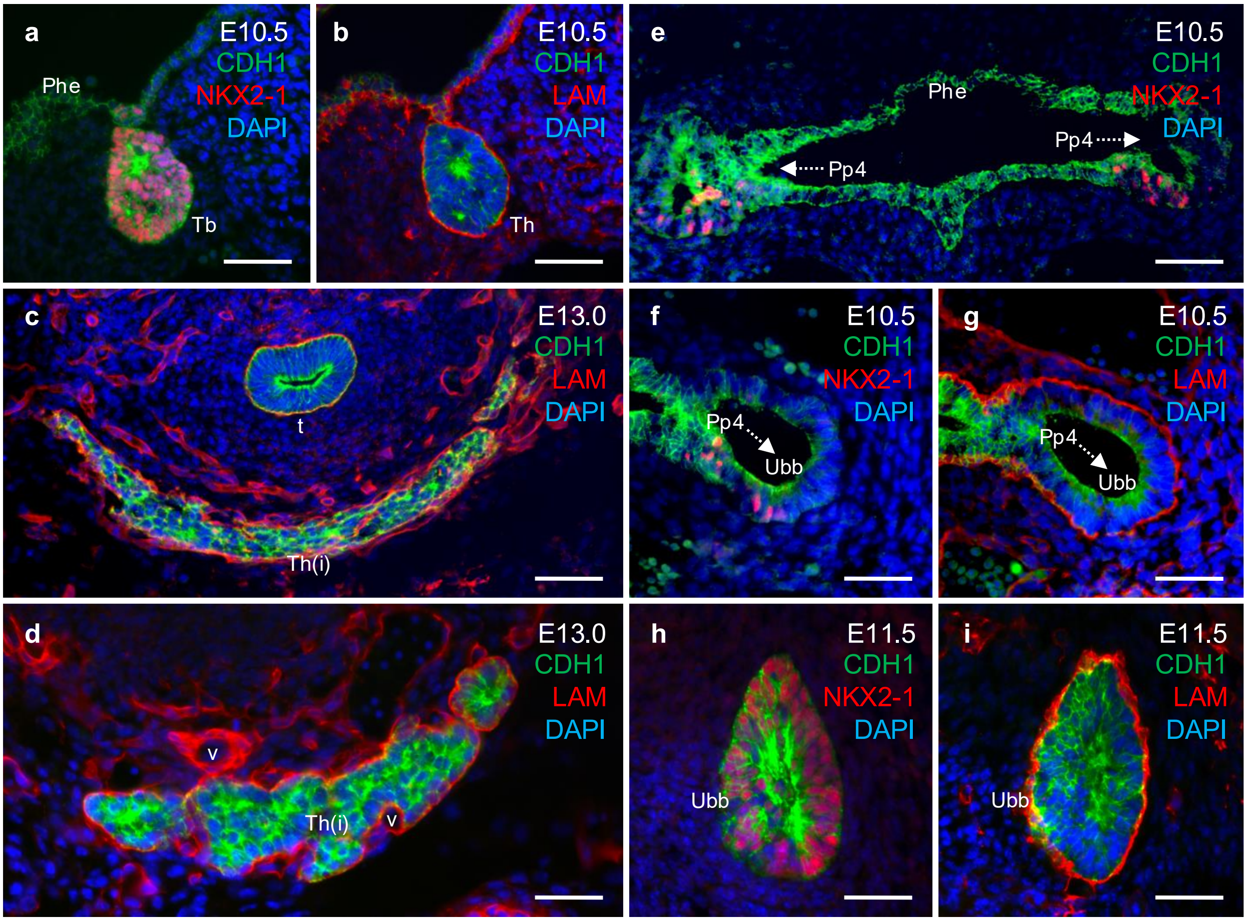


**Supplementary Fig. 14. Laminin in basement membranes of the developing thyroid and ultimobranchial bodies**. Pairwise double immunofluorescence of NKX2-1 versus laminin (LAM) and CDH1/E-cadherin with DAPI nuclear staining for identification of early primordial stages (on parallel sections in: a, b, f, g, h and i). Supplementary to Fig. 6. **a, b** Thyroid bud. **c, d** Presumptive thyroid isthmus. **e-g** Pharyngeal pouch; arrows indicate pouch-Ubb transition. **h, i** Ultimobranchial body. Phe pharyngeal endoderm, Tb thyroid bud, Th(i) isthmus portion of thyroid, Pp4 fourth pharyngeal pouch, Ubb ultimobranchial body, v microvessel. Scale bars: 100 (a-c, e) and 50 (d, f-i) µm.

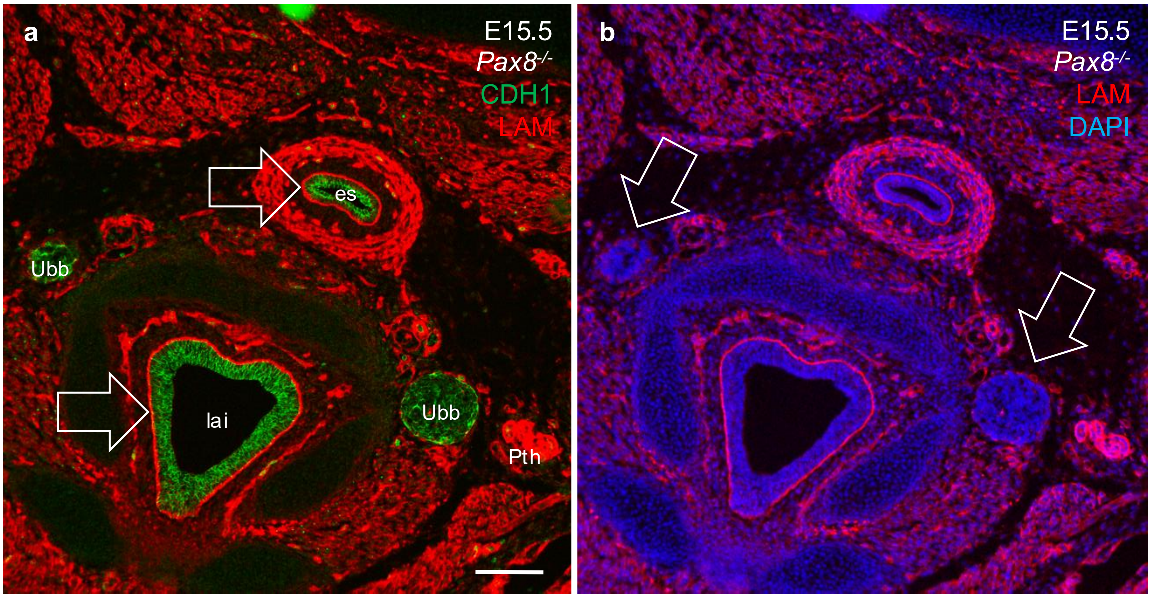


**Supplementary Fig. 15. Selective loss of the Ubb laminin envelope in *Pax8* null mice**. Overview of laminin (LAM) distribution in cross section of anatomical structures in the neck in mutant embryos lacking a thyroid gland. Supplementary to Fig. 6o. **a, b** Identical motifs with co-staining of CDH1/E-cadherin (a) and DAPI (b) for improved visualization of different laminin expression levels among organs. Arrows in (a) indicate LAM$^+$ basement membrane surrounding the esophageal and laryngeal epithelium, respectively. Arrows in (b) indicate discontinuous basement membrane with poor laminin content in Ubb. Pth parathyroid, Ubb ultimobranchial body, es esophagus, lai larynx interior. Scale bar: 100 µm.

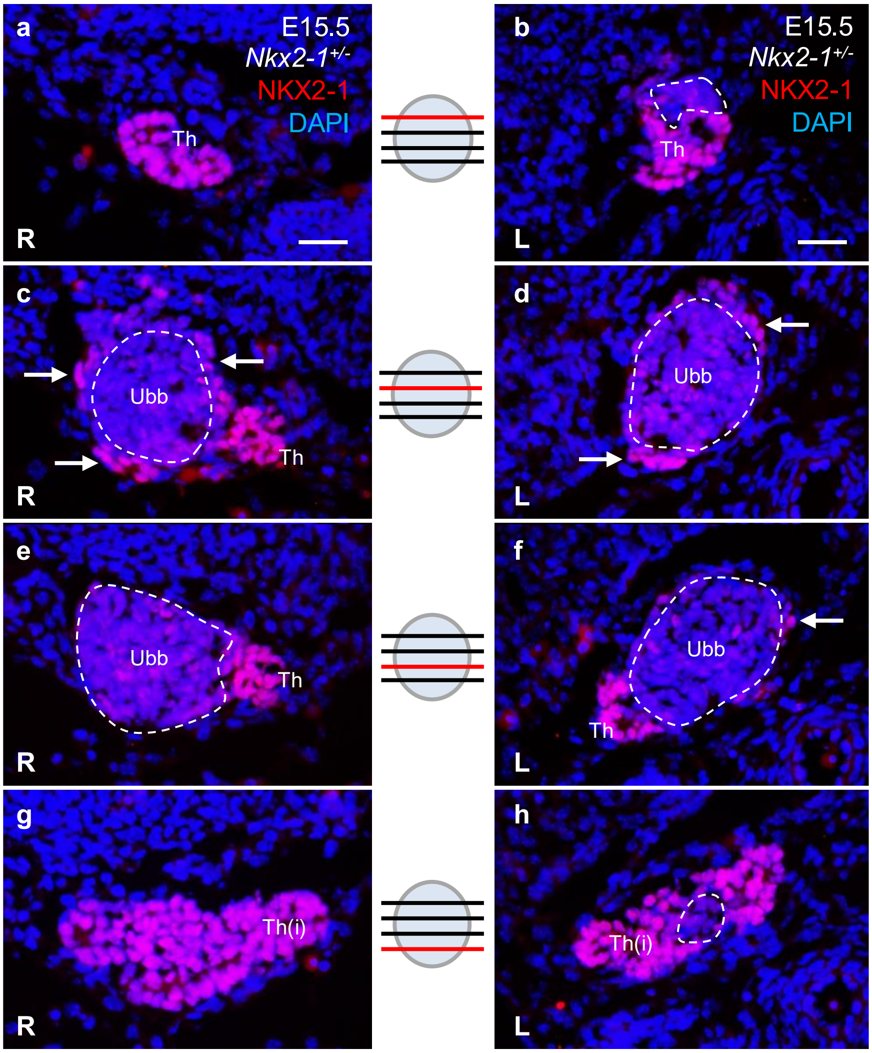


**Supplementary Fig. 16. Impaired thyroid developmental growth in *Nkx2.1* heterozygous knockout mice. a-h** Stacks of NKX2-1 immunofluorescent images from serially sectioned thyroid lobe rudiments obtained at E15.5; corresponding section levels of the two lobes are indicated in centre. Supplementary to Fig. 6p. Residual NKX2-1$^{low}$ cells of the ultimobranchial body (encircled) are incompletely surrounded by NKX2-1$^{high}$ cells (arrows) derived from the thyroid primordium. Lateral ends of the prospective thyroid isthmus are indicated in bottom images. Counterstaining with DAPI identifies all nucleated cells. Th thyroid, (i) isthmus, Ubb ultimobranchial body, R right side, L left side. Scale bar: 25 µm.

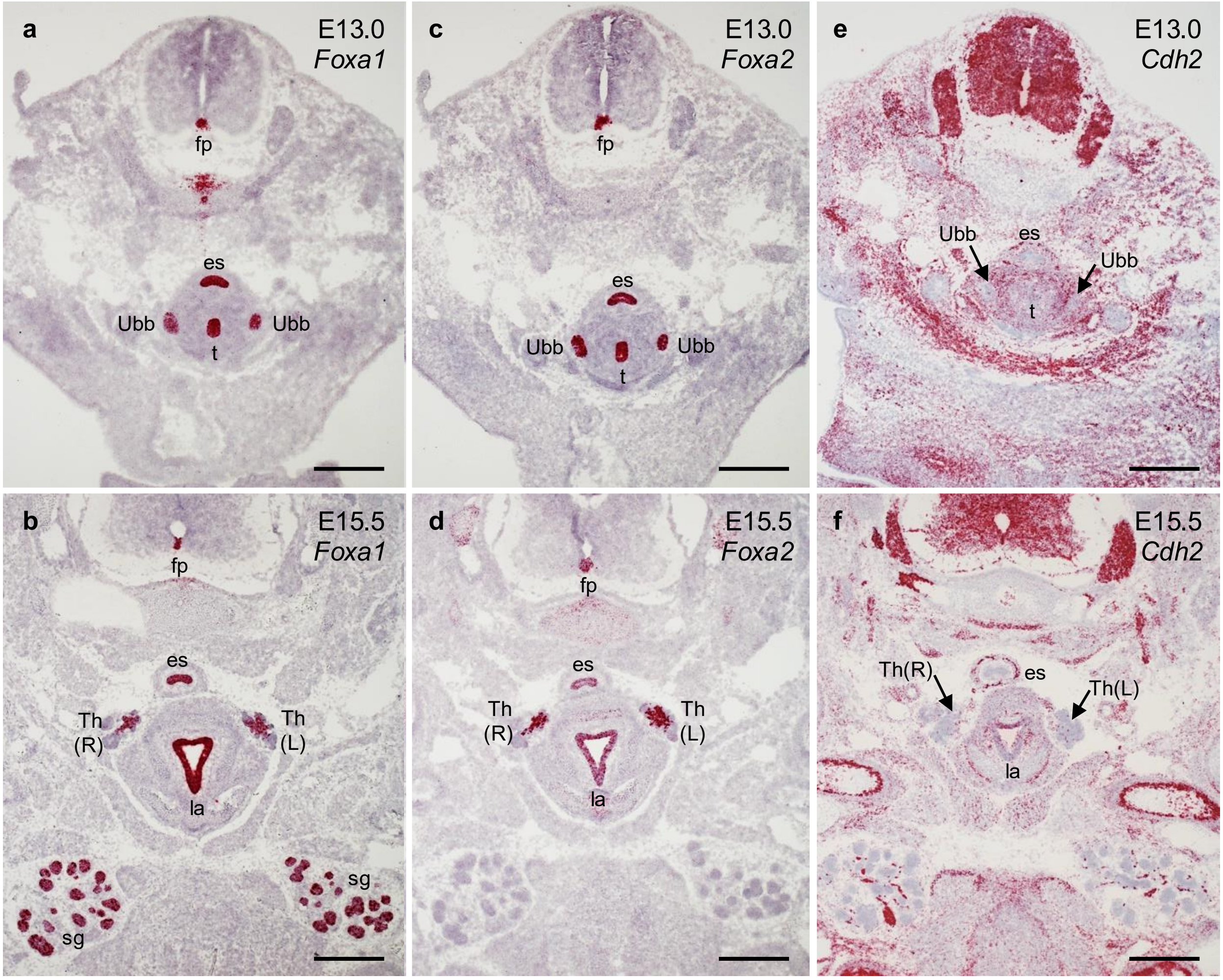


**Supplementary Fig. 17. Expression pattern of Foxa1, Foxa2 and N-cadherin/Cdh2 in embryonic tissues**. Overview RNAscope images of *Foxa1* (**a, b**), *Foxa2* (**c, d**) and *Cdh2* (**e, f**) supplementary to Figs. 7 and 8. Probes were hybridized onto parallel transverse sections obtained from paraformaldehyde-fixed/frozen E13.0 and E15.5 embryos. Fp floor plate, Th thyroid (R, L right and left lobes), Ubb ultimobranchial body, la larynx, es esophagus, sg salivary gland. Scale bars: 500 µm.

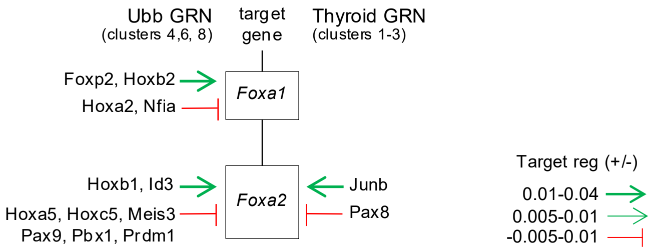


**Supplementary Fig. 18. Transcriptional regulation of *Foxa1* and *Foxa2* in Ubb and thyroid gene regulatory networks as identified by CellOracle.** Predicted up- and downregulation are indicated by sharp (green) or blunt (red) arrows and arrow thickness representing mean cluster-specific GRN TF–target gene interaction scores. Supplementary to Fig. 7g-u.

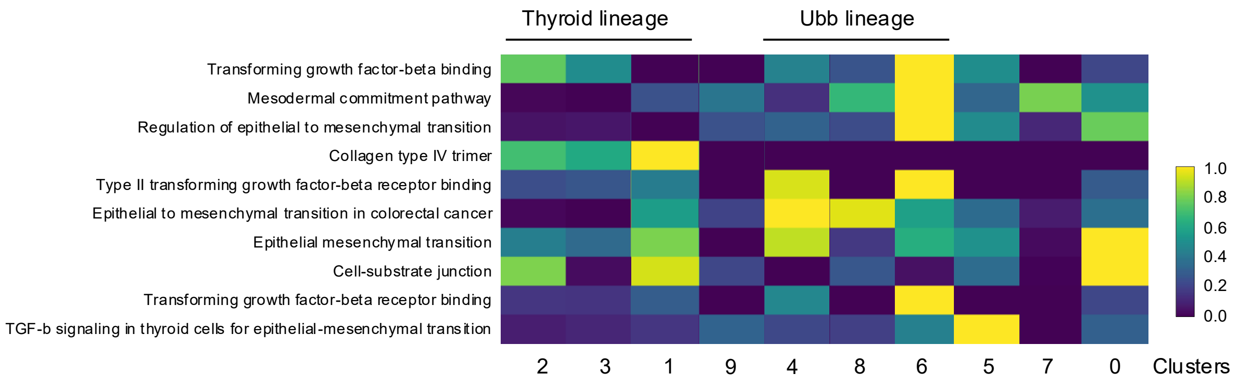


**Supplementary Fig. 19. Differential expression of genes implicated in epithelial-to-mesenchymal transition thyroid and Ubb lineage cells.** Heatmap of a curated list of pathways associated with epithelial-mesenchymal transition (EMT) with heat indicating the row normalized −log10 adjusted p-values (Benjamini–Hochberg). Corresponding cluster numbers and colors are indicated. Ontology terms obtained from Gene Ontology and WikiPathway databases. Supplementary to Fig. 8h-j.
