## Supplemental Table 1 for "Resolving thyroid lineage cell trajectories merging into a dual endocrine gland in mammals"

**Supplementary Table 1.** Predicted Nkx2-1 target genes identified in the gene regulatory network obtained from clusters 1-3 representing the thyroid lineage.

| Nkx2-1 targets | | other predicted regulation by TFs | |
|---|---|---|---|
| gene | | target gene upreg by: | target gene downreg by: |
| upreg (n=33)[1] | | Pax8 (n=5), Foxe1 (n=3), Heyl (n=8) | Pax8 (n=3), Foxe1 (n=2), Heyl (n=6) |
| | coef_mean | | |
| *Rdh10* | 0.029471 | Klf6, Nfe2l2, Nr3c1, Pax8 | Fos |
| *Tcf4* | 0.025189 | Batf3, Fos, Fosb, Heyl, Jun, Nr3c1, Pax8 | Crem, Klf6 |
| *Grip2* | 0.019240 | Egr1, Foxe1, Heyl, Sp3 | Klf6 |
| *Stt3a* | 0.017647 | Egr1, Pax8 | Foxe1, Heyl, Sox9, Sp3 |
| *Fundc2* | 0.015592 | Egr1 | - |
| *Pet100* | 0.014023 | Arntl, Egr1, Elk1, Ets1, Heyl, Id1, Junb | Crem, Fos, Fosb, Id3, Klf6, Nfe2l2, Nr3c1 |
| *Foxp1* | 0.013232 | Fos, Foxe1, Lef1, Nfib, Nr4a2, Sp3 | Heyl, Pax8 |
| *Edc3* | 0.013165 | Klf6 | Creb5, Fosb, Heyl, Junb |
| *Pygm* | 0.012870 | Klf6 | Ets1, Sp3 |
| *Pax8* | 0.012797 | Creb5, Crem, Ets1, Fos, Foxe1, Heyl, Hivep3, Id3, Jun, Maf, Nfic, Rorc, Tcf4 | Fosb, Nfib |
| *Hpcal1* | 0.011732 | Ets1, Klf6, Nr3c1, Pax8, Sp3 | Egr1 |
| *Tmem213* | 0.011022 | Sp3 | Heyl, Id1 |
| *Fance* | 0.010977 | Egr1, Elf3 | Pax8 |
| *S100a10* | 0.010839 | Ets1, Klf6, Nr3c1 | Sp3 |
| *Hoxb2* | 0.010095 | Bcl11b Ets1, Id1, Nr4a2, Sp3 | Egr1, Fos, Klf6, Nr3c1, Tcf4 |
| *Luzp2* | 0.008979 | - | - |
| *Stxbp6* | 0.008855 | Egr1, Ets1, Heyl | Elf3, Klf6, Nfib |
| *Scaper* | 0.007441 | - | - |
| *Aars2* | 0.007268 | - | - |
| *Atp5d* | 0.006988 | Cux1, Egr1, Heyl, Id1, Id3, Nfe2l2, Pax8 | Bach1, Foxb, Foxe1 |
| *Mypop* | 0.006813 | - | - |
| *Kcmf1* | 0.006807 | Egr1, Heyl, Id1 | Id3 |
| *Lmo4* | 0.006045 | Bcl11b, Egr1, Id1, Isl1, Klf6, Maf, Nr3c1 | Heyl, Pax8 |
| *Kctd12b* | 0.005887 | - | - |
| *Mapk1ip1* | 0.005732 | Egr1, Klf6 | Elf3, Fos, Sp3 |
| *Ubash3b* | 0.005693 | Cux1, Egr1, Ets1, Jun, Nr3c1, Sp3 | Foxe1, Klf6 |
| *Flrt1* | 0.005598 | - | - |
| *Greb1l* | 0.005409 | Ets1, Fos, Sp3, Tcf4 | Klf6 |
| *Snx14* | 0.005401 | Heyl, Jun, Nr3c1 | Fos |
| *Faap100* | 0.005353 | - | - |
| *Cyp39a1* | 0.005300 | Klf6 | Egr1, Fos |
| *Sestd1* | 0.005062 | Egr1 | |
| *Invs* | 0.005006 | Id3 | Heyl |
| downreg (n=18)[2] | | Pax8 (n=1), Foxe1 (n=0), Heyl (n=1) | Pax8 (n=1), Foxe1 (n=0), Heyl (n=3) |
| *Cadm1* | -0.028725 | Egr1, Sp3 | Fos, Nr3c1 |
| *Hdhd2* | -0.017271 | Egr1, Elf3, Heyl, Sp3 | - |
| *Igfbp2* | -0.016711 | Fos, Isl1, Maf, Sp3 | Bach1, Egr1, Heyl, Id1, Id3 |
| *Pole3* | -0.013811 | - | Nr3c1 |
| *Uqcrh* | -0.012875 | Elf3 | - |
| *Hspd1* | -0.012783 | Egr1 | Bcl11b, Creb5, Fos, Heyl, Id3, Junb, Klf6, Nr3c1, Nr4a2, Sp3 |
| *Ccar2* | -0.012654 | Elk3, Ets1 | Klf6 |
| *Sdhd* | -0.011841 | Egr1 | Klf6, Nr3c1, Nr4a2 |
| *Snrpd2* | -0.011760 | Egr1, Elf3, Elk3 | Fos, Klf6 |
| *Pno1* | -0.009733 | Egr1 | Arntl, Id3 |
| *Gnas* | -0.008980 | Arntl, Egr1, Elf3, Ets1, Heyl, Id3 | Fos, Hivep3, Klf6, Maf, Sp3 |
| *Pfn2* | -0.008870 | Junb | Egr1, Heyl, Id3 |
| *Cldn9* | -0.007990 | - | Pax8 |
| *Orc6* | -0.007936 | Egr1, Ets1 | Klf6, Sp3 |
| *Sept11* | -0.007780 | Hoxb2 | Id3 |
| *Adprhl2* | -0.007260 | Ets1, Nfe2l1 | - |
| *Angpt2* | -0.006513 | Batf3, Jun, Isl1, Mecom, Pax8 | Klf6, Nr3c1, Sp3 |
| *Exo5* | -0.005435 | Nr3c1 | Egr1, Mecom |

TF transcription factor
[1]listed from highest to lowest scores (coef_mean)
[2]listed from lowest to highest scores (negative coef_mean)
