## Supplemental Table 2 for "Resolving thyroid lineage cell trajectories merging into a dual endocrine gland in mammals"

**Supplementary Table 1.** Predicted Nkx2-1 target genes identified in the gene regulatory network obtained from clusters 4, 6 and 8 representing the Ubb/C-cell lineage.

| Nkx2-1 targets | | other predicted regulation by TFs | |
|---|---|---|---|
| gene | | target gene upregulated by: | target gene downregulated by: |
| upreg (n=35)[1] | | Pax9 (n=4), Prdm1 (n=4) | Pax9 (n=3), Prdm1 (n=7) |
| | coef_mean | | |
| *Pfn2* | 0.049089 | Id2, Id3 | Nkx2-5 |
| *Igfbp2* | 0.032229 | Id2 | Id3, Prdm1 |
| *Hpcal1* | 0.031613 | - | Meis2, Prdm1 |
| *Nrp2* | 0.021344 | Pax9 | Id2, Id3, Prdm1 |
| *Cspp1* | 0.020172 | Irx2 | - |
| *Pno1* | 0.019892 | Id2 | - |
| *Foxp1* | 0.018758 | Foxp2, Id3, Nfia, Nfib, Pax9 | Hoxb2, Hoxb4, Hoxc5, Id2, Irx2, Prdm1 |
| *Stxbp6* | 0.016903 | - | - |
| *Fxyd7* | 0.016773 | Pax9 | - |
| *Msrb2* | 0.015009 | - | - |
| *Flrt1* | 0.014910 | Id2, Pbx1, Prdm1 | Id3, Nfib, Nkx2-5 |
| *Faap100* | 0.013429 | - | Nfia, Pax9 |
| *Cops5* | 0.013040 | - | - |
| *Upk1a* | 0.012801 | Prdm1 | - |
| *St8sia6* | 0.011184 | - | - |
| *Magi1* | 0.010506 | - | - |
| *Pet100* | 0.009633 | Nkx2-5 | Id3, Prdm1 |
| *Rspo1* | 0.009595 | - | - |
| *Hoxb4* | 0.009513 | Hoxa5, Hoxb1, Hoxb2, Hoxb5, Hoxc5, Meis1, Meis2, Prdm1, Zfhx3 | Id2, Id3 |
| *Dbp* | 0.009280 | - | - |
| *Gstk1* | 0.008742 | Prdm1 | - |
| *Nol10* | 0.007874 | - | - |
| *Med26* | 0.007475 | - | - |
| *Copz2* | 0.007414 | Nkx2-5 | - |
| *Tmem213* | 0.006635 | Zfhx3 | Id2 |
| *Aars2* | 0.006408 | - | - |
| *Tnfrsf12a* | 0.005858 | Pax9 | - |
| *Dctd* | 0.005678 | - | Pax9 |
| *Kat5* | 0.005637 | - | - |
| *Timm23* | 0.005542 | Id2 | Pax9 |
| *Poc5* | 0.005315 | - | - |
| *Nefm* | 0.005279 | Id2, Meis1, Nfib, Pbx1 | Hoxb2, Hoxc5, Foxp2, Meis2, Nfia, Prdm1 |
| *Ptcd3* | 0.005201 | Hoxa2, Hoxb6 | Bach2 |
| *Tmem171* | 0.005153 | Hoxb2 | - |
| *Med14* | 0.005080 | - | Prdm1 |
| downreg (n=53)[2] | | Pax9 (n=7), Prdm1 (n=14) | Pax9 (n=0), Prdm1 (n=5) |
| *Fgf8* | -0.096228 | Irx2, Prdm1 | - |
| *Cadm1* | -0.044912 | Prdm1 | Nkx2-5 |
| *Hspd1* | -0.044258 | Id3 | Nkx2-5, Prdm1 |
| *Uqcrh* | -0.040713 | - | Etv1, Prdm1 |
| *Orc6* | -0.038694 | Nkx2-5 | - |
| *Galk1* | -0.035238 | Id3, Nkx2-5 | Bach2 |
| *Lmo4* | -0.033411 | Id2, Meis1, Prdm1 | Hoxa2, Hoxb4, Hoxc5, Id3, Nkx2-5 |
| *Efna5* | -0.026840 | Hoxa2, Hoxb4 | - |
| *Lpar4* | -0.024157 | Prdm1 | - |
| *Rad54l* | -0.022496 | - | Nkx2-5 |
| *Trip13* | -0.021666 | - | Etv1 |
| *Sdhd* | -0.020869 | Pax9 | Nkx2-5 |
| *Tada1* | -0.018832 | Pax9 | - |
| *Anxa2* | -0.016896 | Nfib, Nkx2-5 | Prdm1 |
| *Foxg1* | -0.016151 | Id2 | Id3, Nkx2-5, Prdm1 |
| *Pole3* | -0.015971 | - | - |
| *Gtf3c6* | -0.015043 | Id3, Prdm1 | - |
| *Cited2* | -0.014739 | Id2, Id3, Prdm1 | - |
| *Hoxb6* | -0.012521 | Hoxb2, Hoxb4, Hoxc5, Prdm1 | Id3, Meis2, Nkx2-5 |
| *Casd1* | -0.011648 | - | - |
| *Atp5d* | -0.011592 | Meis1, Pax9 | Id2, Prdm1 |
| *Edc3* | -0.011537 | - | - |
| *Abcb8* | -0.011396 | Id3, Prdm1 | - |
| *Haus5* | -0.011052 | - | - |

| gene | coef_mean | target gene upregulated by: | target gene downregulated by: |
|---|---|---|---|
| *Tgif1* | -0.010553 | Bach2, Hoxb2, Id2, Meis1, Meis2, Nkx2-5, Sox4 | Meis3, Nfib |
| *Nexn* | -0.010532 | - | - |
| *Hdhd2* | -0.010479 | Bach2, Id3 | - |
| *Capn10* | -0.010371 | Prdm1 | - |
| *Rps6ka3* | -0.009940 | - | Nkx2-5 |
| *Gas2* | -0.009886 | Hoxb6, Id2, Meis1, Meis2, Meis3, Pbx1 | Hoxb2, Nkx2-5 |
| *Tmem65* | -0.009833 | Prdm1 | - |
| *Arl4a* | -0.008867 | - | - |
| *Dera* | -0.008383 | - | Nfia, Nkx2-5 |
| *Prep* | -0.008370 | - | - |
| *Smndc1* | -0.008251 | Pax9 | Nkx2-5 |
| *Hoxb2* | -0.007594 | Hoxa5, Hoxb1, Hoxb4, Hoxb5, Hoxc5, Id3, Meis2, Nfib, Prdm1, Zfhx3 | Nfia |
| *Slc6a15* | -0.007575 | Meis1 | - |
| *Cog8* | -0.007176 | Nkx2-5, Prdm1 | - |
| *Dhx32* | -0.007130 | Bach2, Pax9, Prdm1 | Meis2 |
| *Pgam2* | -0.006947 | - | - |
| *Zmiz2* | -0.006737 | - | Nkx2-5 |
| *Rmi2* | -0.006449 | - | - |
| *Sgk3* | -0.006283 | Id3 | - |
| *Slitrk4* | -0.006168 | - | - |
| *Cnot3* | -0.006048 | Nfia, Pax9 | - |
| *Kcnj2* | -0.006029 | - | Id3 |
| *Mapk1ip1* | -0.005805 | Pax9 | - |
| *Pcca* | -0.005648 | - | - |
| *Ubash3b* | -0.005576 | - | - |
| *Pitx1* | -0.005548 | Id3 | Hoxb2, Id2, Pbx1 |
| *Polr1a* | -0.005503 | - | - |
| *Xab2* | -0.005454 | Prdm1 | Etv1 |
| *Snx14* | -0.005389 | - | - |

TF transcription factor
[1]listed from highest to lowest scores (coef_mean)
[2]listed from lowest to highest scores (negative coef_mean)
