## Supplemental Table 3 for "Resolving thyroid lineage cell trajectories merging into a dual endocrine gland in mammals"

**Supplementary Table 3.** Nkx2-1 target genes shared by thyroid and Ubb/C-cell lineage cells.

| Gene | Thyroid lineage (clusters 1-3) | Ubb lineage (clusters 4, 6, 8) | Main function(s) |
|---|---|---|---|
| *Aars2* | ↑ | ↑ | mitochondrial function (Alanyl-tRNA synthetase 2) |
| *Atp5d* | ↑ | ↓↓ | mitochondrial function (ATP synthase delta subunit) |
| *Cadm1* | ↓↓↓ | ↓↓↓↓↓ | cell adhesion (Cell adhesion molecule 1) |
| *Edc3* | ↑↑ | ↓↓ | mRNA turnover (Enhancer of mRNA decapping protein 3) |
| *Faap100* | ↑ | ↑↑ | DNA repair (Fanconi anemia core complex protein) |
| *Flrt1* | ↑ | ↑↑ | cell adhesion (Fibronectin leucine rich transmembrane protein 1) |
| *Foxp1* | ↑↑ | ↑↑ | development (Forkhead box P1 transcription factor) |
| *Hdhd2* | ↓↓ | ↓↓ | metabolism (Haloacid dehalogenase-like hydrolase domain containing protein 2) |
| *Hoxb2* | ↑↑ | ↓ | development (Homeobox B2 transcription factor) |
| *Hpcal1* | ↑↑ | ↑↑↑↑ | calcium binding, ferroptosis, autophagy (Hippocalcin-like protein 1) |
| *Hspd1* | ↓↓ | ↓↓↓↓↓ | mitochondrial function (Heat shock protein 60) |
| *Igfbp2* | ↓↓ | ↑↑↑↑ | growth regulation (Insulin-like growth factor binding protein 2) |
| *Lmo4* | ↑ | ↓↓↓↓ | development, transcriptional coregulator (LIM domain only 4) |
| *Mapk1ip1* | ↑ | ↓ | growth regulation (MAPK scaffold protein 1) |
| *Orc6* | ↓ | ↓↓↓↓ | DNA replication (Origin recognition complex subunit 1) |
| *Pet100* | ↑↑ | ↑ | Mitochondrial function (Cytochrome C oxidase chaperone) |
| *Pfn2* | ↓ | ↑↑↑↑↑ | actin-based cytoskeleton (Profilin 2) |
| *Pole3* | ↓↓ | ↓↓ | DNA replication (DNA polymerase epsilon catalytic subunit 1) |
| *Pno1* | ↓ | ↑↑ | Ribosomal assembly (Partner of NOB1 homolog) |
| *Sdhd* | ↓↓ | ↓↓↓ | mitochondrial function (Succinate dehydrogenase) |
| *Snx14* | ↑ | ↓ | intracellular transport, autophagy (Sorting nexin 14) |
| *Stxbp6* | ↑ | ↑↑ | exocytosis, cell adhesion (Syntaxin-binding protein 6) |
| *Tmem213* | ↑↑ | ↑ | membrane transport (Transmembrane protein 213) |
| *Ubash3b* | ↑ | ↓ | cellular signaling (Ubiquitin-associated and SH3 domain-containing protein B) |
| *Uqcrh* | ↓↓ | ↓↓↓↓↓ | mitochondrial function (Ubiquinol-Cytochrome C reductase hinge protein) |

Arrow numbers corresponding to coef_mean: ↑: 0.005-0.01; ↑↑: 0.01-0.02; ↑↑↑: 0.02-0.03; ↑↑↑↑=: 0.03-0.04; ↑↑↑↑↑: >0.04; ↓: -0.005-0.001; ↓↓: -0.01-0.02; ↓↓↓: -0.02-0.03; ↓↓↓↓: -0.03-0.04; ↓↓↓↓↓: -0.04-0.05
