## Supplemental Table 4 for "Resolving thyroid lineage cell trajectories merging into a dual endocrine gland in mammals"

**Supplementary Table 4.** Identity and features of differentially expressed genes in immature and differentiated cells of the ultimobranchial lineage

*Calca*_pos vs *Calca*_neg Ubb cells

| Gene | Protein | Development | Endocrine | Function(s) |
|---|---|---|---|---|
| *Nnat* | Neuronatin | neural fate, brain | pituitary, pancreatic beta-cells | calcium regulation |
| *Mest* | Mesoderm-specific transcript homolog | mesoderm | adrenal | EMT |
| *Foxa2* | Forkhead box A | endoderm, lung, liver, pancreas | thyroid C cells, pancreas | pioneer transcription factor |
| *Btg2* | B-cell translocation gene 2 | neurogenesis | pancreatic beta-cells | cell cycle |
| *Clpx2* | Complexin 2 (synaphin) | - | neuroendocrine | exocytosis |
| *Ascl1* | Achaete-scute homolog 1 (MASH1) | neurogenesis | adrenal, thyroid C cells, MTC | pioneer transcription factor |
| *Cdc25b* | Cdc25 isoform | early embryogenesis | MTC | cell cycle |
| *Socs2* | Suppressor of cytokine signaling 2 | neuronal differentiation | pancreatic beta-cells | JAK/STAT signaling |
| *Hair1* | Hoxa adjacent long noncoding RNA 1 | ESC lineage differentiation | - | RA regulation |
| *Sst* | Somatostatin | neuroendocrine differentiation | neuroendocrine, MTC | peptide hormone |
| *Krt7* | Keratin 7 | epithelial differentiation | neuroendocrine | cytoskeleton |
| *Prox1* | Prospero homeobox 1 | cell fate determination | neuroendocrine, thyroid C cells, MTC | EMT, secretory pathway |

*Calca*_neg vs *Calca*_pos Ubb cells

| Gene | Protein | Function(s) |
|---|---|---|
| *Hmcn1* | Hemicentin 1 (HMCN1) | ECM, basement membrane organization, cell anchorage |
| *Plagl1* | PLAGL1 (ZAC-1) | transcription factor, ECM regulation, anti-proliferative |
| *Hs6st2* | Heparan sulfate 6 sulfotransferase 2 | ECM regulation, morphogen/growth factor activation |
| *Rbm12b2* | RNA binding motif protein 12 | post-transcriptional regulation |
| *Smtnl2* | Smoothelin-like 2 | apical actin turnover |
| *Dscc1* | DNA replication and sister chromatid cohesion 1 | DNA replication |

List of genes relates to scRNAseq data presented in Ordinary Fig. 7e (upper panel) and 7f (lower panel).
Based on scRNAseq analysis of E12.5 cells confined to Ubb clusters 4, 6 and 8
Abbreviations: Ubb ultimobranchial body, MTC medullary thyroid carcinoma, EMT epithelial-mesenchymal transition, ECM extracellular matrix
